# Particulate matter constituents trigger the formation of extracellular amyloid β and Tau-containing plaques and neurite shortening *in vitro*

**DOI:** 10.1101/2024.03.27.586796

**Authors:** Aleksandar Sebastijanović, Laura Maria Azzurra Camassa, Vilhelm Malmborg, Slavko Kralj, Joakim Pagels, Ulla Vogel, Shan Zienolddiny-Narui, Iztok Urbančič, Tilen Koklič, Janez Štrancar

## Abstract

**Introduction:** Air pollution is an environmental factor associated with Alzheimer’s disease, characterized by decreased cognitive abilities and memory. The limited models of sporadic Alzheimer’s disease fail to replicate all pathological hallmarks of the disease, making it challenging to uncover potential environmental causes. Environmentally driven models of Alzheimer’s disease are thus timely and necessary.

**Methods:** We used live-cell confocal fluorescent imaging combined with high-resolution stimulated emission depletion (STED) microscopy to follow the response of neuron-like cells to nanomaterial exposure. Here, we report that a high dose rate *in vitro* exposure of neuron-like cells to particulate matter constituents reproduces neurodegenerative phenotype, including extracellular amyloid-β containing plaques and decreased neurite length.

**Results:** Consistent with the existing *in vivo* research, we observed detrimental effects, specifically a substantial reduction in neurite length and formation of amyloid beta plaques, after exposure to iron oxide and diesel exhaust particles. Conversely, after exposure to engineered cerium oxide nanoparticles, the lengths of neurites were maintained, and almost no extracellular amyloid beta plaques were formed.

**Discussion:** Although the exact mechanism behind this effect remains to be explained, the high dose rate *in vitro* model, comprising wild-type neuron-like cells, could serve as an alternative environmentally driven model of Alzheimer’s disease.

Graphical abstract
High dose rate *in vitro* exposure of neuron-like cells to particulate matter constituents, like diesel exhaust and iron oxide nanoparticles, reproduces neurodegenerative phenotype, including extracellular amyloid-β-containing plaques and reduction in neurite length and density.

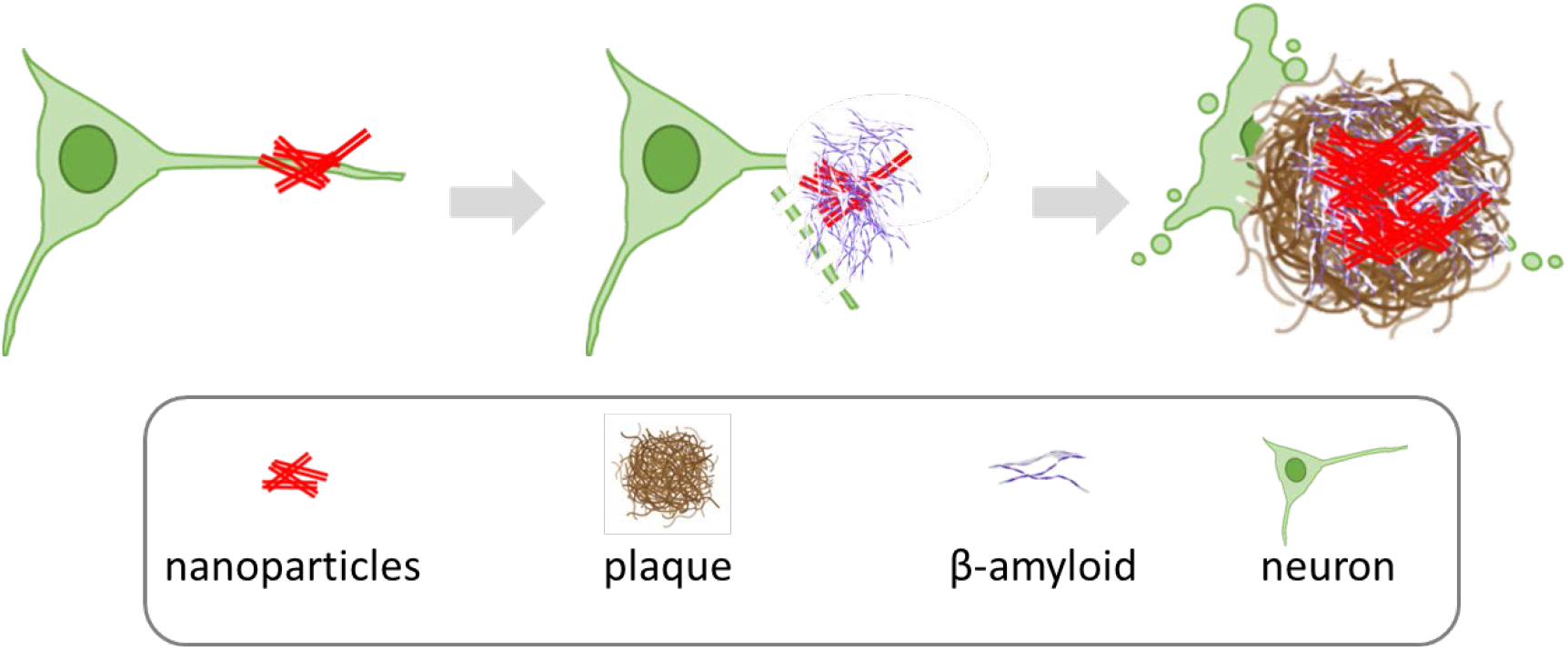

## 1 Introduction

### 1.1 Particulate matter in polluted air is associated with cognitive decline and Alzheimer’s disease, yet causality is still not fully established

Strong correlations between particulate air pollution and dementia suggest airborne particulate matter as a possible environmental trigger of neurodegenerative diseases (Peeples, 2020; Shi et al., 2020, 2021). However, whether the particulate matter in air pollution can cause dementia remains controversial (Underwood, 2017). It is well-established that high levels of air pollution, particularly particulate matter smaller than 2.5 μm (PM2.5), are associated with cognitive decline and Alzheimer’s disease (AD) (Carey et al., 2018; Younan et al., 2020; X. Zhang et al., 2018). However, it is important to note that the strength of this association does not necessarily imply causation. How air pollution particles enter the brain and possibly trigger neurodegeneration remains an open question (Underwood, 2017). Nevertheless, a recent study has observed epigenetic gene regulation changes that are typically found in Alzheimer’s patients in both young adults and mice exposed to particulate air pollution (Calderón-Garcidueñas et al., 2020). In another study, authors indicate that there may be no safe exposure limit to outdoor fine particulate air pollution (PM_2.5_) in relation to the risk of dementia (*How Low Can You Go? Air Pollution Affects Mortality at Very Low Levels | Science Advances*, n.d.). Considering the emerging evidence, the Lancet Commission recognized air pollution as a risk factor for dementia, suggesting that reducing air pollution could potentially prevent or delay the onset of dementia, which is vital because pre-symptomatic diagnosis of disease is still lacking (Livingston et al., 2020).

AD, in particular, is a most prevalent form of dementia characterized by early memory loss and progressive damage to brain cells in the hippocampus, a region responsible for learning and memory (Drew, 2018). Despite more than half a century of research, the exact causes of disease remain uncertain, and no effective treatment yet exists (Drew, 2018), despite recently FDA-approved therapies (Couzin-Frankel, 2023; Piller, 2023). Neurodegenerative diseases, such as AD, Parkinson’s, amyotrophic lateral sclerosis, and Huntington’s disease, are characterized by the misfolding and aggregation of specific proteins into large plaques (Eftekharzadeh et al., 2016). Epidemiological data suggest that nanoparticles accumulated in the brain might trigger the formation of AD-like plaques. Furthermore, imaging of thin sections of amyloid cores from AD patients has corroborated the presence of ambient magnetite nanoparticles in the plaques (Plascencia-Villa et al., 2016). Although, most studies linking ambient air pollution with neurodegenerative disease have considered PM_2.5_ as exposure metric, ultrafine particles (PM_0.1_) or nanoparticles (NP) are more even numerous, cause more inflammation, and can translocate to essentially all organs (Schraufnagel, 2020). It has been suggested that combustion emissions such as diesel exhaust particles and engineered nanoparticles are of special importance in neurogenerative disease (Calderón-Garcidueñas & Ayala, 2022). Existing research thus shows the need for new methods to confirm the causal connection between the components of particulate matter and neurodegeneration.

In this paper, we investigated whether exposure of neuron-like cells to particulate matter constituents could replicate neurodegenerative phenotypes *in vitro*. These include formation of extracellular amyloid-β-containing plaques and reductions in both neurite density and length. Employing a high dose rate *in vitro* exposure of wild-type neuron-like cells allowed us to emulate a dose accumulated over a lifetime, delivering it within approximately a day, depending on the sedimentation rate of the nanomaterial. Previously, we employed a similar high dose rate *in vitro* exposure technique in a coculture of lung epithelial cells and alveolar macrophages, successfully replicating phenotypic characteristics observed in long-term lung inflammation occurring 90 days after exposure in mice (Danielsen et al., 2020; Kokot et al., 2020).

We tested four types of nanomaterials: two engineered nanomaterials and two nanomaterials commonly found in polluted air:

1. **anatase TiO_2_ nanotubes** are an engineered nanomaterial relevant predominantly to occupational risk. In general, TiO_2_ particulate matter, also known as E171, is widely used as a white colorant in food, paints, coatings, pharmaceuticals, cosmetics, and even in toothpaste. E171 contains a fraction of nanosized primary particles (<100 nm) (Peters et al., 2014, 2020). Several *in vivo* studies reviewed by Song et al. (Song et al., 2015) have demonstrated that the TiO_2_ NP can be transported and accumulated in the brain, but mostly in intranasal instillation or inhalation studies, eventually leading to central nervous system dysfunctions. Another group showed that titanium content in mouse hippocampus after 90 days of continuous intranasal exposure to TiO_2_ nanoparticles was 0.6 mg/g tissue at the highest administered dose of 10 mg/kg body weight (Ze et al., 2014), much higher than the levels observed in humans. Titanium can be found in human tissue at lower average concentrations ranging from 1 μg/g (Peters et al., 2020) up to 80 μg/g (Baj et al., 2023) in the human brain. Thus, neuroinflammation and impairment of spatial memory observed in mice occur at more than ten times higher brain content than observed in humans. The effect of TiO_2_ NP on the brain is thus still controversial since detrimental effects are observed only at very high doses. Here, we also use a very high concentration of 3 mg/g, akin to *in vivo* dosages, to show that inflammatory, engineered TiO_2_ nanotubes induce the formation of extracellular amyloid β-containing plaques and cause shortening of neurites. Yet, they do not cause such extensive neurite shortening as iron oxide and diesel exhaust particles.
2. **Iron oxide (γ-Fe_2_O_3_) nanoparticles** are commonly found in road traffic emissions from brake wear (Maher, 2019). The PM_2.5_ concentrations of iron oxide particles is high in underground railway stations. Sheikh et al. (Sheikh et al., 2022) showed that the majority of iron oxide particulate matter exposure in the London Underground was maghemite, the material used in our study. Occupational exposure to welding fume particles (iron-rich particles and magnetite) is associated with an increased risk of developing AD. Finnegan et al. showed that non-chemically preserved AD patients’ post-mortem tissue samples contained the average concentration of iron peaking at 400 mg/g (Finnegan et al., 2019). Furthermore, airborne iron oxide (magnetite) particles can be concentrated in smaller regions of the human brain (Maher et al., 2016), thus reaching even higher local concentrations. Such magnetite nanospheres are ubiquitous and abundant in airborne particulate matter pollution. They are smaller than 200 nm in diameter and can enter the brain directly via the olfactory bulb (Maher et al., 2016). Examination of the human frontal cortex brain samples obtained from subjects who lived in Mexico City revealed that magnetite nanoparticles have an external, rather than an endogenous source (Maher et al., 2016). Although the highest brain magnetite concentration was 10 µg/g of dry tissue, the transmission electron micrographs of brain thin sections showed that the nanoparticles are in about 1% of the total area; thus *in vitro* concentrations of 1 mg/g might be more relevant to human exposure to polluted air.
3. Since iron is present in most **diesel exhaust** samples, representing on average about 20 % w/w (Viskup et al., 2020), we estimate that the brain burden of diesel exhaust particles might reach up to 2 mg/g tissue weight, coinciding with the dosage used in this manuscript. Although iron might not be a good exposure metric for diesel exhaust, we also considered an exposure concentration of 100 μg/m^3^ as elemental carbon (Debia et al., 2017), to estimate the exposure over an entire working life (Supplement chapter 7 Dose relevance). The specific surface area of diesel exhaust particles is similar to particles generated by welding, with particle sizes mostly below 200 nm, so the concentration of insoluble surface area that might accumulate in the brain may be in the same range as occupational exposure of welding workers, given that the transport mechanisms across barriers are similar for the two types of aerosols.
4. As an example of reportedly nontoxic particulate matter, we tested **CeO_2_ nanoparticles**, which are used as fuel additive in motor vehicles to reduce carbon monoxide, nitrogen oxides, and hydrocarbons in exhaust gases. Thus, the particles are released into the air. Effects of CeO_2_ nanoparticles reported in the literature are contradictory, demonstrating all kinds of effects, from protective to toxic (Gagnon & Fromm, 2015). For example, CeO_2_ nanospheres and nanorods improved cognitive impairment following mild traumatic brain injury in VC57BL/6J male mice at the dose of 0.02 mg/g body weight. However, the concentration in the brain was not reported (Fiorani et al., 2015; Gagnon & Fromm, 2015).

### 1.2 The challenge of reproducing all hallmarks of Alzheimer’s disease (AD) in animal models: the limitations of genetically modified models and the need for an environmentally triggered wild-type model

AD is defined by a triad of pathological hallmarks (1) amyloid β peptide plaques, 2) aggregation of Tau protein, and 3) neuronal degeneration. Accumulation of amyloid β plaques precedes cognitive impairment by at least 20 years (Scheltens et al., 2016). Extracellular deposits of amyloid β have long been considered the initial event of the disease process (Leng & Edison, 2021), at least in less prevalent familial cases (Karran et al., 2011). The accumulation of Tau into large aggregates, initially appearing in dystrophic neurites surrounding Aβ plaques, is usually referred to as neuritic plaque tau (NP tau, followed by the formation and spread of neurofibrillary tangles (NFT) and neuropil threads to other neurons. Recently He et al. presented a new mouse model, which offered a new way to explain how the amyloid β plaque environment accelerates the spread of Tau pathology in AD patients’ brains, consistent with imaging studies and investigations of post-mortem AD brains (He et al., 2018). Axonal and dendritic degeneration and neuronal death, reflected in reduced neuronal arborization and decreased neuron density, is the final pathological hallmark that coincides with the appearance of intracellular Tau deposits and symptoms of advanced cognitive decline (Falke et al., 2003).

Here, we show that high dose rate *in vitro* exposure of wild-type neuron-like cells to particulate matter constituents, such as diesel exhaust and iron oxide nanoparticles, replicates hallmarks of AD pathology by triggering amyloid β plaque formation and reducing the density and length of neurites. Since less than 0.1% (Blennow et al., 2006) of AD is caused by a single genetic mutation, the model system consisting of wild-type neurons without familial mutations might thus be regarded as an alternative, perhaps more relevant, environmentally driven AD model system.

## 2 Materials and methods

### 2.1 Cell line

Human neuroblastoma cell line commercially available from the American Type Culture Collection (ATCC, www.atcc.org or www.lgcstandardsatcc.org, catalogue number CRL-2266™). grew as a mixture of floating and adherent cells in DMEM/F-12 (Gibco) medium supplemented with 1% penicillin/streptomycin (Sigma-Aldrich), 2 mM GlutaMax (Gibco), and 10 % FBS (BioChrom AG). For SH-SY5Y differentiation, we seeded 1 x 10^6^ cells cm^-2^ on the collagen-coated, microscopy-adapted plates (8-wells, Ibidi). After 24 hours we added the differentiation medium containing Dulbecco’s modified eagle medium DMEM/F12 without phenol red (#21041025, Thermo Fisher), supplemented with 1% penicillin/streptomycin (Sigma-Aldrich), 2 mM GlutaMax (Gibco), 0,5 % FBS (BioChrom AG) and 10 uM retinoic acid (fisher scientific, #AC207341000). Old media was replaced with 100 µL fresh differentiation media every other day, until 7^th^ day when we terminated the differentiation.

### 2.2 Nanomaterial preparation

We employed a dispersion technique using cup horn sonication to disperse materials, namely anatase TiO_2_, maghemite γ-Fe_2_O_3_, CeO_2_, and diesel exhaust particles, in a buffer. The buffer solution utilized in this study was 1 mM bicarbonate buffer with a pH of 10, which was specifically chosen due to its low osmolarity and high pH properties that minimize the charge screening of the active surface of the materials.

To ensure uniform dispersion of the materials in the bicarbonate buffer, the materials were resuspended so that each 3 µL of the suspension contained 3 cm^2^ of the material surface. The weighted mass of each material and the volumes of bicarbonate buffer utilized to prepare the solutions are presented in the table below.

The synthesis of TiO_2_ nanotubes followed a rigorous procedure and has been extensively documented (Umek et al., 2007). In brief, the process began with the controlled hydrothermal synthesis of sodium titanate nanotubes. These sodium titanate nanotubes were subsequently subjected to ion exchange to transform them into hydrogen titanate nanotubes. Finally, through a carefully conducted thermal treatment, the hydrogen titanate nanotubes were successfully converted into the desired TiO_2_ nanotubes. The detailed methodology and experimental conditions can be found in the comprehensive study by Umek et al. (Umek et al., 2007).

The iron oxide nanoparticles in the form of maghemite (γ-Fe_2_O_3_) were synthesized by co-precipitation from aqueous solutions of Fe^2+^ and Fe^3+^ salts. The synthesis methodology is described in detail in our previous publications (Kralj et al., 2012; Kralj & Makovec, 2014; Nemec et al., 2020). The saturation magnetization of as-synthesized nanoparticles is ∼64 emu g^-1^. The size of nanoparticles was 11.6 ± 2.1 nm which was estimated by analysing 100 transmission electron microscopy images using ImageJ software. For the preparation of stable colloidal suspension of superparamagnetic nanoparticles its surface was electrostatically stabilized using citric acid as described elsewhere (Nemec & Kralj, 2021). The final concentration of citric-acid stabilized iron oxide maghemite nanoparticles in aqueous suspension was 1.8 mg ml^-1^.

DEP9.7 diesel exhaust particles were collected after-treatment from an experimental modern heavy-duty diesel engine under highly controlled combustion conditions in a laboratory environment. The particle production and collection has been described in detail previously (Gren et al., 2020). The engine was operated to control the physical and chemical properties of the diesel exhaust particles (DEP9.7). To produce the DEP9.7 particles, the engine was run at an engine intake O_2_ concentration of 9.7% using petroleum-based ultralow-sulfur diesel fuel of Swedish MK1 standard. The diesel engine combustion was modified to this low-temperature combustion condition using a high amount of exhaust gas recirculation. The exhaust gas recirculation lowers flame temperatures and reduces both soot formation and soot oxidation processes. Using the high control of the engine, the DEP9.7 particles were intentionally designed to have a high content of polycyclic aromatic hydrocarbons (PAH) and refractory organic carbon relative to the content of elemental carbon. DEP9.7 has been characterized extensively(Bendtsen et al., 2020, p. 6; Gren et al., 2020). DEP were collected on a Teflon filter and extracted using methanol with >80% recovery. The DEP9.7 had a primary particle size of 22 nm and an estimated specific surface area of 152 m^2^g^-1^. The carbon content in DEP9.7 is 33% elemental carbon and 67% organic carbon of which 33% is refractory organic carbon. DEP9.7 contains a number of metals (Bendtsen et al., 2020, p. 6) predominately Cu (2349 ug g^-1^), Fe (220 ug g^-1^), Sr (99 ug g^-1^) and Mn (92 ug g^-1^). DEP9.7 has a high content of PAH (26.8 mg g^-1^) and a list of specific PAH can be found in (Bendtsen et al., 2020, p. 6). In comparison to four other sample types from the same engine, DEP9.7 had the lowest formation of Reactive Oxygen Species in the cell free DCFH assay on mass basis. Extractable organic matter from DEP9.7 had the highest genotoxicity potential in lung epithelial (A549) cells of the five samples (Rothmann et al., 2023).

CeO_2_ NPs "50nm agglomerates" composed of 5 nm CeO2 nanoparticles as determined from TEM images (batch AN-123) were produced and characterized by Applied NanoParticles - A Nanotech Engineering Company (Spain) (https://www.appliednanoparticles.eu/). CeO_2_ NPs have a mean diameter of 6.2 +-1.3 nm as measured by TEM, the agglomerates have a mean diameter of 200 +-6 nm as measured by DLS and a zeta potential −32.5 +- 0.07 mV measured in miliQ water at pH 9 at 0.5mS cm^-1^.

The resulting suspensions of the nanomaterials were then subjected to cup horn sonication in an ice bath for 15 minutes, utilizing 5 seconds on - 5 seconds off regime for a total duration of 30 minutes, at a power setting of 20-30 W (amplitude 70) to guarantee optimal dispersion.

The volume of material dispersion containing 10 times larger surface than that of neurons was added in a dropwise manner, prior to microscopy, to cover the surface of the *in vitro* model. The volume applied to cells never exceeded 10% of the media volume.

### 2.3 Fluorescent labelling

Different fluorescent labels used in this study are given in Table 2, below.

**Table 1:** Preparation of nanomaterial dispersions used in this study.

| Material | Mass (mg) | Volume (mL) | BET ($\text{m}^2 \text{g}^{-1}$) | Final concentration ($\text{mg mL}^{-1}$) |
| --- | --- | --- | --- | --- |
| $\text{TiO}_2$ nanotubes | 1.0 | 1.6 | 150 | 0.65 |
| $\gamma\text{-Fe}_2\text{O}_3$ | 1.80 | 1.0 | 109 | 1.80 |
| $\text{CeO}_2$ | 1.6 | 1.0 | 65 | 1.60 |
| Diesel exhaust particles (DEP9.7) | 1.0 | 2.0 | 152 | 0.50 |

**Table 2.** List of fluorescent labels used in this study.

| Fluorophore | Manufacturer (cat. nr) | Excitation peak (nm) | Emission peak (nm) | Concentration ($\mu\text{M}$) |
| --- | --- | --- | --- | --- |
| CellTracker™ Green CMFDA (CTG) | Thermo Fisher (#C2925) | 492 | 517 | 1 |
| ATTO490 LS conjugated to anti-TAU ab | ATTO-TEC | 495 | 658 | 0.01 |
| CellMask™ Deep Red (CMDR) | Thermo Fisher (#C10046) | 650 | 685 | 1 |
| Alexa-647 conjugated to anti-Aβ ab | Thermo Fisher (#A20006) | 650 | 665 | 0.01 |

In our experimental procedures, a specific set of labels was utilized for each experiment, which will be detailed in the Materials and Methods section and the corresponding figure legends. In general, live neurons were differentiated and incubated with nanomaterial, followed by staining with fluorescent labels according to the manufacturers’ guidelines.

It is noteworthy that CTG was the only label added one day prior to microscopy, while other labels were added immediately before imaging. Additionally, only CTG was washed with PBS, whereas other labels were not washed. Antibodies used in this study will be described later in this section. Briefly, they were conjugated with fluorescent dyes and were added in specified concentrations just prior to microscopy.

Anti- Aβ antibodies are raised against amino acids 1-40 of β-Amyloid of human origin (https://www.scbt.com/p/beta-amyloid-antibody-20-1). The β-Amyloid Antibody (20.1) is a mouse monoclonal antibody of the IgG2b κ isotype specifically designed to target β-amyloid and amyloid precursor protein (APP) fragment from human sources.

The extracellular tau deposits were labelled with Tau (Tau-5) Mouse Monoclonal Antibody (<u>Product # AHB0042</u>) of IgG1 kappa κ isotype. It recognizes both phosphorylated and non-phosphorylated human tau proteins.

Both antibodies were added at 2 μg mL^-1^ concentration to 7-day RA differentiated neurons in 1% BSA at 37°C. Imaging started immediately after the addition of antibodies.

### 2.4 Microscope setup

Following incubation with nanomaterial, the living samples were labelled and imaged using a state-of-the-art imaging system consisting of an Olympus (Olympus IX83) inverted microscope body equipped with a laser-scanning confocal and STED unit (Stedycon, Abberior). Stage top incubator (Okolab H301-MIN) maintains atmosphere with the 37°C, 5% CO_2_, and >95% humidity to enable long term imaging of living cells.

The images were captured using a confocal microscope equipped with either a 20x magnification objective and 0.8 numerical aperture (NA) lens or a 60x magnification objective with a 1.2 NA lens. The microscope system incorporates four pulsed laser sources (Abberior) with a pulse duration of 120 ps and a maximum power of 50 µW at the sample plane. Additionally, four avalanche photodiode (APD) detectors are utilized for signal detection. The pulse repetition frequency was 80 MHz. The STED depletion laser, operating at a wavelength of 775 nm, had the same repetition frequency as the excitation lasers, a pulse duration of 1.2 ns, and a maximum power of 170 mW at the sample plane.

We detected nanoparticles in the label-free, backscatter detection mode, utilizing the 488 / 488 ± 5 nm excitation / detection. Due to the large coherence of the laser, the backscattered light exhibited a strong speckle pattern, which was removed by Fourier transform bandpass filter (1 – 100 pixels) on scattering images. These images were subsequently binarized to obtain nanomaterial masks.

Please see the supplement for information regarding nanomaterial aggregation in cell medium, and automated-multi region of interest (ROI) data analysis and quantification of extracellular amyloid β containing plaques and neurite lengths.

## 3 Results and discussion

### 3.1 TiO_2_ nanotubes trigger the formation of extracellular amyloid β, Tau, and neuronal cytosolic components containing plaques

Since anatase TiO_2_ nanotubes induce retention of organic debris in alveolae with nanoparticles at the core of these deposits (Danielsen et al., 2020; Kokot et al., 2020), we tested whether the formation of similar deposits of organic molecules might occur inside the brain. We employed a human neuroblastoma cell line called SH-SY5Y, which is extensively utilized in studies due to their ability to differentiate into neuron-like cells. When differentiated, these cells exhibit characteristics resembling dopaminergic neurons found in the ventral hippocampus, which are involved in cognitive functions (Titulaer et al., 2021).

After differentiation of SH-SY5Y, resulting in neuron-like cells with extensively branched neurites (Figure 1 A green) (Agholme et al., 2010; Kovalevich et al., 2021; Kovalevich & Langford, 2013), we exposed them to TiO_2_ nanotubes (Figure 1 A red). Most of the added TiO_2_ nanoparticles aggregate (Supplement Figure S 1 to 18) and settle to the glass surface never interacting with the cells (purely red signal in Figure 1 A), whereas some portion of added nanoparticles, mostly single nanotubes (Urbančič et al., 2018), translocate into the cells and gets excreted after a day (Kokot et al., 2020). We identified the nanoparticles that interact with cells by their co-localisation with the cytosolic label CellTracker™ Green CMFDA (CTG) (green, Figure 1 B-D), with which we labeled the neurons prior to exposure. This label, once internalized by the cells, is acted upon by cellular esterases rendering it fluorescent throughout the cytoplasm. Without this interaction, it remains non fluorescent. In addition, thiol reactivity increases its intracellular retention through several generations (Beem & Segal, 2013), thus precluding it from freely exiting the cells to adhere to the surface of TiO_2_ nanotubes that had not interacted with the cells’ interior.

**Figure 1.**
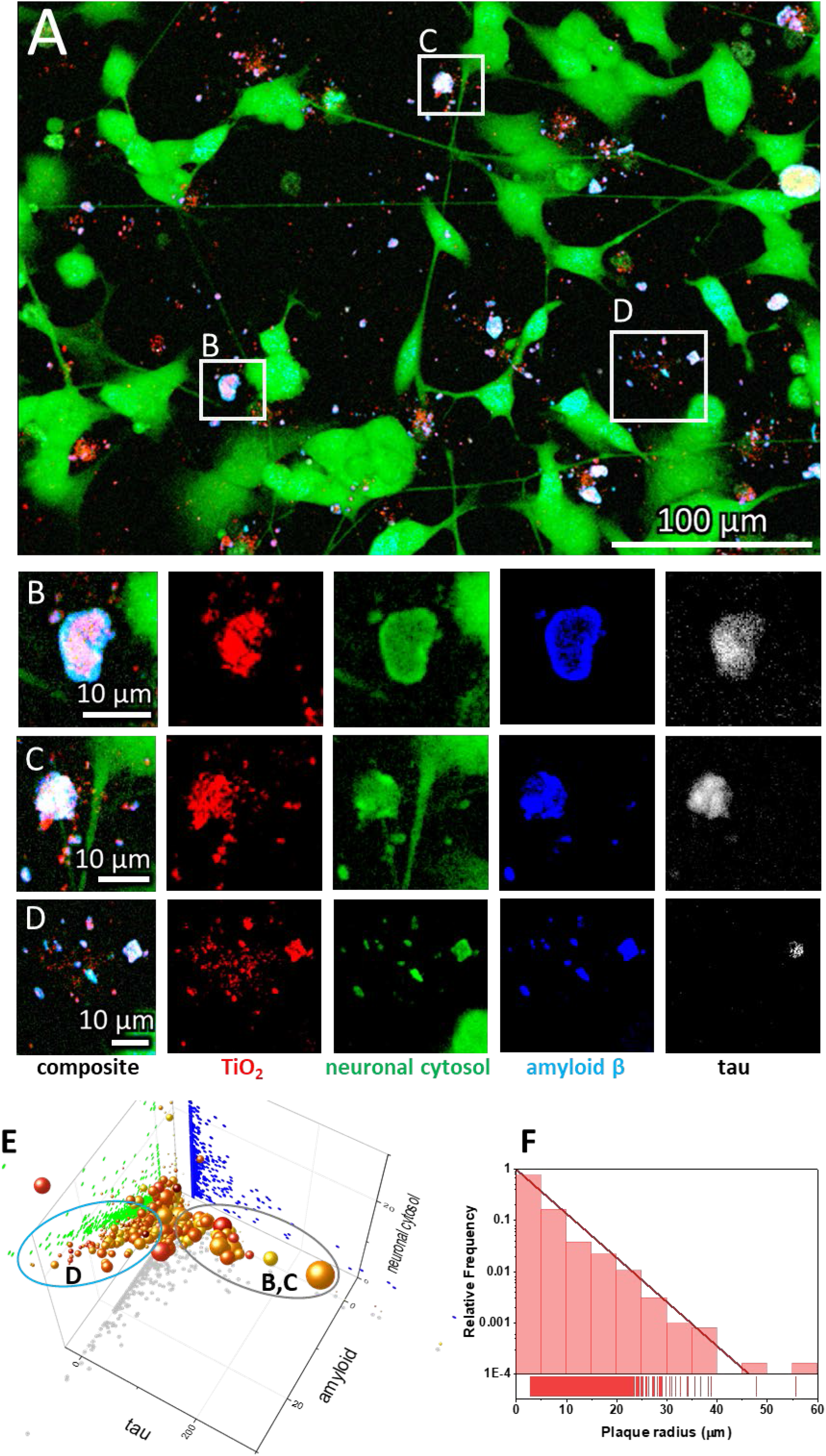
SH-SY5Y-derived neurons exposed to anatase TiO_2_ nanotubes form amyloid-β and Tau protein-containing extracellular plaques. A) Representative image of differentiated SH-SY5Y cells in culture; for differentiation, cells were grown for seven days in DMEM/F12, 10% fetal bovine serum, and 10 µM retinoic acid, followed by seven days in serum-free DMEM/F12 and 10 ng/mL brain-derived neurotrophic factor. Note the altered morphology and pronounced neurite outgrowth upon differentiation. Live cell culture was stained with cytosolic fluorophore CellTracker™ Green (green), which is retained and fluoresces only within a cell, a mouse monoclonal antibody raised against amino acids 1-40 of Aβ of human origin, for detection of amyloid precursor protein and Aβ (blue), and Tau-5 mouse monoclonal antibody against total Tau (white). Within the large field of view, the rectangles denote regions enlarged in the panels (B-D) showing examples of heterogeneous extracellular plaques at higher magnification: B-C) large plaques with low Aβ surface density (fluorescence intensity signal divided by the area of the signal) and high tau surface density; D) small plaques with high Aβ surface density and low tau surface density; E) Properties of all the Aβ plaques from 15 regions of interest, each covering 120 x 120 μm^2^ in terms of surface densities with x-axis: tau antibody, y-axis: Aβ antibody, z-axis: neuronal cytosolic label, size of a sphere: plaque radius, and color: density of nanoparticles in a plaque, each sphere representing one plaque. Most of the nanomaterial is deposited on the glass surface without interacting with neurons (red), whereas the nanomaterial interacting with the neurons contains Aβ (blue), nanomaterial (red), and neuronal cytoplasmic components (green), while Tau (white) can be missing; The grey and blue ellipses denote plaques with high and low density of Tau, such as those in panels B/C and D, respectively. F) The size distribution of the plaques, showing exponential decay of the number of plaques versus plaque size.

Moreover, we observed that many of the above-mentioned aggregates also contained extracellular deposits of amyloid β (Aβ) peptide, as indicated by immunostaining with a mouse monoclonal antibody raised against amino acids 1-40 of Aβ of human origin, for detection of amyloid precursor protein and Aβ (Figure 1 B-D blue). Namely, aggregation of Aβ is a proteolytic product of amyloid precursor protein (AβPP) by β- and γ-secretases, resulting in two major Aβ isoforms: Aβ42 (42 residues long) and Aβ40 (40 residues long). Although the concentration of Aβ40 in cerebral spinal fluid is several-fold higher than that of Aβ42, the Aβ42 is the predominant constituent of amyloid β plaques in AD brains (Iwatsubo et al., 1994, 1995), while Aβ40 is present in only some of the plaques (Gravina et al., 1995; Mak et al., 1994), suggesting that initial plaque formation involves only Aβ42 (Gu & Guo, 2013). Therefore, we are aware of a limitation that quantifying only Aβ40 might not be suitable for detection of early Aβ containing plaques *in vitro*.

Phosphorylated Tau (p-tau) is also one of the biomarkers for early stages of AD. For example, Tau phosphorylated at threonine-231 (T231) increase early in the preclinical stage of AD, probably in response to Aβ pathology (Suárez-Calvet et al., 2020). Therefore, we labeled the extracellular tau deposits with tau (Tau-5) mouse monoclonal antibody that should recognize both phosphorylated and non-phosphorylated human tau proteins (Figure 1 B-D white). Since cells can release Tau even if not exposed to a toxicant (Karch et al., 2012), we cannot deduce whether nanoparticles bind Tau within a cell and transport it out or the cells release more Tau, which binds to nanoparticles outside of the cells. Tau is a neuronal microtubule-associated protein found predominantly in axons. The C-terminus binds axonal microtubules while the N-terminus binds neural plasma membrane components (Brandt et al., 1995; Padmanabhan et al., 2022). Many other roles for Tau were speculated (Sotiropoulos et al., 2017), including its role in insulin signaling (Marciniak et al., 2017) and being released extracellularly as a free protein (De La-Rocque et al., 2021), which can bind to receptors on living cells (Mudher et al., 2017), including amyloid precursor protein (Takahashi et al., 2015). Yet the precise mechanisms of tau pathology remain unanswered (H. Zhang et al., 2021).

We observed the formation of extracellular deposits of Aβ and Tau (Figure 1 composite, white deposits near green neurons), which agrees with the current knowledge of extracellular plaque composition found in AD patients. There, two types of brain lesions are commonly observed: Aβ plaques and tau tangles, arising from the aggregation of misfolded Aβ and hyperphosphorylated tau proteins, respectively (Walker, 2020). Aβ plaques have long been considered the initial event of the disease process (Leng & Edison, 2021). However, recent studies have shown that Aβ plaques can act as seeds for the initial Tau accumulation into large aggregates containing both proteins. This initial accumulation occurs in dystrophic neurites surrounding Aβ plaques, referred to as neuritic plaque tau (NP Tau), consistent with imaging studies and investigations of post-mortem AD brains (He et al., 2018). Our data (Figure 1 B-D) corroborates this notion, showing protein deposits containing both Aβ and tau protein, resembling structures observed in mice and patients by He et al. (He et al., 2018).

To examine the heterogeneity of extracellular Aβ plaques, we analyzed 15 regions of interest (ROI), each covering 120 x 120 μm^2^. Example of the algorithm we used for calculating densities of plaque components is shown in the supplement (Figure S 21). We identified more than six thousand plaques, the properties of which are plotted in a 5-dimensional graph in Figure 1 E. As x coordinate, we assigned the Aβ signal density (fluorescence signal intensity divided by the surface area of an extracellular deposit = plaque), as y coordinate, we assigned the Tau signal density, and as z coordinate, we assigned the cytoplasmic signal density. In this way one can see that some of the points lie in the xz plane (y=0, no tau signal), yet there are no points in xy (z=0) and yz (x=0) plane indicating that all extracellular deposits contain Aβ (blue) and cytoplasm of neurons (green). Each sphere in the graph represents one plaque, with the size of the sphere proportional to the plaque radius and the colour representing the density of the nanomaterial according to the colour scale. Interestingly, some plaques lack Tau (x axis), whereas others contain lots of it. This observation aligns with the heterogeneous nature of plaques found in AD patients, where there is high diversity in all plaque components except for Aβ, which is universally present in all plaques (Walker, 2020).

Notably, the amount of Aβ in our plaques is proportional to the amount of cytosolic label observed, indicating that the neurons generate Aβ because fluorescing cell tracker label must have originated from neuronal cytoplasm (Figure 1 E distribution of green dots in an amyloid-neuronal cytosol plane). This is in accordance with the finding that plaque formation initiates within the neuron, and subsequential accumulation of fibrillar Aβ occurs intraluminally (Le Bras, 2022; Lee et al., 2022; Pensalfini et al., 2014). Furthermore, many other cell proteins have been identified within Aβ plaques. For example, a comparison of biomolecules in Aβ plaques (amyloidome) of human Aβ plaque regions versus non-plaque regions of AD and healthy controls revealed significantly higher levels of neuronal proteins (Bai et al., 2021; Drummond et al., 2017; Xiong et al., 2019), further strengthening the idea that the plaque initially forms within the neuron.

The number of the plaques decreases exponentially with the radius of a plaque (Figure 1 F), with the characteristic length of 3.3 μm and most plaques having a radius below 40 μm, corresponding to cross-section area of 5000 μm^2^. Such a size distribution, including exponential behavior and maximum size, is similar to the distribution of Aβ oligomeric structures observed *in vitro* on lipid membranes by Tahirbegi et al. (Tahirbegi et al., 2020). However, the plaque sizes we observed are larger than suggested by Serrano-Pozo et al., who showed that in AD, patients maximum plaque sizes reach up to 500 μm^2^ (Serrano-Pozo et al., 2012). We speculate this difference might be due to the absence of microglia in our cell culture model, as the microglia are known to influence plaque formation(Bai et al., 2021; Leng & Edison, 2021). For example, Shabestari et al. showed that re-engraftment the population of microglia in AD mice brains resulted in formation of more compact, smaller plaques (Kiani Shabestari et al., 2022). Nevertheless, the size of plaques seems to be loosely governed by the density of Tau and Aβ proteins within them. Larger plaques tend to have a higher density of Tau, and smaller ones have higher Aβ density (Figure 1 D).

### 3.2 Amyloid β (Aβ) plaques form at the sites of damaged neurites

Above we have shown that exposure to nanoparticles can trigger the formation of Aβ plaques, possibly due to the direct damage to neurons, as evidenced by the presence of Tau and neuronal cytosolic components within the plaques. This is not that surprising considering our previous studies where we showed that these nanotubes can disturb plasma membrane (Urbančič et al., 2018) and cytosolic organelles (Kokot et al., 2020), including microtubule structure, within minutes after exposure. We next conducted high temporal and spatial resolution imaging of neurite morphology at different confocal planes (Figure 2 A, B) to investigate whether nanotubes can directly damage the neurites they contact. Next to an exemplary Aβ plaque that spans a height of 3 μm, we observed an axon extending from the neuron body at the lowest z plane (Figure 2 B, z = 0 μm) that stopped at the AB plaque. Interestingly, another axon above the first remained intact and reached over the plaque three μm above the lowest point (Figure 2 B, z = 3 μm).

**Figure 2.**
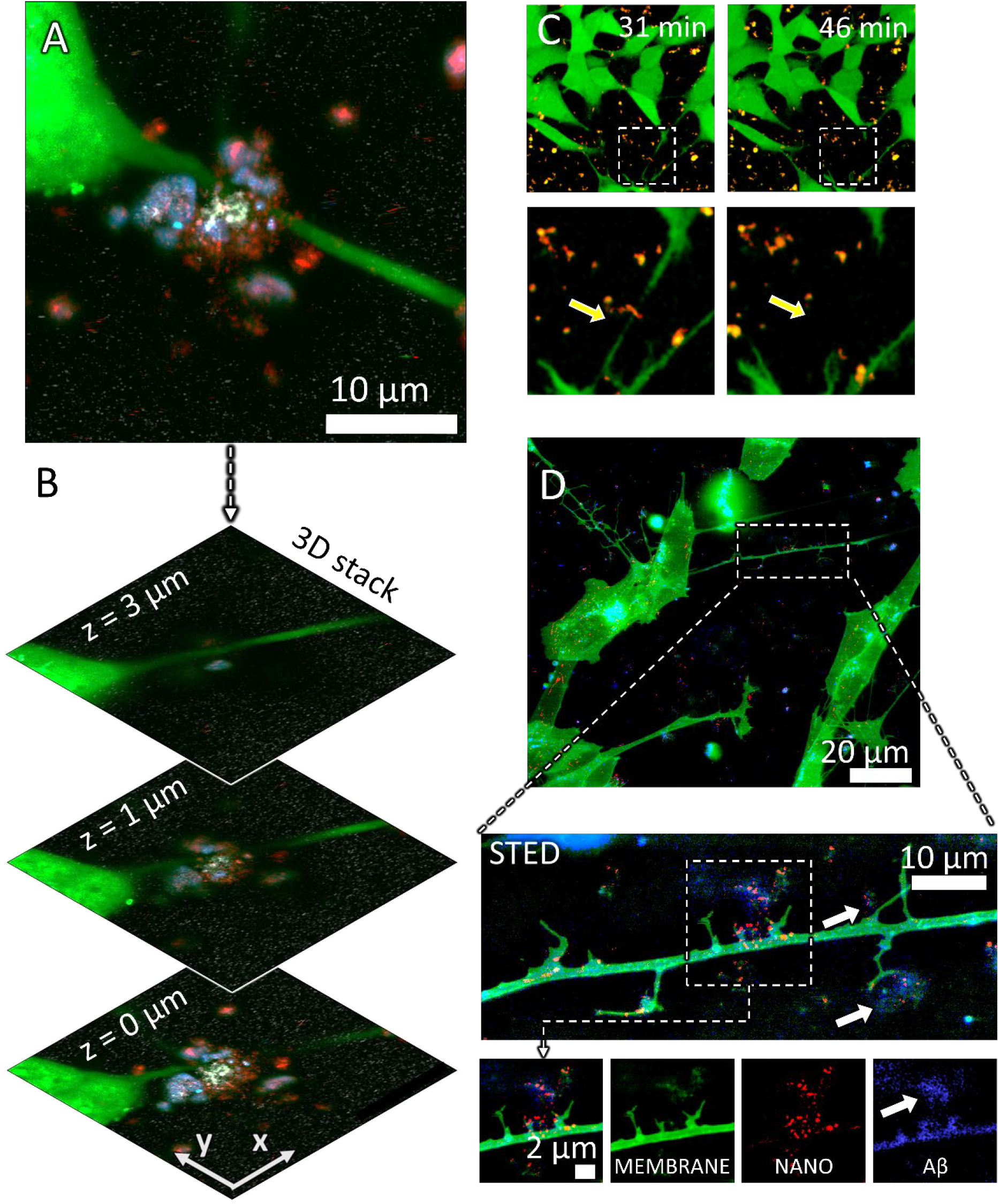
Neurite dystrophy and early amyloid β plaque formation at the sites of retracted neurites caused by anatase TiO_2_ nanotubes. A) A maximum intensity projection image of the confocal planes in B) Composite confocal fluorescence image of an Aβ containing plaque at three different z heights from the signal of probes for neuronal cytosol (green), TiO_2_ nanotubes (red), Aβ (blue), and Tau (white); C) live cell imaging of neurite collapse occurring within minutes after nanoparticle exposure, arrow points to a neurite that collapses in contact with the nanotubes; D) Neuronal plasma membrane labeled with CellMask™ Deep Red (CMDR) (green), High spatial resolution fluorescence image (stimulated emission depletion STED) of synapse damage and early plaque formation (white arrows) at the sites of damaged synapses (green, MEMBRANE) by the nanotubes (red, NANO) containing Aβ (blue, Aβ).

To gain a better understanding of these observations, we employed long-term time-lapse monitoring of neurites (Figure 2 C, green) to discover that TiO_2_ nanotube aggregates (red) quickly colocalized with components of neuronal cytosol get (yellow, overlap of red and green), leading to discontinuation of a neurite within minutes after exposure as indicated by the yellow arrows in Figure 2 C. Furthermore, high-resolution imaging using stimulated emission depletion (STED) microscopy revealed damaged synapses (green) where nanotubes (red) concentration was the highest (Figure 2 D white arrows). We also observed an early formation of a plaque near the neurite containing nanoparticles (red, Figure 2 D, STED zoom-in NANO), neuronal plasma membrane components (green, MEMBRANE), and Aβ (blue, Aβ). Conversely, in the absence of TiO_2_ nanotubes, neurites of that same axon retained their integrity and no Aβ was detected in their proximity.

3.3 Diesel exhaust and iron oxide nanoparticles induce the most neurite shortening Despite a strong epidemiological correlation between exposure to airborne particulate matter and the development of neurodegenerative diseases (Peeples, 2020; Shi et al., 2020, 2021), the causal relationship between particulate matter in air pollution and Aβ plaque deposition with neurite degeneration in AD remains controversial (Budson, 2020; Underwood, 2017). To address this, we investigated whether various types of particulate matter commonly found in polluted air, including metal oxides and carbonaceous particles, could trigger the formation of Aβ plaques and neurite shortening as observed above for the TiO_2_ nanotubes.

For this purpose, we compared the effect of TiO_2_ nanotubes, CeO_2_, γ-Fe_2_O_3_, and diesel exhaust particles on neurite lengths and their number in neuronal cell culture (Figure 3). We analyzed neurite lengths at 1 and 100 hours after exposure to these materials using the Fiji ImageJ with NeuronJ plug-in, a program that can be used for semi-automated tracing of individual neurons (Popko et al., 2009). Representative images of chosen neurites, each connecting two distinct cells, are shown in violet in Figure 3 and in the supplement Section 5. The extracted distribution of neurite lengths across multiple images are plotted in the right-most column in Figure 3.

**Figure 3.**
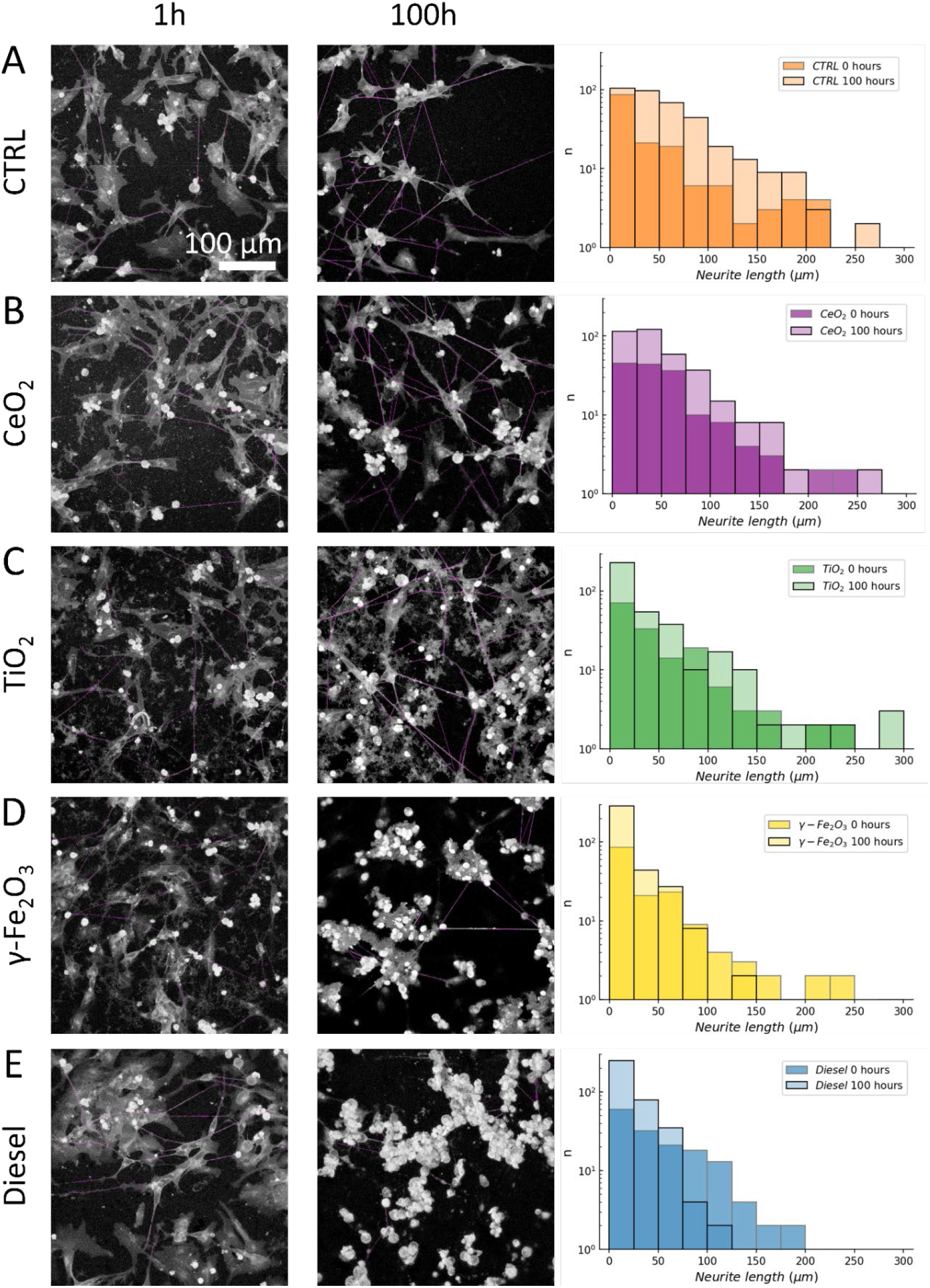
Neurite length shortening from 1 hour to 100 hours after exposure to nanomaterial. We determined the lengths of neurites that connected two distinct neuronal cells at 1 and 100 hours after exposure to nanomaterials using the NeuronJ plug-in. Representative images of neurites selected for analysis are shown in violet in the left (1h) and middle (100h) column. The right-most column shows the distributions of neurite lengths 1h (transparent) and 100h (solid with black edges) after exposure to different nanomaterials: A) control, B) TiO_2_ nanotubes, C) CeO_2_, D) γ-Fe_2_O_3_, and E) diesel exhaust particles exposed neurons. Median values of neurite lengths of all samples were significantly shortened for TiO_2_ nanotubes, γ-Fe_2_O_3_, and diesel exhaust particles (P<0.001, Mann–Whitney U test).

After 100 hours of exposure to nanomaterials, a significant reduction occurred in median length of neurites in neurons exposed to TiO_2_ nanotubes (from 59.4 μm to 39.8 μm, P<0.001), γ-Fe_2_O_3_ (from 56.8 μm to 32.2 μm, P<0.001), and diesel exhaust particles (from 60.7 μm to 29.6 μm, P<0.001), compared to the same sample after 1-hour exposure. In contrast, CeO_2_ did not exhibit any reduction in length even after 100 hours, compared the measurements taken an hour after exposure (Mann– Whitney U test applied due to the non-normal distribution of data). We observed the greatest effect in γ-Fe_2_O_3_ and diesel exhaust particles exposed neurons, where the median length decreased almost to a half of that observed in control. Alongside, the relative proportion of short neurites (<10 μm) dramatically increased, while there were almost no very long neurites (>100 μm) left (see histograms of number of neurites (n) versus neurite length in Figure 3 right).

### 3.4 Amyloid β (Aβ) plaques might have a protective role in neuronal survival after nanomaterial exposure

To gain insight into why some types of particulate matter (CeO_2_) do not affect neurite lengths and density to such an extent as other types (TiO_2_ nanotubes, γ-Fe_2_O_3_ and diesel exhaust particles) we analyzed the amount of Aβ deposited after nanomaterial exposure and assessed its colocalization with nanomaterial (Figure 4).

**Figure 4.**
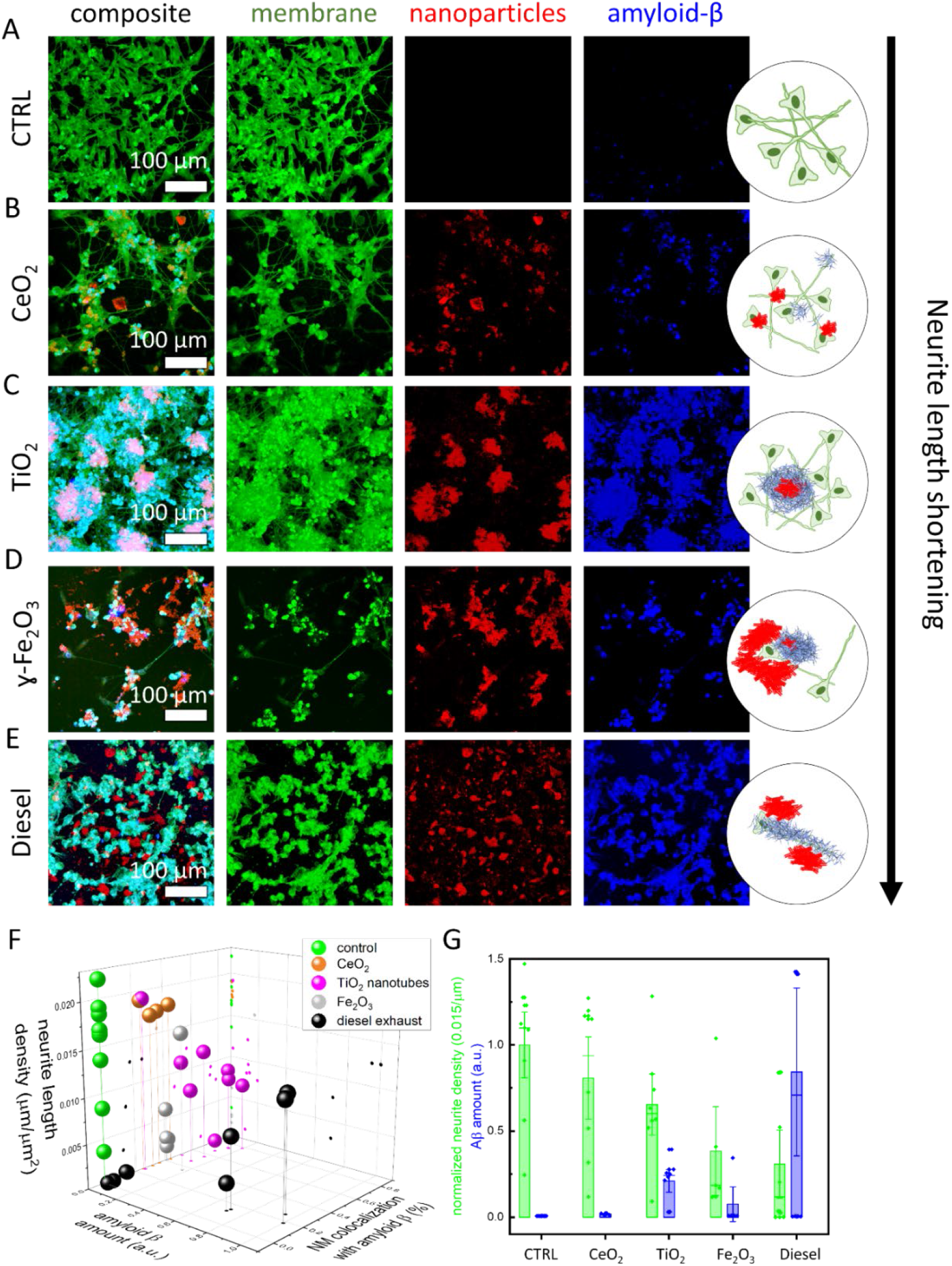
Particulate matter triggered amyloid β-containing plaques in neuronal cell culture 2 days after exposure to a nanomaterial. Confocal images of differentiated SH-SY5Y live neuronal cells were stained with fluorescent CellMask™ Deep Red Dye (membrane, green), unlabelled nanoparticles were detected via scattering (nanoparticles, red), Aβ plaques were immunostained with mouse monoclonal anti-β-Amyloid antibody (Santa Cruzblue, # sc-53822; blue color) conjugated with ATTO490 LS, antibodies were added to living cells in 10 nM concentration. A) control – unexposed neurons; B) neurons exposed to TiO_2_ nanotubes; C) CeO_2_ nanoparticles; D) γ-Fe_2_O_3_ nanoparticles; E) Diesel exhaust particles; F) neurite length density (z axis), amount of Aβ (x axis), and proportion of nanoparticles colocalized with Aβ (y axis), each symbol represents values obtained from one ROI; G) neurite density was calculated as total neurite length per total area (green bars), amount of Aβ was estimated as an integral of fluorescence intensity signal of Aβ antibody (blue bars), each symbol represents value obtained from one region of interest (ROI with a size of 400 μm x 400 μm, the height of the bar represents mean value, whiskers are standard deviation, long horizontal line is the median).

We first notice that all types of particles, except CeO_2_ nanoparticles (Figure 4 B), trigger the release of Aβ (compare the height of blue bars in Figure 4 G and the intensities of blue signal in Figure 4 A-E). TiO_2_ nanotubes triggered the formation of the large plaques containing neuronal plasma membrane (green), nanomaterial (red), and Aβ (blue), resulting in the overlapping signal of all three colors (Figure ^4^ C, light pink). TiO_2_ nanotubes were also the most efficiently covered with Aβ, up to about 80% (pink spheres in Figure 4 F).

On the other hand, iron oxide and diesel exhaust particles colocalized less with Aβ, about 40% and less than 20% on average, respectively (gray and black spheres in Figure 4 F). In addition, diesel exhaust also trigged the highest release of Aβ (blue bar in Figure 4 G), but these aggregates colocalized with neurons rather than the nanoparticles themselves (Figure 4 E; cyan color indicated colocalization of plasma membrane and Aβ signal). Furthermore, diesel exhaust particles caused massive neuronal death across the whole sample, as indicated by the presence of numerous rounded neurons with high membrane label intensity and no axons in Figure 4 E. Such effects were also observed by others in a coculture system (Block et al., 2004; Seo et al., 2023). Interestingly, in the regions where the amount of Aβ deposits was high and diesel exhaust particles colocalized with Aβ deposits, the lengths of neurites were about ten times longer compared to regions with minimal Aβ deposition. Each such region is presented by a black sphere with higher Aβ amount values in Figure 4 F. One can therefore see two groups of black spheres indicating the potential protective role of Aβ accumulation at the diesel exhaust particles. We speculate that this observation could potentially provide a mechanistic explanation to the conundrum observed in AD patients who possess a high burden of Aβ plaques but do not experience adverse symptoms (Gómez-Isla & Frosch, 2022).

In Figure 4 G, we show that exposure to nanomaterials that induce higher Aβ deposits (blue bars) generally leads to shortening of neurites (green bars) compared to the control. Despite inducing more Aβ deposits than γ-Fe_2_O_3_ nanoparticles, TiO_2_ nanotubes cause less neurite loss. This discrepancy might be attributed to the property of TiO_2_ nanotubes to bind more Aβ, thus again highlighting a protective role of Aβ deposition surrounding the nanomaterial.

These findings suggest that Aβ plaque formation does not necessarily coincide with neurite loss, which could explain why not all individuals with plaques develop AD (Gómez-Isla & Frosch, 2022), as well as why numerous clinical trials directed at reducing Aβ have failed in humans, despite their success in genetic mouse models of AD. Namely, it has been proposed that the reason for failures of clinical trials in humans is that mouse models only exhibit plaques without other hallmarks of the disease, whereas AD patients develop plaques, tau deposits, and neuronal degeneration (Busche & Hyman, 2020). Therefore, the induction of Aβ and tau plaque formation accompanied by neurite atrophy and neuronal degeneration through particulate matter exposure might represent a more relevant model of AD compared to existing genetically induced mouse models.

### 3.5 Conclusion

We here report the formation of extracellular Aβ plaques, resembling those found in AD, after exposure of an *in vitro* neuronal cell culture to engineered anatase TiO_2_ nanotubes, for which we previously showed to trigger chronic lung inflammation and nanoparticle-containing organic debris formation in mice (Danielsen et al., 2020; Kokot et al., 2020). These plaques closely resemble those in mice and AD patients observed by He et al. (He et al., 2018), which were induced in wild-type mice via intracerebral injection of Tau. Here we show that similar plaques can be triggered with exposure to some nanomaterials. We exposed wild-type SH-SY5Y-derived human neurons to a very high dose of engineered anatase TiO_2_ nanotubes and monitored the formation of Aβ and Tau-containing plaques 24 to 48 hours later. TiO_2_ nanotubes caused neurites (i.e., cellular protrusions extending from neurons) to shorten (median length decreased from 59.4 μm to 39.8 μm, P<0.001) and atrophy (Figure 2 C-D), causing decrease in neurite lengths (Figure 3 C). Oppositely, CeO_2_ nanoparticles do not cause neurite shortening (Figure 3 B) and do not induce Aβ plaque formation, which is consistent with their beneficial effect observed in traumatic brain injury (Fiorani et al., 2015; Gagnon & Fromm, 2015).

We also tested other nanomaterials that can be found in polluted air, diesel exhaust and iron oxide particles, which are associated with cognitive decline in areas with high air pollution (Underwood, 2017). Iron oxide (γ-Fe_2_O_3_ (from 56.8 μm to 32.2 μm, P<0.001)) and diesel exhaust particles (from 60.7 μm to 29.6 μm, P<0.001), decreased neurite lengths the most (Figure 3 C) and induced the accumulation of Aβ mainly in apoptotic neurons, as suggested by their rounded-up morphology, typical of apoptotic cells. On the other hand, TiO_2_ nanotubes induced the accumulation of Aβ in extracellular space often co-localized with the nanoparticles, suggesting that aggregation of the nanoparticles with Aβ might be a protective mechanism that "hides" material’s surface and prevents its harmful contact with neurons, thus safeguarding neurites.

Overall, this study identifies how certain nanoparticles relevant to particulate matter in ambient air pollution might contribute to AD by triggering hallmarks of the disease: shortening of neurites, formation of Aβ plaques and, in case of TiO_2_, accumulation of Tau protein within these plaques. Furthermore, we speculate that Aβ plaque colocalization with a nanomaterial could potentially protect neurites. While the exact mechanism behind this effect remains to be explained, it is possible that exposure to different types of ambient air pollution could explain earlier observations of individuals experiencing a high burden of plaques yet no adverse cognitive symptoms (Gómez-Isla & Frosch, 2022).

## Acknowledgements

The authors acknowledge the financial support from the Slovenian Research Agency (research core funding No. P1-0060 Experimental biophysics of complex systems and imaging in biomedicine, and No. P2-0089). The authors acknowledge the projects (Intelligent Content-Aware Nanospectroscopy (iCAN) of molecular events in nanoparticles-induced neurodegeneration, J7-2596, Concept development for mechanistic prediction of fibrosis and initiation of blood coagulation induced by inhaled materials (uCellnNet), N1-0240, L7-4535, J2-3043, J2-3040, J2-3046, J3-3079, and J7-4420, and bilateral ARRS project No. BI-FR/23-24-PROTEUS-005 (PR-12039), BI-US/22-24-100) financially supported by the Slovenian Research and Innovation Agency (SRA-ARIS). This work was also funded by European Union (grant no. 101092741, nanoPASS).

## 4 Author contributions

Conceptualization and design: A.S., T.K., I.U. and J.Š.; data collection: A.S.; analysis and interpretation of the data: A.S., T.K., I.U. and J.Š.; the drafting of the paper: A.S., T.K.; revising the paper critically for intellectual content: A.S., L.M.A.C, V. M., S.K., J.P., U.V., S.Z.N., I.U., T.K., and J.Š.; All authors gave the final approval of the version to be published; all authors agree to be accountable for all aspects of the work.

## 6 Data availability

The authors confirm that the data supporting the findings of this study are available upon request within the article and its supplementary materials.

