## Supplement for "Particulate matter constituents trigger the formation of extracellular amyloid β and Tau-containing plaques and neurite shortening *in vitro*"

* To whom correspondence should be addressed

### 1 Nanomaterial aggregation in cell medium

In the following images, we show the aggregation state of each material when suspended in the cell media. Each image represents measurements that were made on different days.

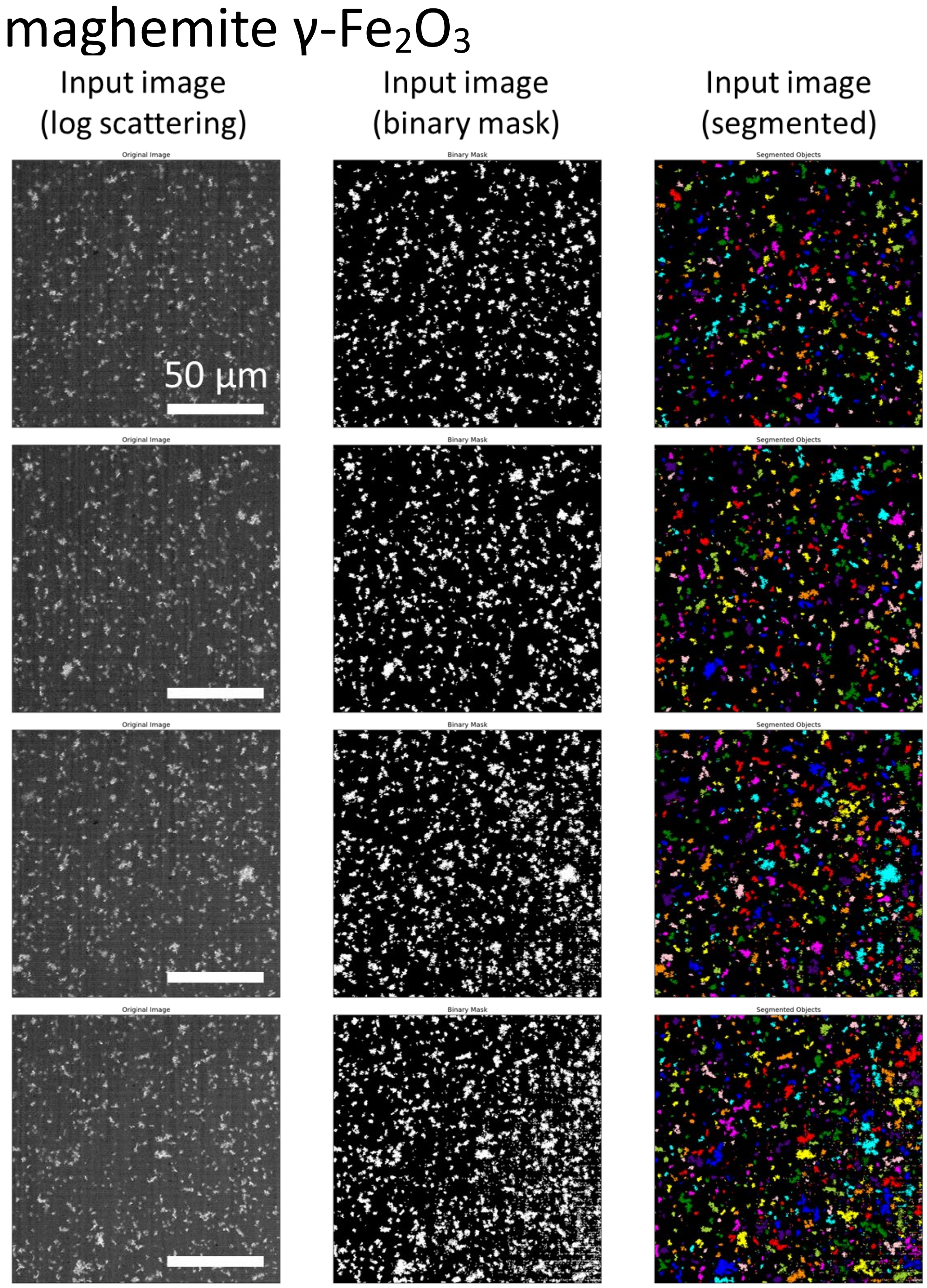

*Figure S 1: Log-transformed backscattered signal of 3 µl of maghemite γ-$Fe_2O_3$ resuspended in the neuronal DMEM/F-12 media. Imaged with the 60X magnification objective after overnight incubation to allow all the material to deposit.*

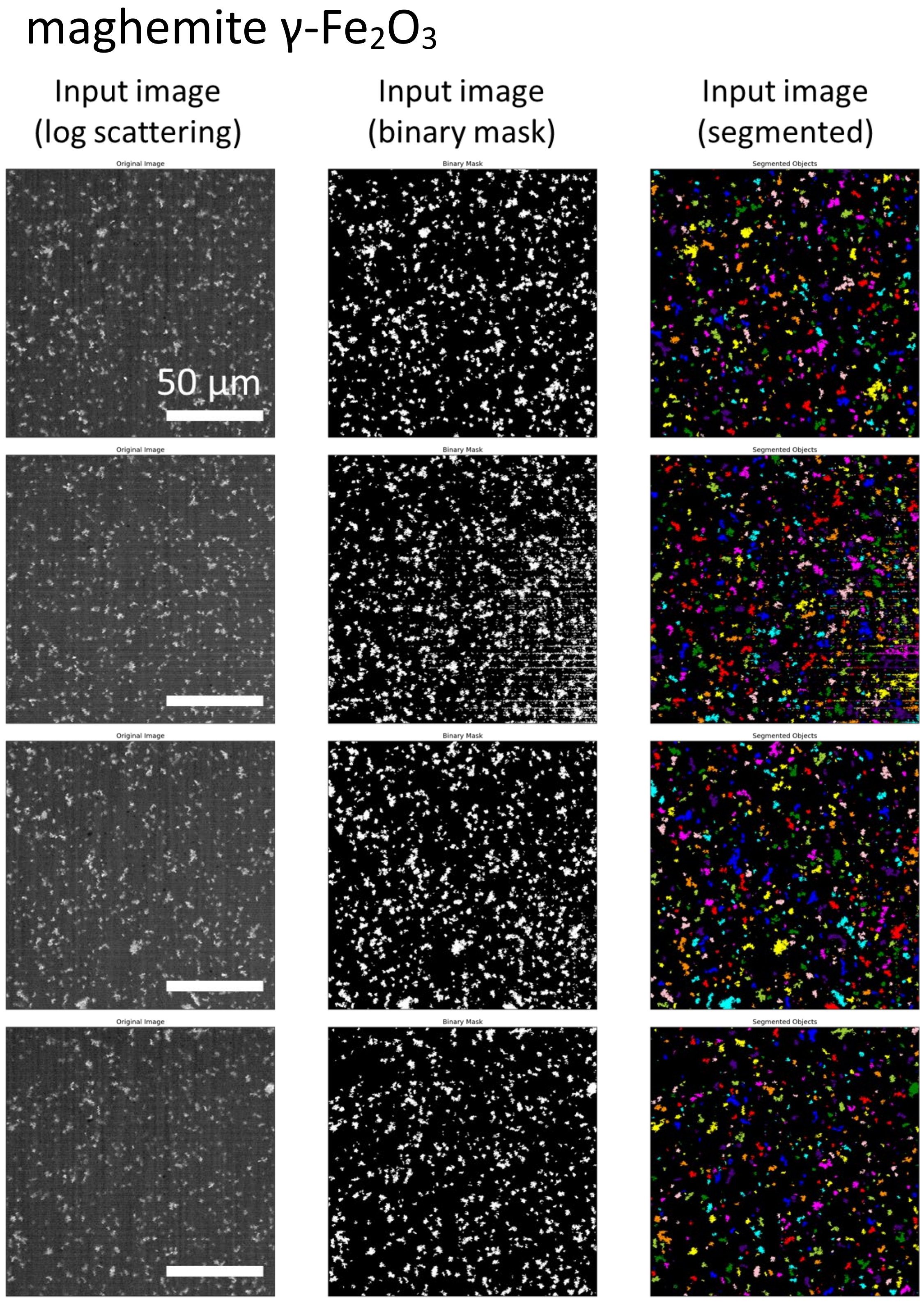

*Figure S 2: Log-transformed backscattered signal of 3 µl of maghemite γ-$Fe_2O_3$ resuspended in the neuronal DMEM/F-12 media. Imaged with the 60X magnification objective after overnight incubation to allow all the material to deposit.*

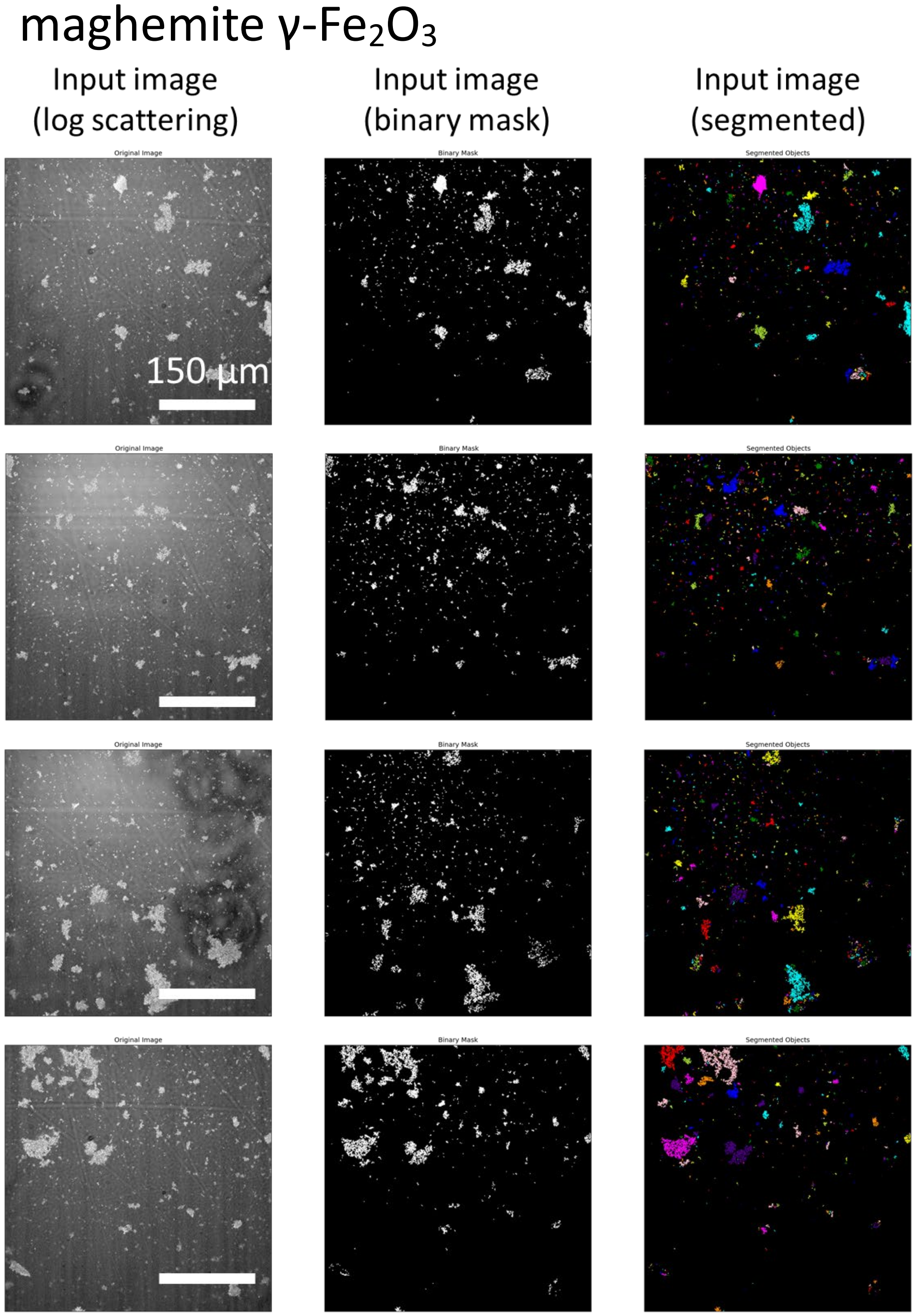

*Figure S 3: Log-transformed backscattered signal of 3 µl of maghemite γ-$Fe_2O_3$ resuspended in the neuronal DMEM/F-12 media. Imaged with the 20X magnification objective after overnight incubation to allow all the material to deposit.*

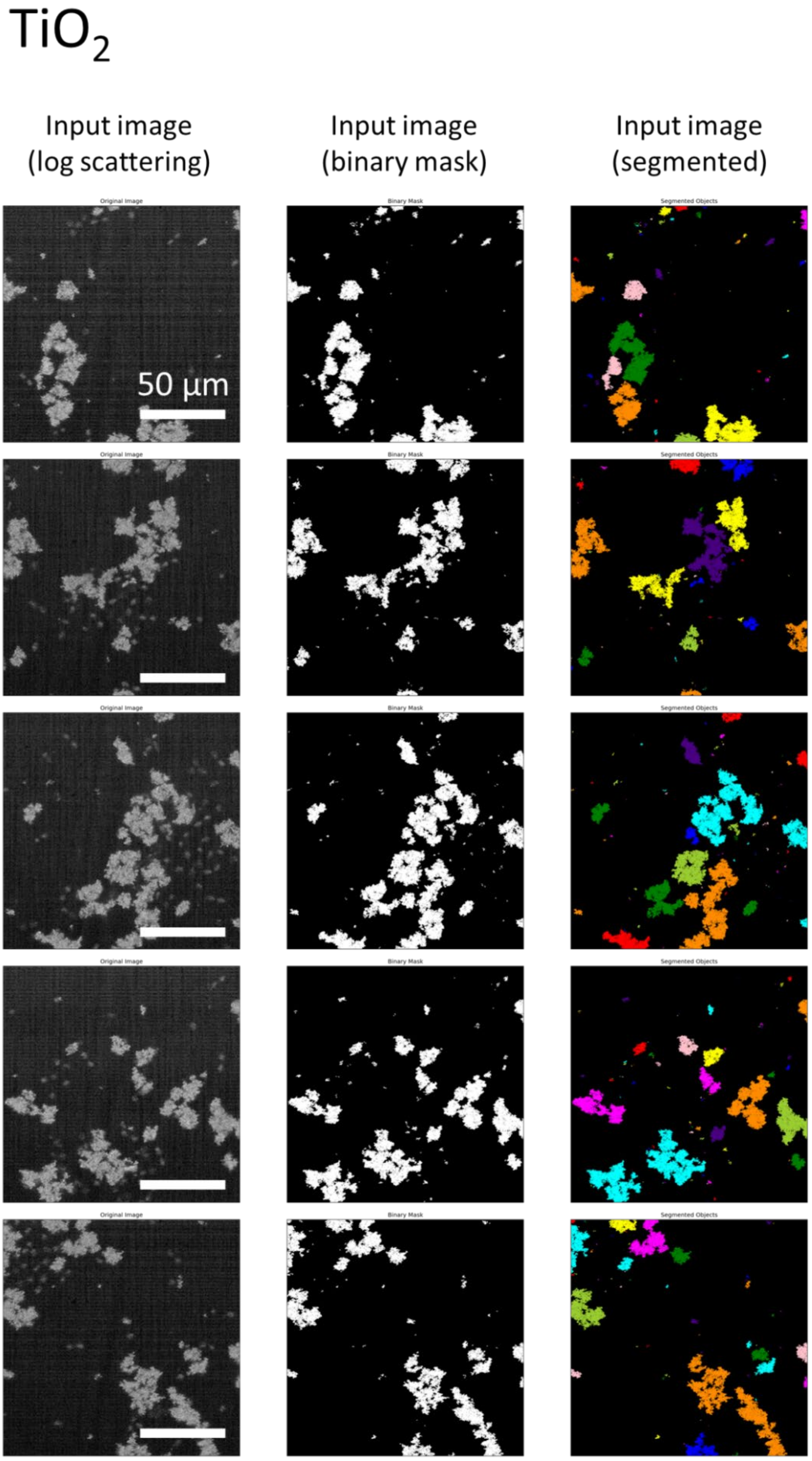

*Figure S 4: Log-transformed backscattered signal of 3 µl of $TiO_2$ resuspended in the neuronal DMEM/F-12 media. Imaged with the 60X magnification objective after overnight incubation to allow all the material to deposit.*

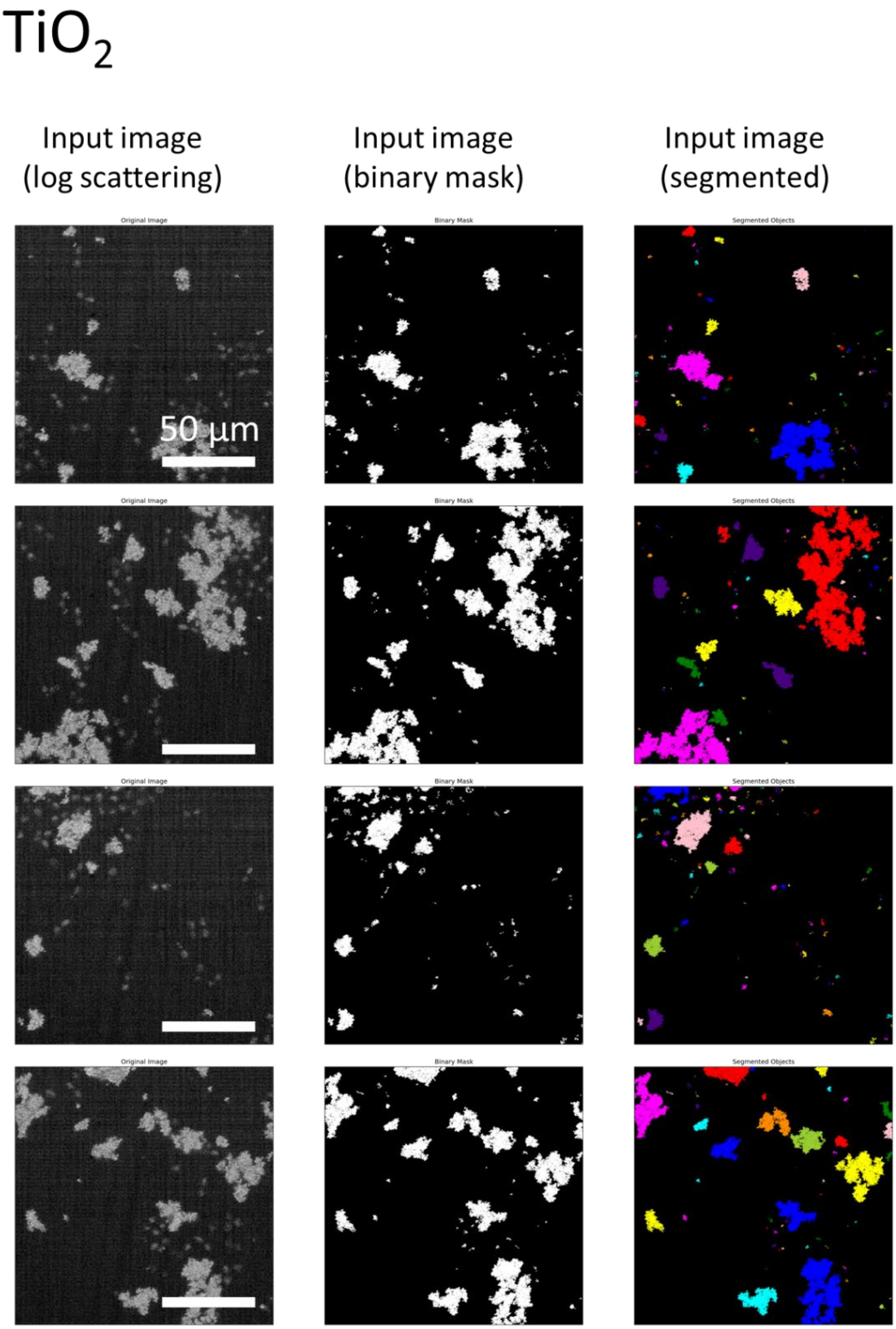

*Figure S 5: Log-transformed backscattered signal of 3 µl of $TiO_2$ resuspended in the neuronal DMEM/F-12 media. Imaged with the 60X magnification objective after overnight incubation to allow all the material to deposit.*

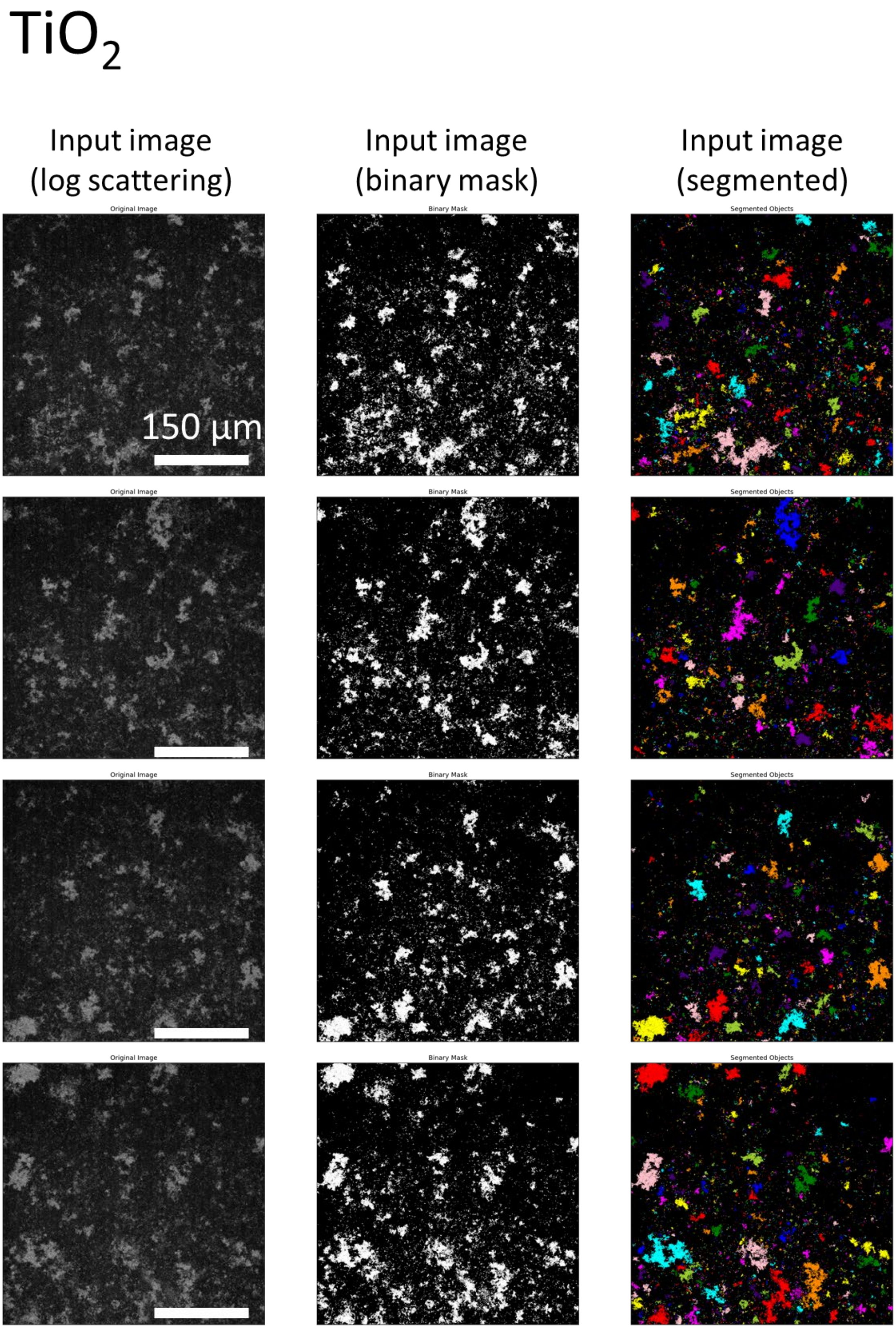

*Figure S 6: Log-transformed backscattered signal of 3 µl of $TiO_2$ resuspended in the neuronal DMEM/F-12 media. Imaged with the 20X magnification objective after overnight incubation to allow all the material to deposit.*

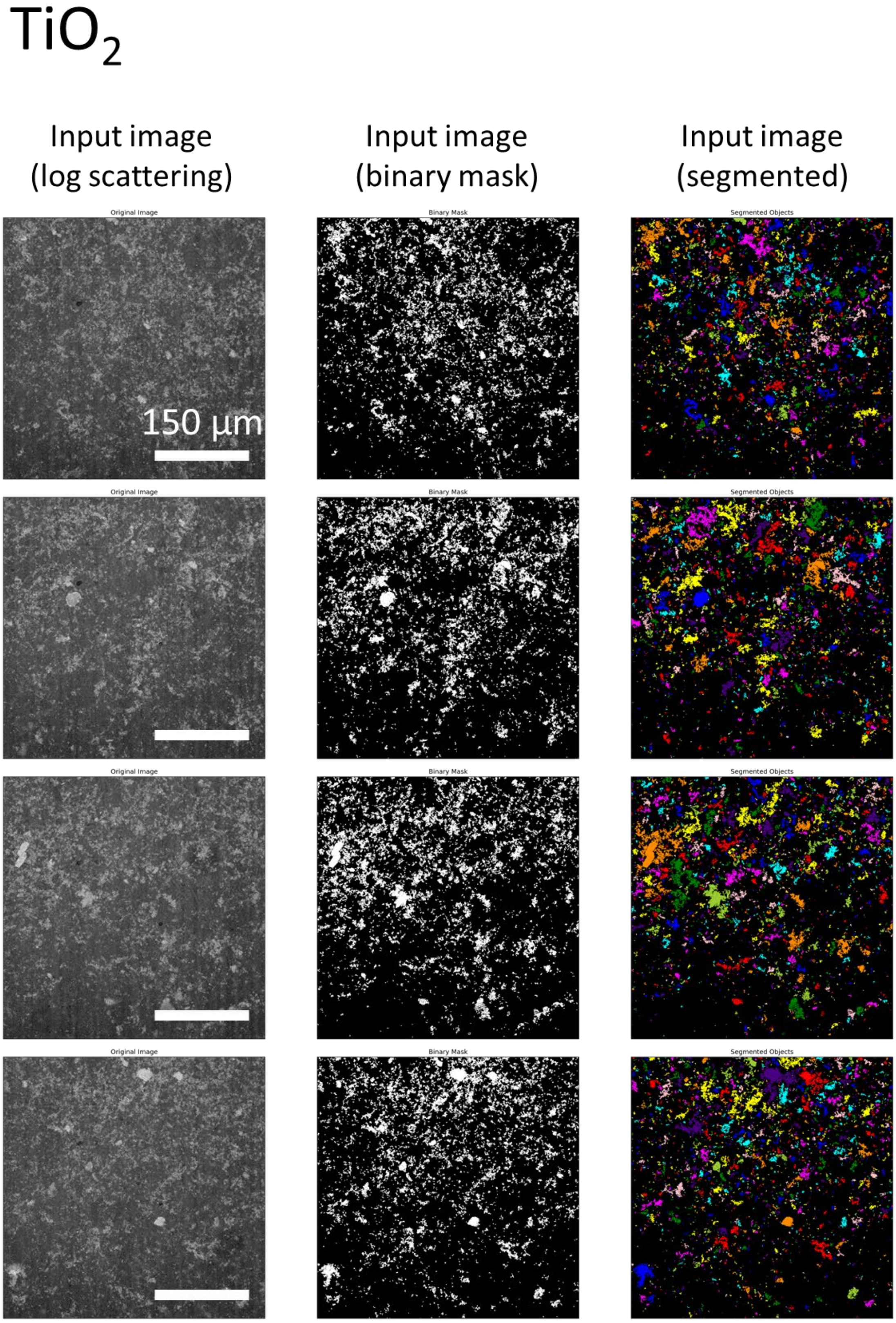

*Figure S 7: Log-transformed backscattered signal of 3 µl of $TiO_2$ resuspended in the neuronal DMEM/F-12 media. Imaged with the 20X magnification objective after overnight incubation to allow all the material to deposit.*

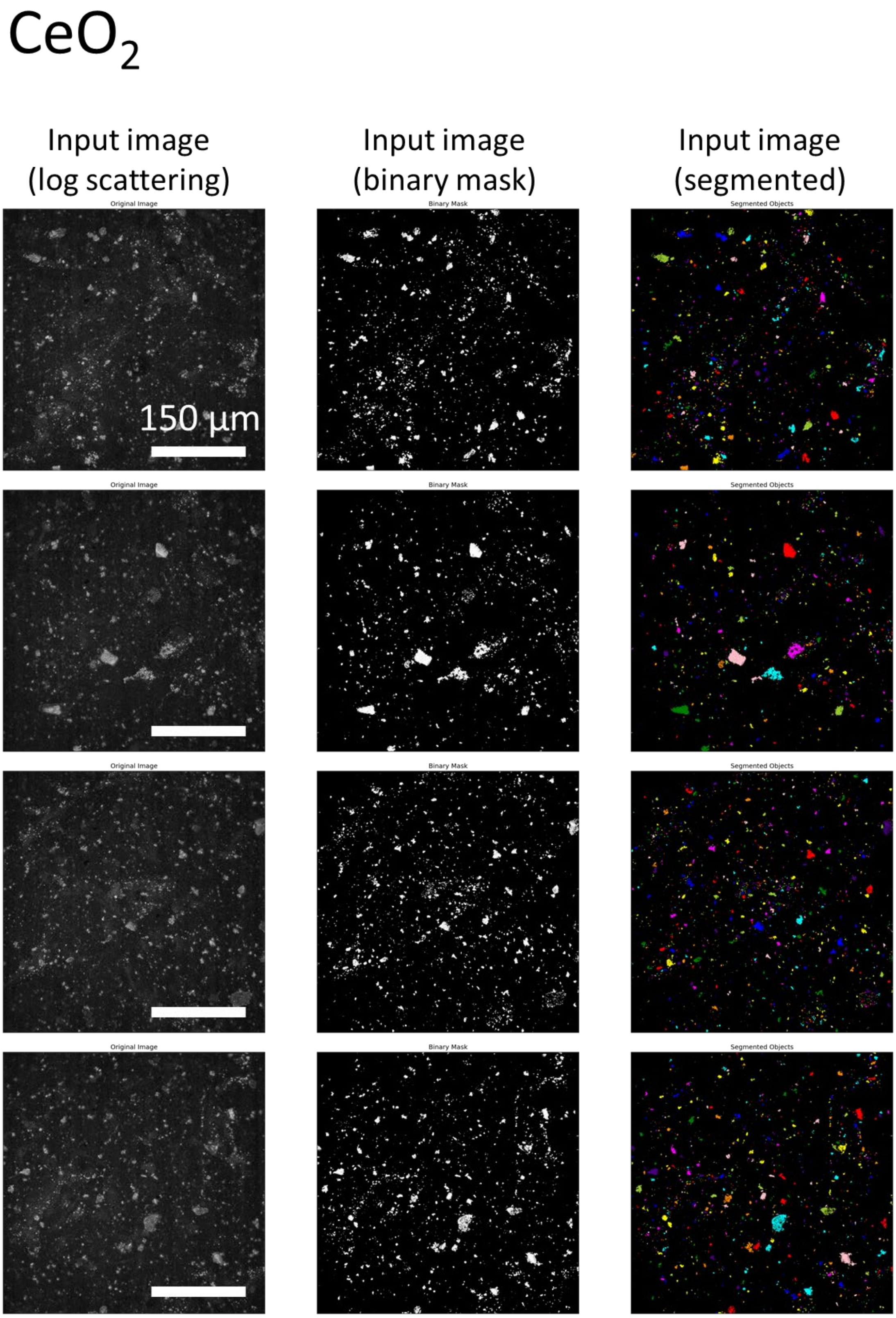

*Figure S 8: Log-transformed backscattered signal of 3 µl of $CeO_2$ resuspended in the neuronal DMEM/F-12 media. Imaged with the 20X magnification objective after overnight incubation to allow all the material to deposit.*

$CeO_2$

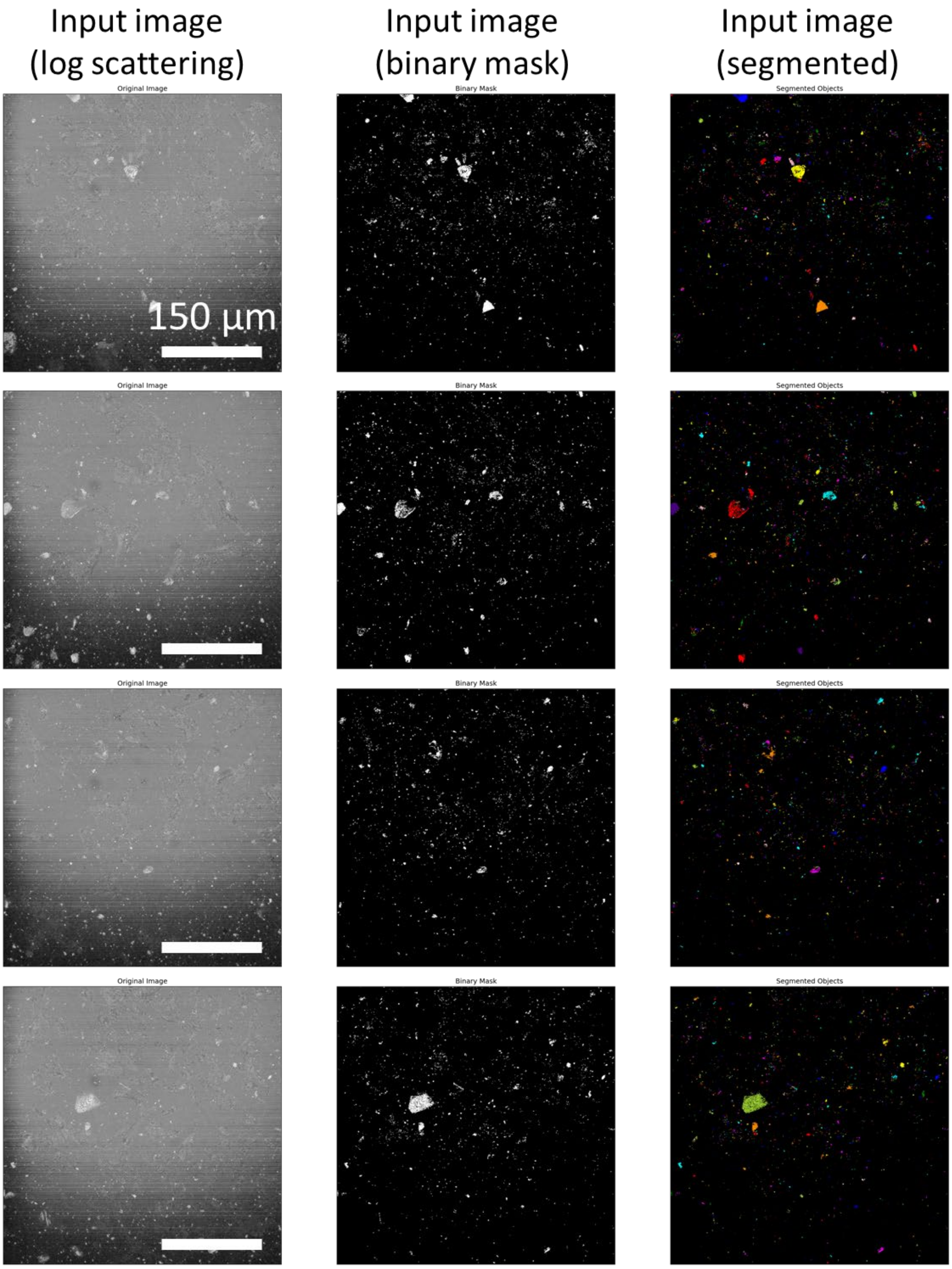

*Figure S 9: Log-transformed backscattered signal of 3 µl of $CeO_2$ resuspended in the neuronal DMEM/F-12 media. Imaged with the 20X magnification objective after overnight incubation to allow all the material to deposit.*

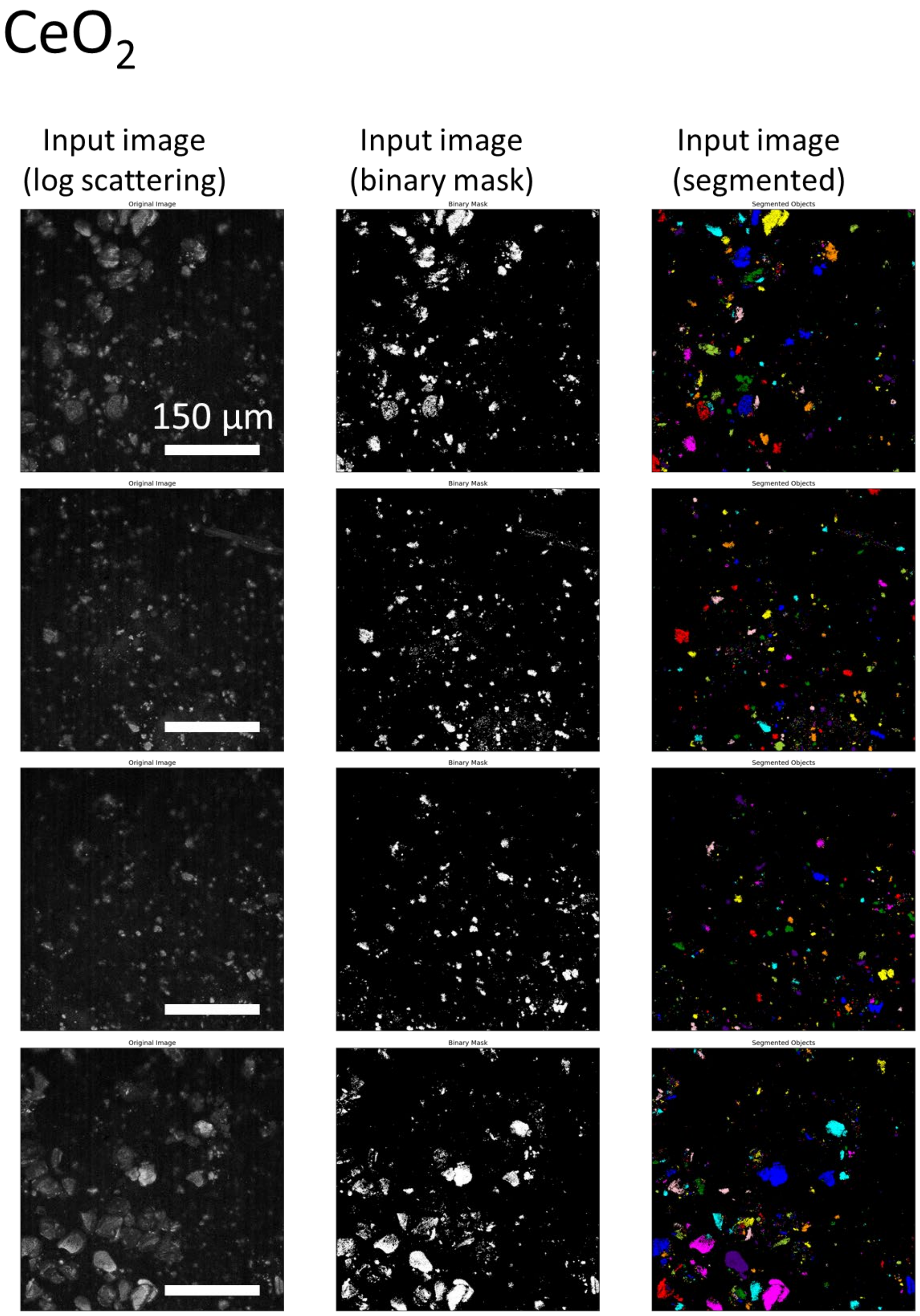

*Figure S 10: Log-transformed backscattered signal of 3 µl of $CeO_2$ resuspended in the neuronal DMEM/F-12 media. Imaged with the 20X magnification objective after overnight incubation to allow all the material to deposit.*

### Diesel

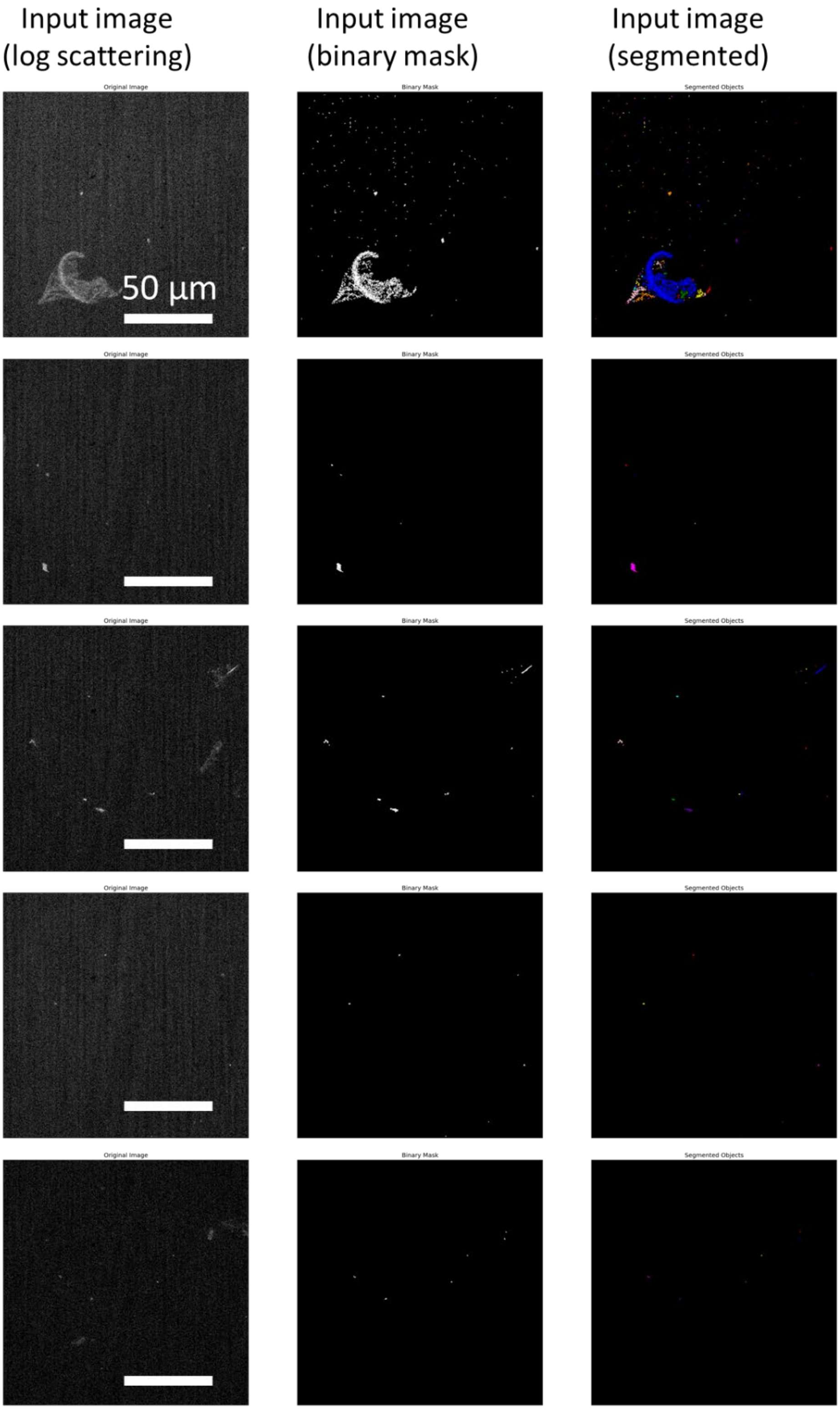

*Figure S 11: Log-transformed backscattered signal of 3 µl of Diesel resuspended in the neuronal DMEM/F-12 media. Imaged with the 60X magnification objective after overnight incubation to allow all the material to deposit.*

### Diesel

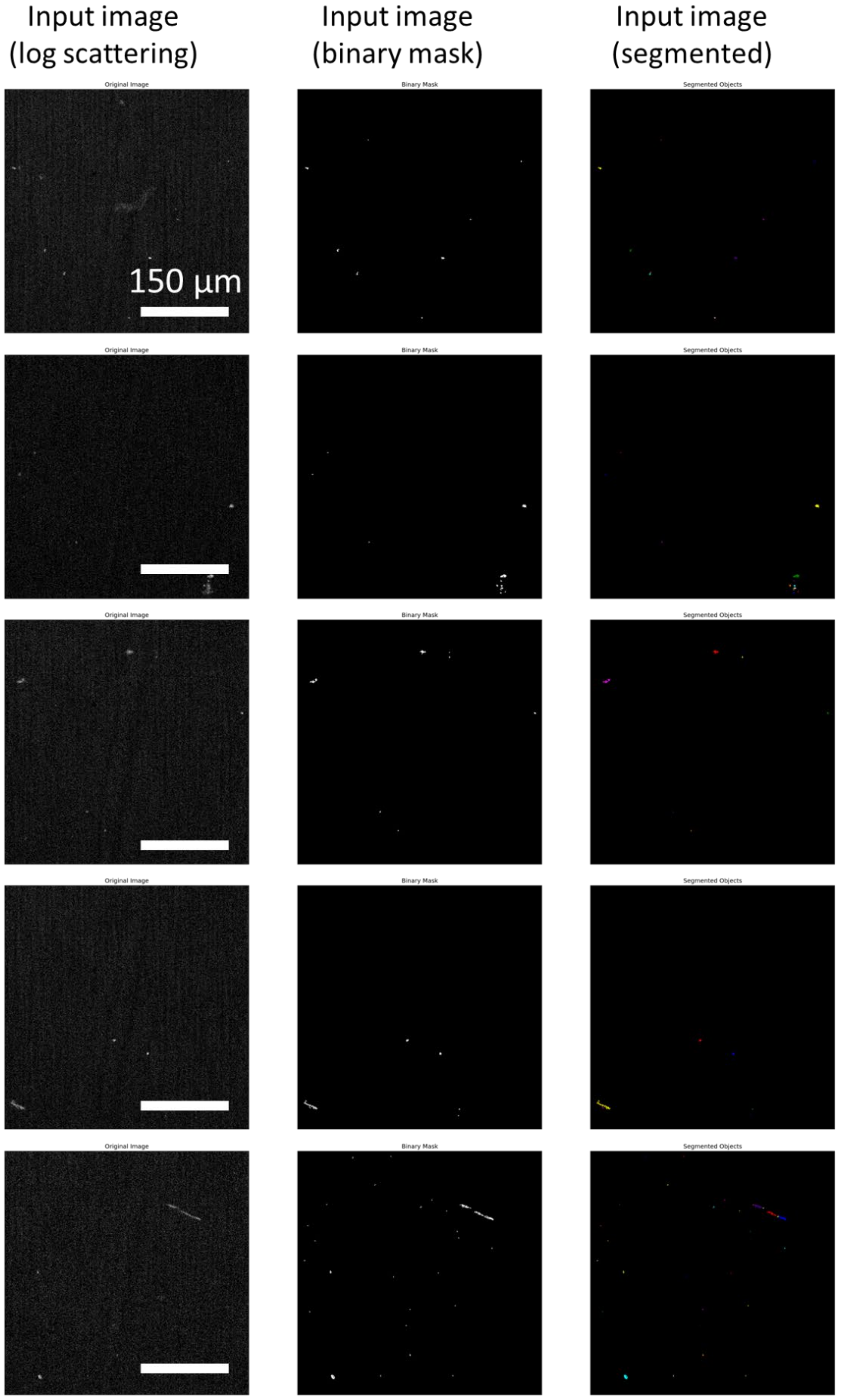

*Figure S 12: Log-transformed backscattered signal of 3 µl of Diesel resuspended in the neuronal DMEM/F-12 media. Imaged with the 20X magnification objective after overnight incubation to allow all the material to deposit.*

### Diesel

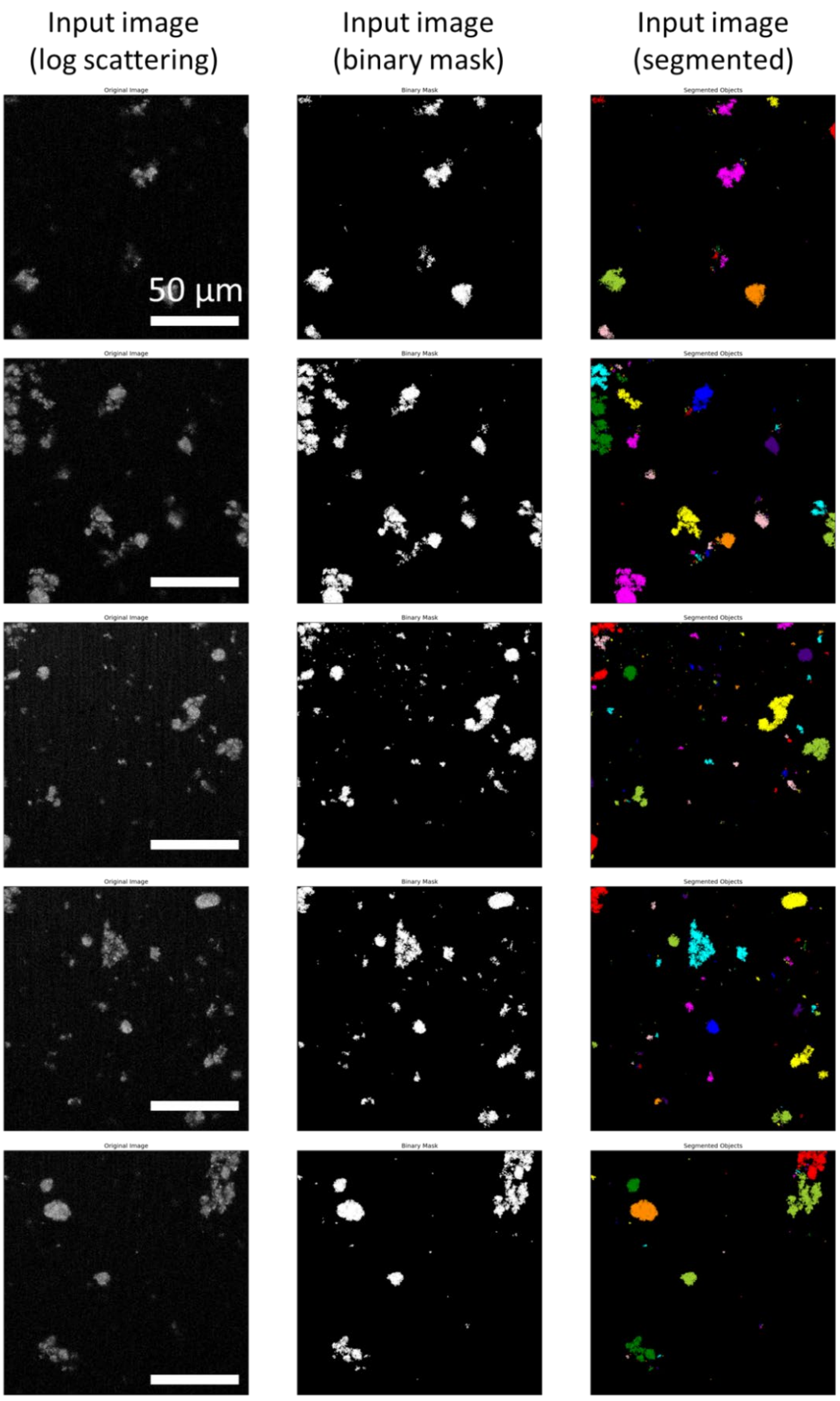

*Figure S 13: Log-transformed backscattered signal of 3 µl of Diesel resuspended in the neuronal DMEM/F-12 media. Imaged with the 60X magnification objective after overnight incubation to allow all the material to deposit.*

### Diesel

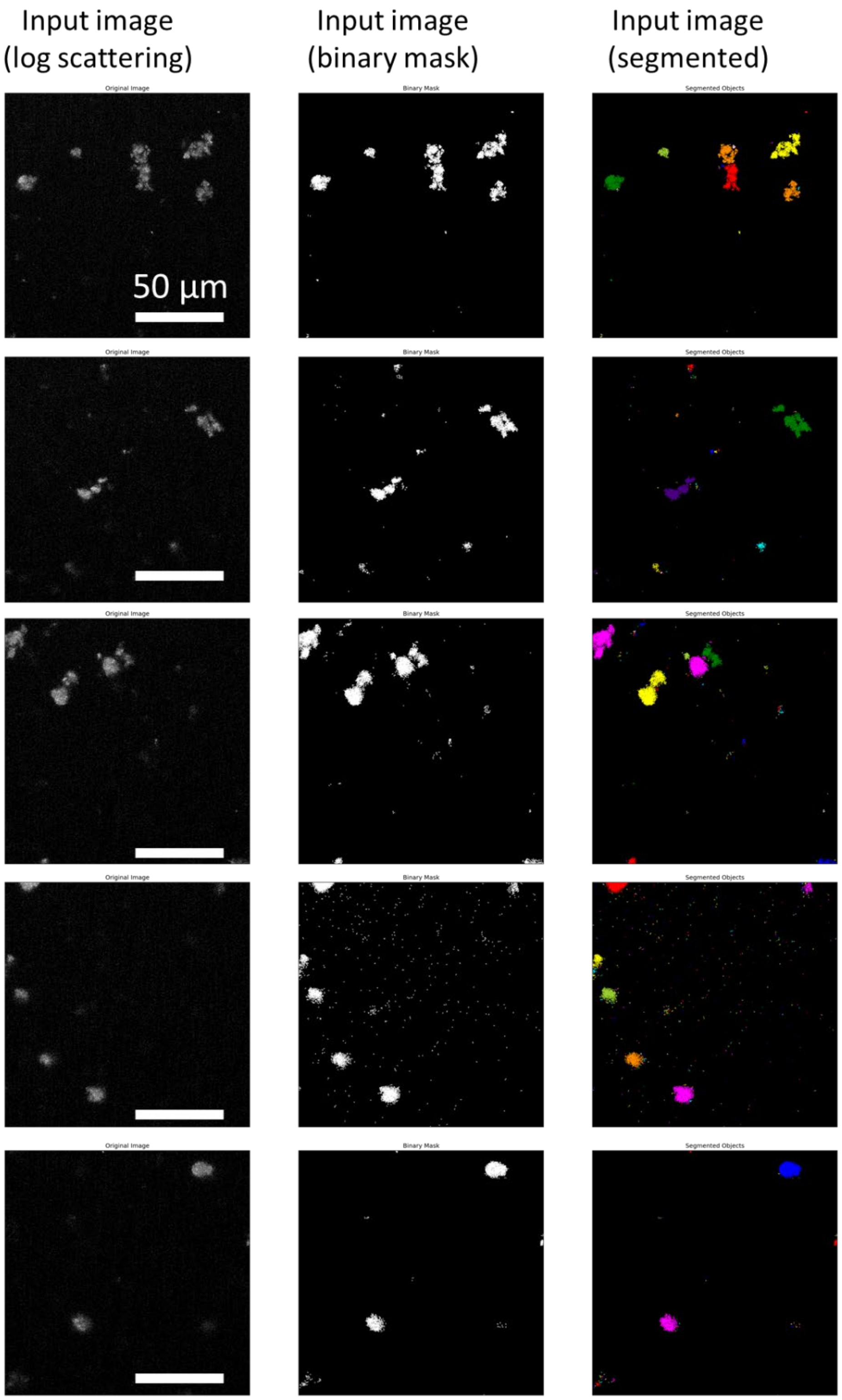

*Figure S 14: Log-transformed backscattered signal of 3 µl of Diesel resuspended in the neuronal DMEM/F-12 media. Imaged with the 60X magnification objective after overnight incubation to allow all the material to deposit.*

### Diesel

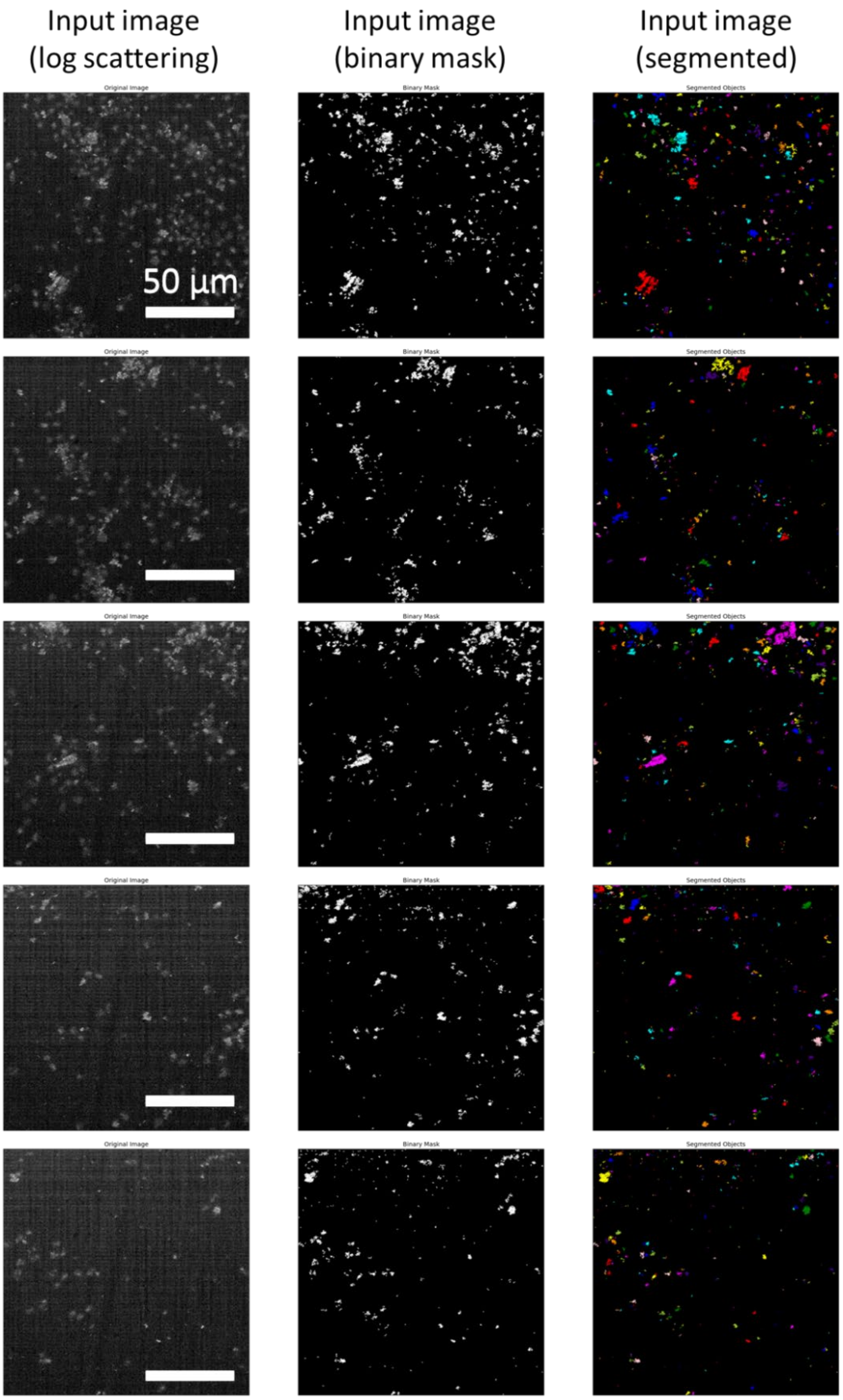

*Figure S 15: Log-transformed backscattered signal of 3 µl of Diesel resuspended in the neuronal DMEM/F-12 media. Imaged with the 60X magnification objective after overnight incubation to allow all the material to deposit.*

### Diesel

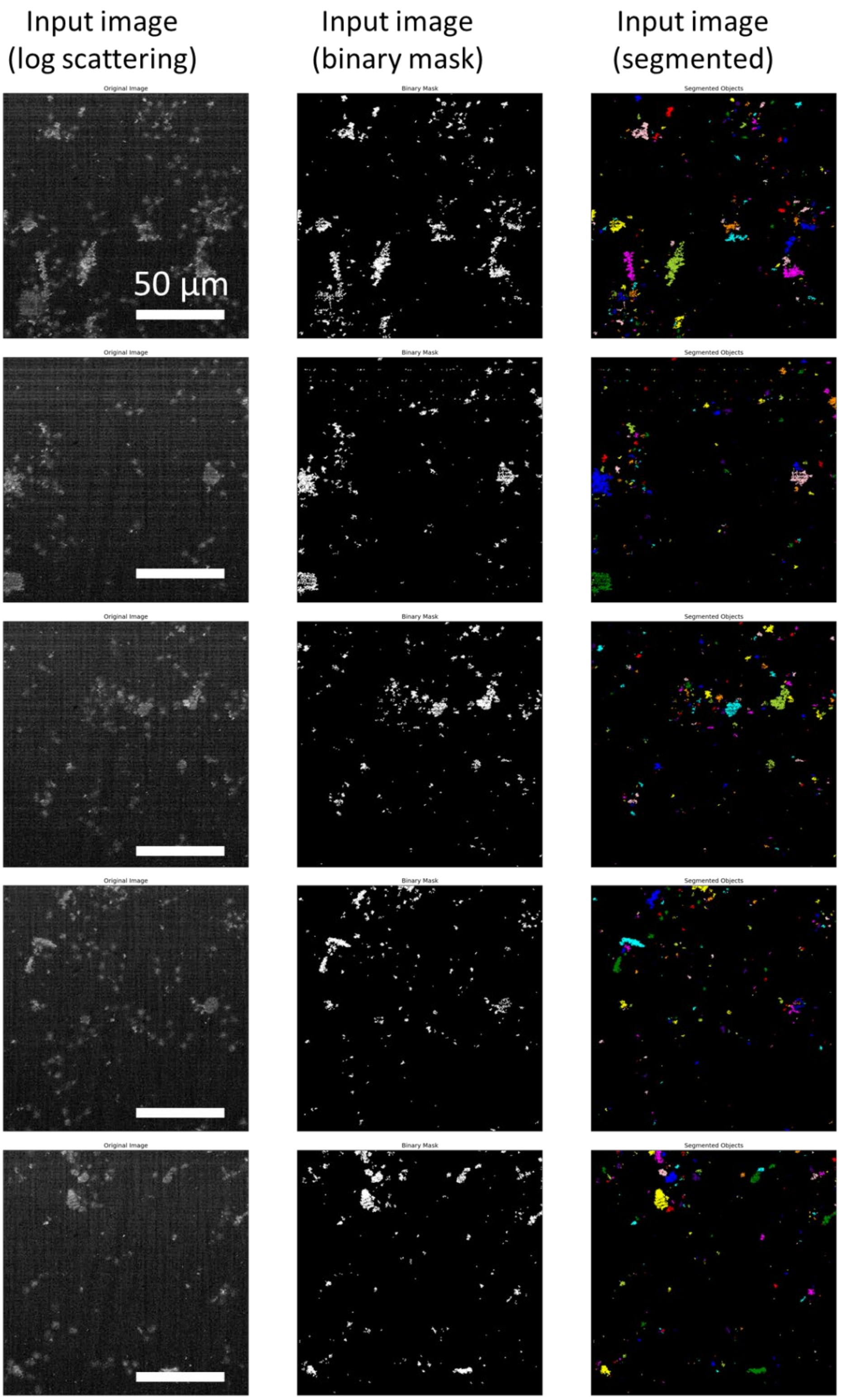

*Figure S 16: Log-transformed backscattered signal of 3 µl of Diesel resuspended in the neuronal DMEM/F-12 media. Imaged with the 60X magnification objective after overnight incubation to allow all the material to deposit.*

### Diesel

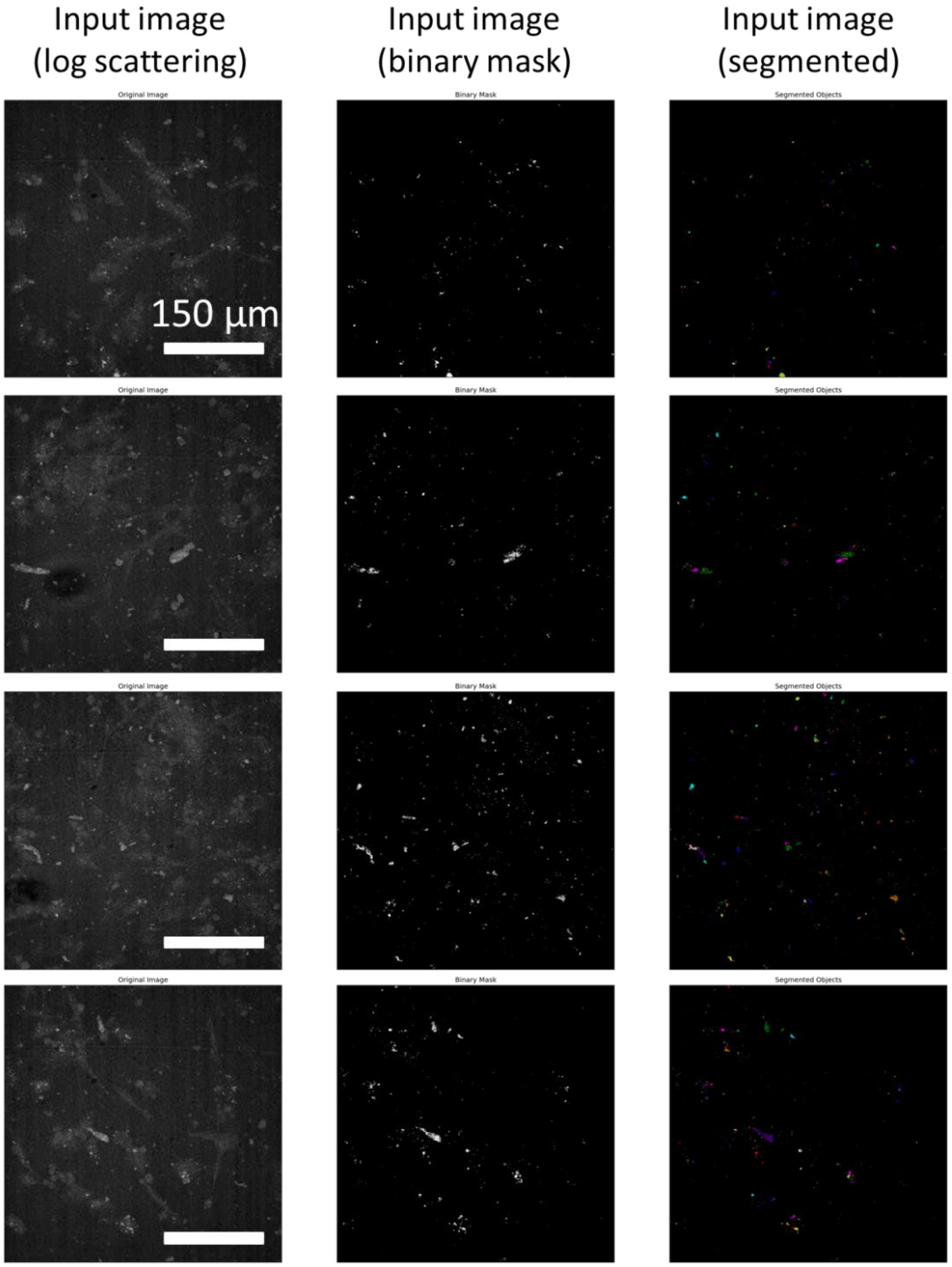

*Figure S 17: Log-transformed backscattered signal of 3 µl of Diesel resuspended in the neuronal DMEM/F-12 media. Imaged with the 60X magnification objective after overnight incubation to allow all the material to deposit.*

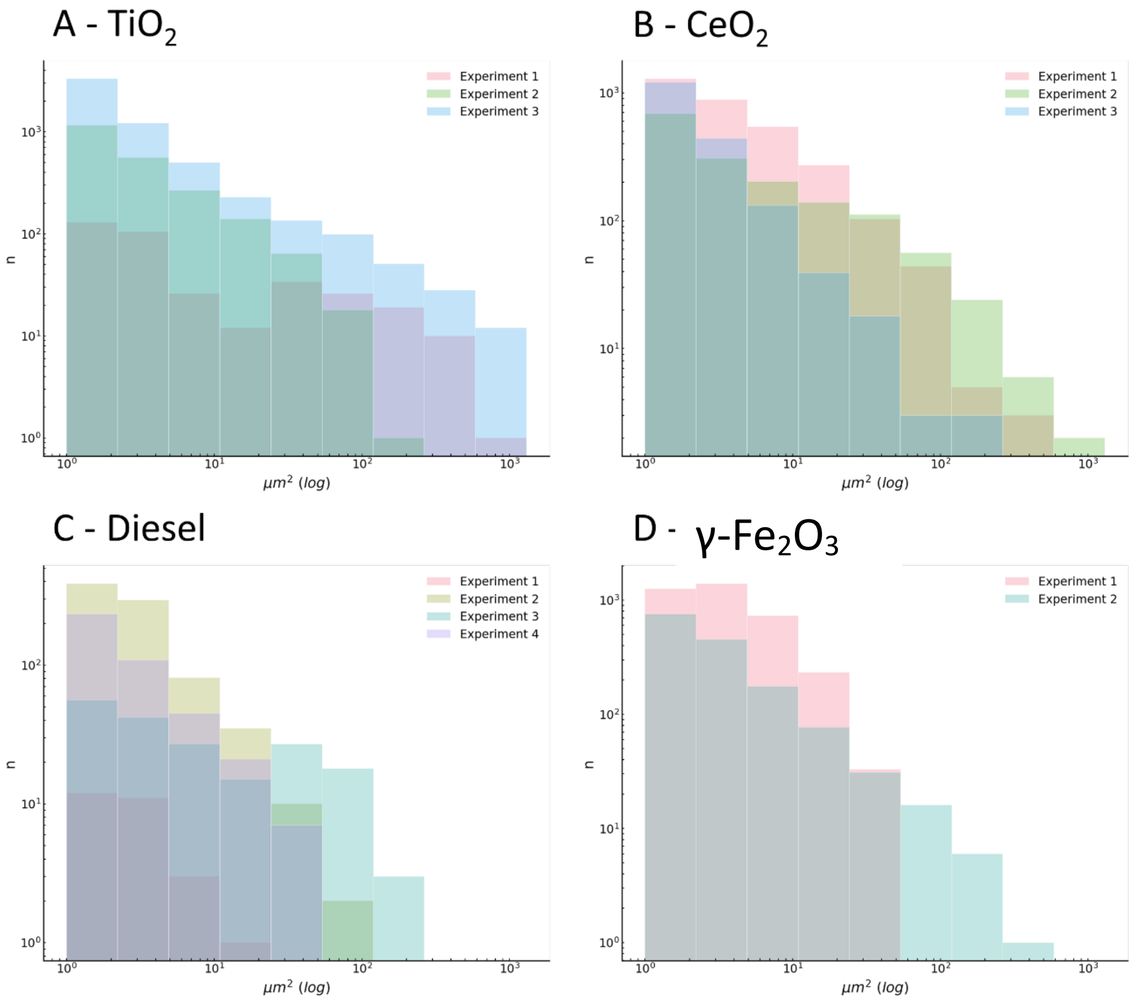

*Figure S 18: Log-log distribution of the size of individual particles and aggregates in terms of surface area ($\mu m^2$) resuspended in the neuronal cell media for A) $TiO_2$, b) $CeO_2$, C) Diesel, and D) maghemite γ-$Fe_2O_3$.*

### 2 Microscope setup

Following incubation with nanomaterial, the living samples were labelled and imaged using a state-of-the-art imaging system consisting of a Microscope Stage Top Chamber (Okolab H301-MIN) incubator mounted on the Olympus (Olympus IX83) confocal fluorescent microscope, which was equiped with the STED laser (Stedycon, Abberior). Stage top incubator maintains atmosphere with the 37°C, 5% $CO_2$, and >95% humidity to enable long term imaging of living cells.

The images were captured using a confocal microscope equipped with either a 20x magnification objective and 0.8 numerical aperture (NA) lens or a 60x magnification objective with a 1.2 NA lens. The microscope system incorporates four pulsed laser sources (Abberior) with a pulse duration of 120 ps and a maximum power of 50 µW at the sample plane. Additionally, four avalanche photodiode (APD) detectors are utilized for signal detection. Table provides the most frequently used combinations of lasers and filters employed during the experiments. The pulse repetition frequency was 80 MHz. The STED depletion laser, operating at a wavelength of 775 nm, had the same repetition frequency as the excitation lasers, a pulse duration of 1.2 ns, and a maximum power of 170 mW at the sample plane. In cases where the microscope settings were adjusted for maximal resolution during specific experiments, these customized settings will be described alongside the corresponding recorded images later in Supporting Information.

*Table S 1. Stedycon lasers and detection filters.*

| *Laser $\lambda_{EXCITATION}$ (nm)* | *Laser $\lambda_{EMMISION}$ (nm)* |
|---|---|
| 402 | 485-495 |
| 488 | 505-550 |
| 561 | 575-625 |
| 488 | 650-700 |

We successfully detected nanoparticles in the label-free, backscatter detection mode, utilizing the 488 / 488 ± 5 nm excitation / detection combo. Due to the large coherence of the laser, the backscattered light exhibited a strong speckle pattern, which was removed by Fourier transform bandpass filter (1 – 100 pixels) on scattering images. These images were subsequently binarized to obtain nanomaterial masks.

### 3 Main article FIGURE 1 supplementary material

SH-SY5Y cells were subjected to a differentiation process for seven days in DMEM/F12 cell media supplemented with 1% FBS and 10 µM retinoic acid (RA). The media containing RA was refreshed every other day to ensure optimal differentiation. Following the seven-day differentiation period, samples displaying noticeable changes in morphology and significant neurite outgrowth were carefully selected for further experimentation.

To visualize the live cell culture (green), a cytosolic fluorophore called CellTracker™ Green was employed. This fluorophore emits fluorescence exclusively within the cells. Additionally, a mouse monoclonal antibody targeting amino acids 1-40 of human amyloid β (Aβ) was used to detect amyloid precursor protein and Aβ (blue). Furthermore, a white fluorescence signal was obtained using the Tau-5 mouse monoclonal antibody, which specifically targets total tau. Both antibodies were added in 2 µg $mL^{-1}$ concentration before imaging.

Imaging was performed using a 20x magnification objective with a 0.8 numeric aperture lens, pixel size 250 nm, and the pixel dwell time 10 µs. Specific combinations of lasers and emission filters utilized during the imaging process are provided in Table S Table S2. A total of 15 regions of interest (ROIs), each measuring 400 x 400 µm, were scanned, resulting in the analysis of over 1000 neurons and more than 6000 plaques.

*Table S 3. Experimental setup and labels used for the analysis of extracellular plaque content in Figure 1.*

| *Label* | *$\lambda_{EXCITATION}$ (nm)* | *Laser power (%)* | *$\lambda_{EMMISION}$ (nm)* |
|---|---|---|---|
| CellTracker Green | 488 | 1 | 505-550 |
| anti-β-Amyloid Ab | 488 | 10 | 650-700 |
| Backscattering | 488 | 0.3 | 488 |

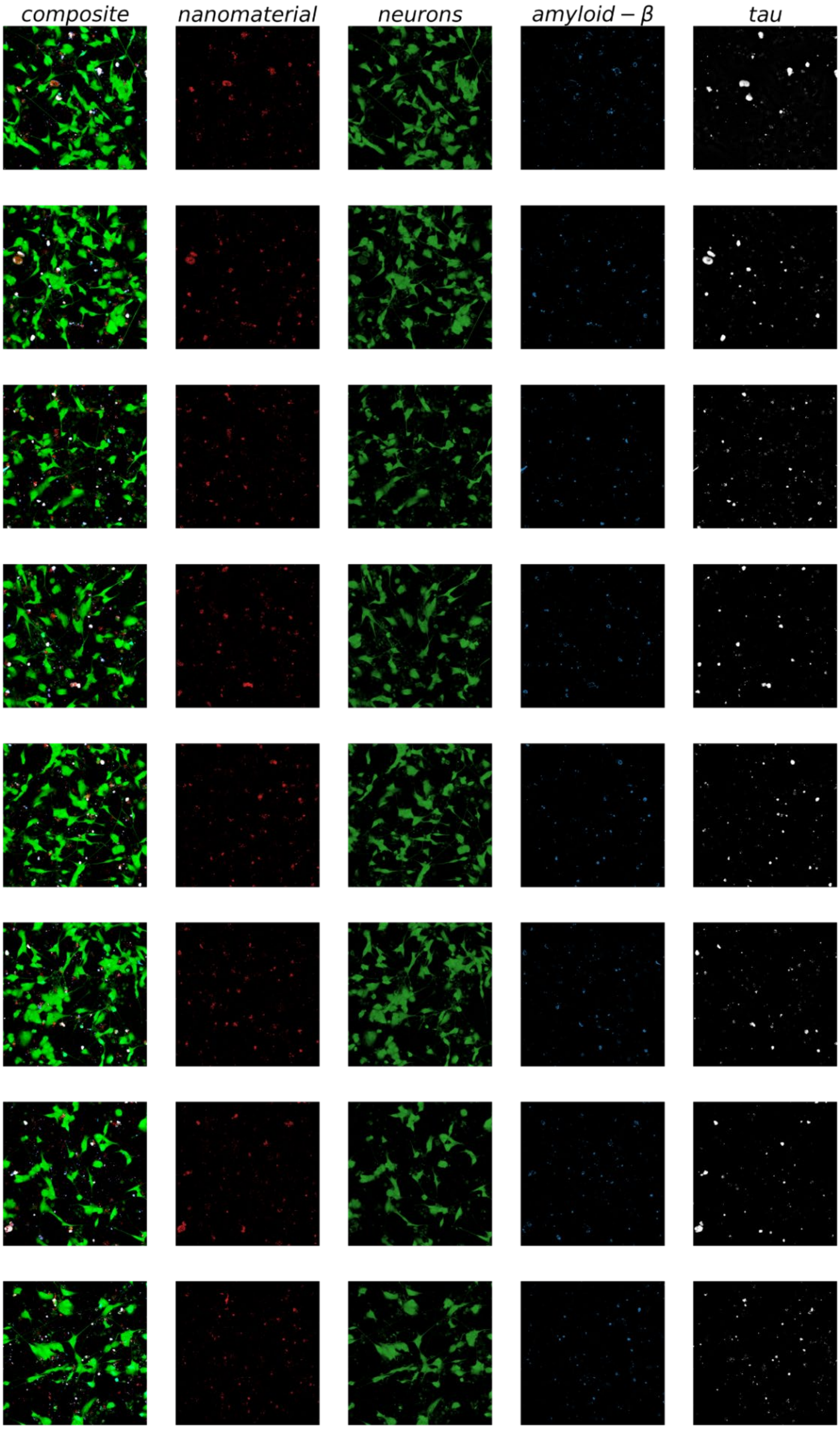
composite
nanomaterial
neurons
amyloid − β
tau

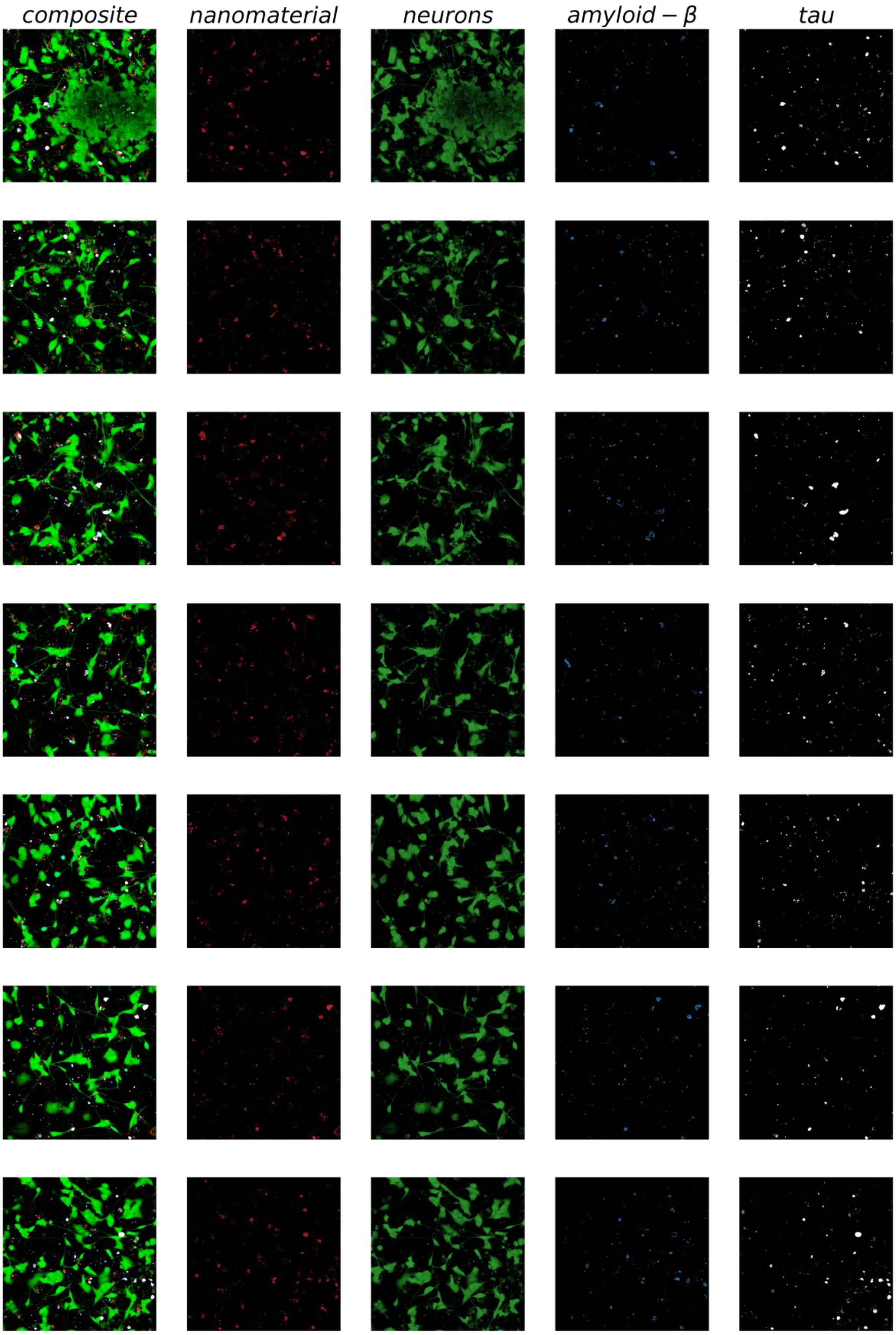

*Figure S 19: 15 ROIs analysed for the composition of TiO2 triggered Aβ plaques in Figure 1F-G. The first column shows the RGB image, while the subsequent columns display decomposed channels representing different components. Specifically, the second column represents the nanomaterial scattering signal (red), the third column represents the cytoplasmic CellTracker Green label (green), the fourth column represents the anti-Aβ antibody signal (blue), and the fifth column represents the anti-TAU antibody signal (gray).*

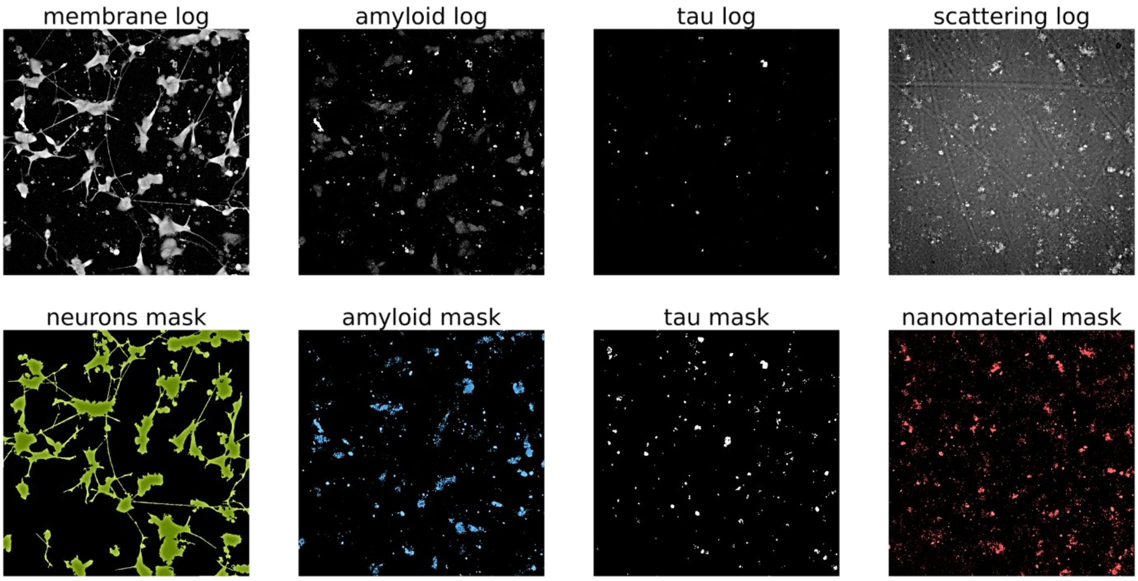

*Figure S 20: An example of how image binarization was performed on a single ROI. First, a Fast Fourier Transform (FFT) was applied to all log-transformed images. For the neuron mask, frequencies 2-1500 were kept. For Aβ, frequencies 1-50 were kept. For tau, frequencies 1-50 were kept. For the nanomaterial, frequencies 1-100 were kept. Afterward, reverse FFT images were thresholded to remove any remaining noise.*

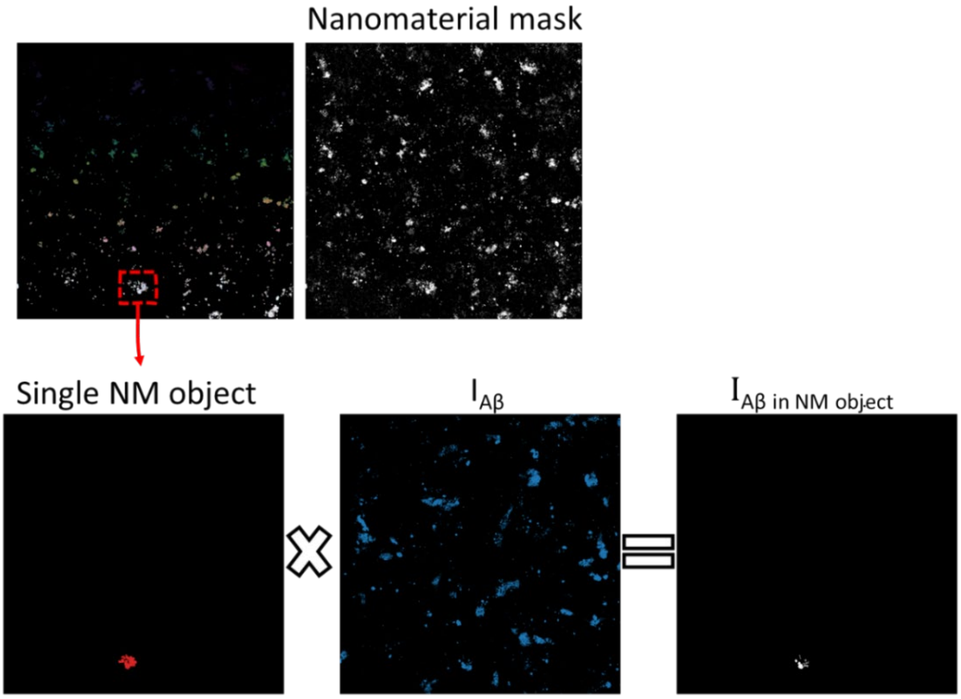

$$d_{A\beta} = \frac{\Sigma I_{A\beta \text{ in NM object}}}{S_{\text{NM object}}}$$

*Figure S 21: Example algorithm for calculating densities of plaque components. This algorithm focuses on the calculation of Aβ density The process begins with the generation of a binary mask of the nanomaterial using the FFT method, as explained before. Once the mask is obtained, unique IDs are assigned to each nanomaterial object, allowing for individual analysis. Next, a loop is conducted, processing each object individually. The nanomaterial object is multiplied with the input Aβ micrograph (IAβ), resulting in an image displaying Aβ fluorescence exclusively within that specific nanomaterial object (IAβ in NM object). The quotient of the sum of Aβ intensity within the object and the object's surface area provides the density of Aβ within that object (dAβ). The same algorithm is applied to the cytoplasmic and TAU signals.*

### 4 Main article FIGURE 2 supplementary material

Neurons were incubated with TiO2 nanotubes for a duration of 30 hours, followed by labelling and meticulous examination for the presence of TAU and Aβ-containing plaques. Imaging was conducted using a confocal microscope equipped with a 60x magnification objective and 1.2 NA lens.

For the time-lapse acquisition in Figure 3C, the pinhole was set at 40 µm (1.6 AU at 650 nm), with a pixel dwell time of 3.3 µs and a pixel size of 300 nm. The nanomaterial used were the same $TiO_2$ nanotubes used throughout the study, but they were labelled with Alexa 647. TiO2 labelling was explained in details previously(Kokot et al., 2020). The frame rate of the experiment was set to 5 seconds. Links to movies from the main articles are Movie 3C1 (https://portal.ijs.si/nextcloud/s/sX5weEJQGf5oqCc for Figure3 C1) and Movie 3C2 https://portal.ijs.si/nextcloud/s/DZrcdcJ3CQMesAo for Figure 3C2.

*Table S 4. Experimental setup for time-lapse in Figure 2C.*

| *Label* | *$\lambda_{EXCITATION}$ (nm)* | *Laser power (%)* | *$\lambda_{EMMISION}$ (nm)* |
|---|---|---|---|
| CellTracker Green | 488 | 10 | 505-550 |
| Alexa 647 | 641 | 0.3 | 651-691 |

To increase the resolution and enhance our understanding of the observed axonal damage, we employed a 3D super resoluton STED microscopy displayed in Figure 2A-C. Experimental setup is layed out in Table S5. The pixel size was 60 nm, pinhole 40 µm (1.01 AU @650 nm), and pixel dwell time 10 µs with 1 line accumulation. 3D stacks in Figure 3A and B were acquired with 1µm increments for a total of 7 µm in hight.

*Table S 5. Experimental setup for STED images in Figure2D.*

| *Label* | *$\lambda_{EXCITATION}$ (nm)* | *Laser power (%)* | *$\lambda_{EMMISION}$ (nm)* | *STED power (%)* |
|---|---|---|---|---|
| CellTracker Green | 488 | 1 | 505-550 | 50 |
| anti-β-Amyloid Ab | 488 | 10 | 650-700 | 74 |
| anti-TAU Ab | 640 | 10 | 650-700 | 74 |
| Backscattering | 488 | 0.3 | 488 | / |

### 5 Main article FIGURE 3 supplementary material

Neurite lengths were analysed using the NeuronJ plug-in, which is available at the following link: https://imagescience.org/meijering/software/neuronj/. The analysis was performed in Fiji ImageJ. Manual measurements, including axonal tracing, were conducted to determine the lengths of axons, dendrites, and neurites that connected two distinct cells. Images used in the analysis are depicted in Figure S 22 - Figure S 31. Neurites that were traced and included in analysis are overlaid with the violet colour, a default output of NeuronJ plug-in. Subsequently, these measurements were used to plot the distributions of neurite lengths of neurons (Figure S 32 - Figure S 35) Mann–Whitney U test was performed on the neurite lengths of control, $TiO_2$, $CeO_2$, γ-$Fe_2O_3$, and diesel exposed neurons due to the non-normal distribution of data. Significant difference occurred between the neurite lengths of control and γ-$Fe_2O_3$and Diesel exhaust nanoparticles exposed neurons and between $TiO_2$ and Diesel exhaust particles exposed neurons.

*1 hour after exposure:*

CTRL

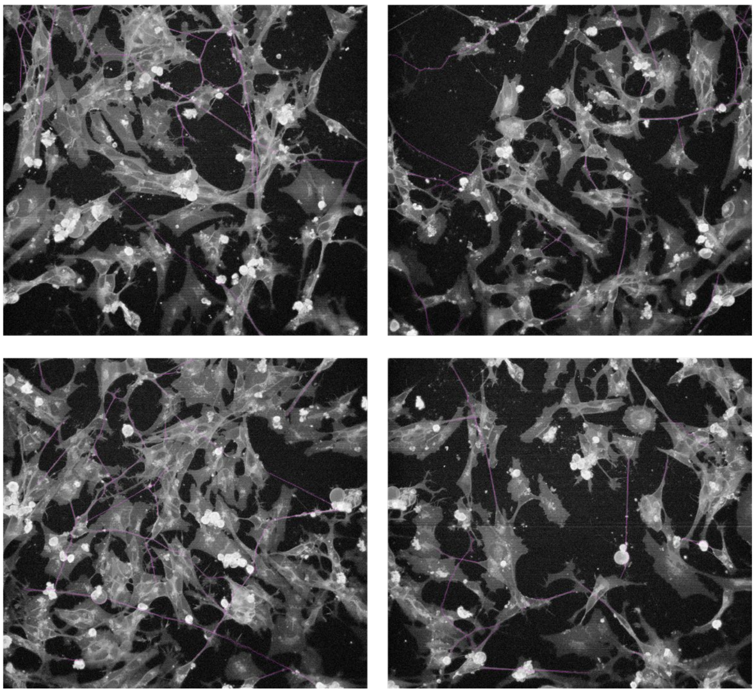

*Figure S 22 Images of unexposed control neurons used for the analysis of neurite length (violet) after 1h of incubation.*

$TiO_2$

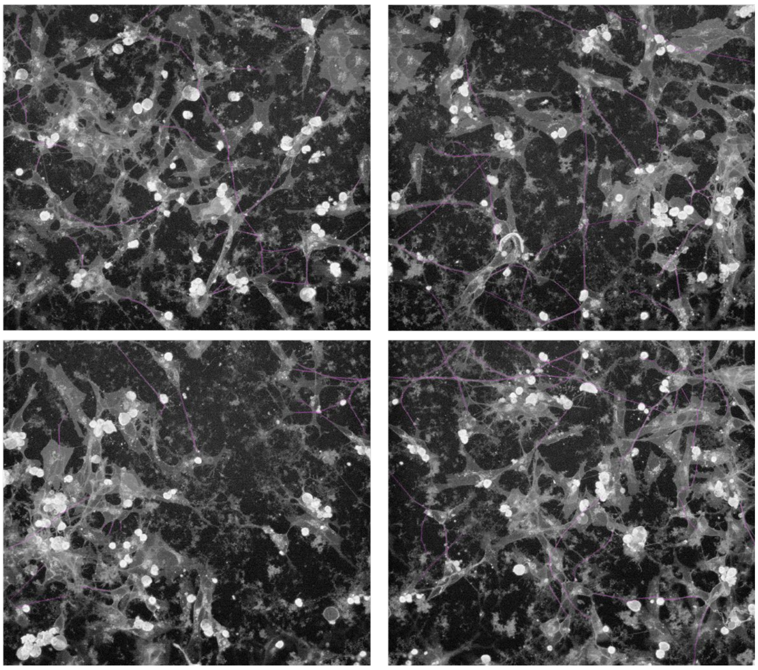

*Figure S 23 Images of $TiO_2$ exposed neurons used for the analysis of neurite length (violet) after 1h of incubation.*

maghemite γ-$Fe_2O_3$

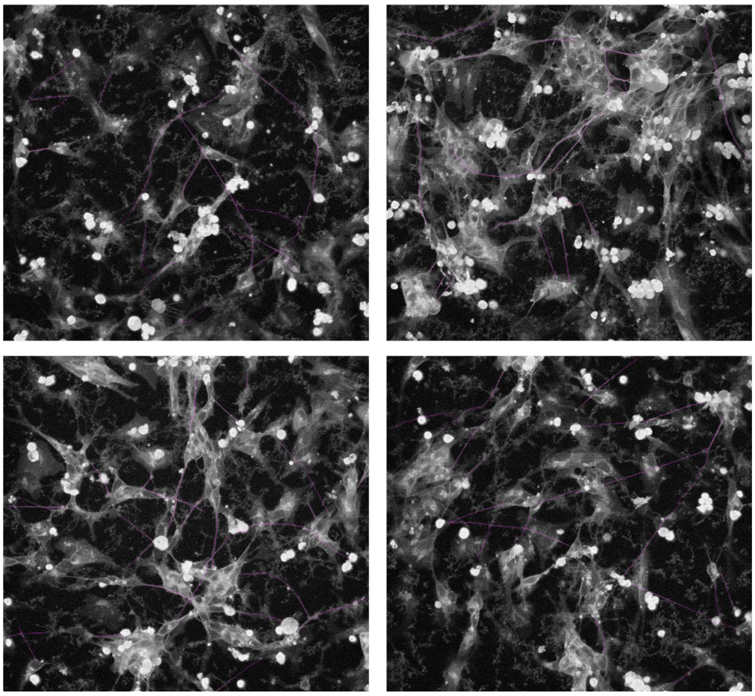

*Figure S 24 Images of $Fe_2O_3$ exposed neurons used for the analysis of neurite length (violet) after 1h of incubation.*

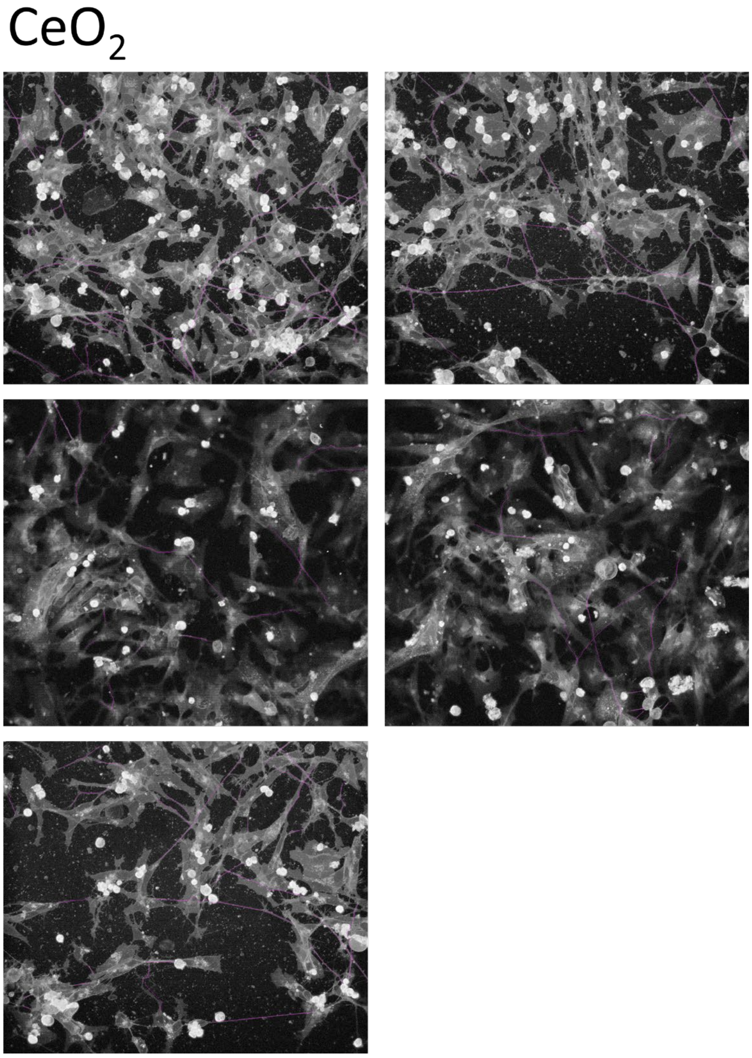

*Figure S 25 Images of $CeO_2$ exposed neurons used for the analysis of neurite length (violet) after 1h of incubation.*

### Diesel

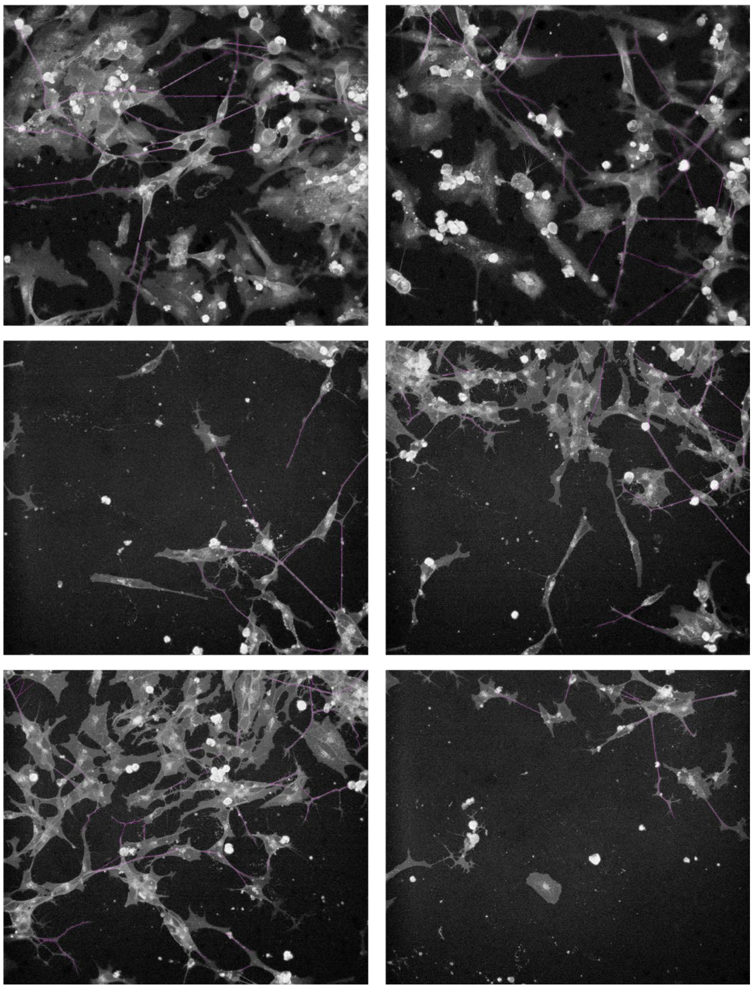

*Figure S 26 Images of diesel exposed neurons used for the analysis of neurite length (violet) after 1h of incubation.*

*100 hours after exposure:*

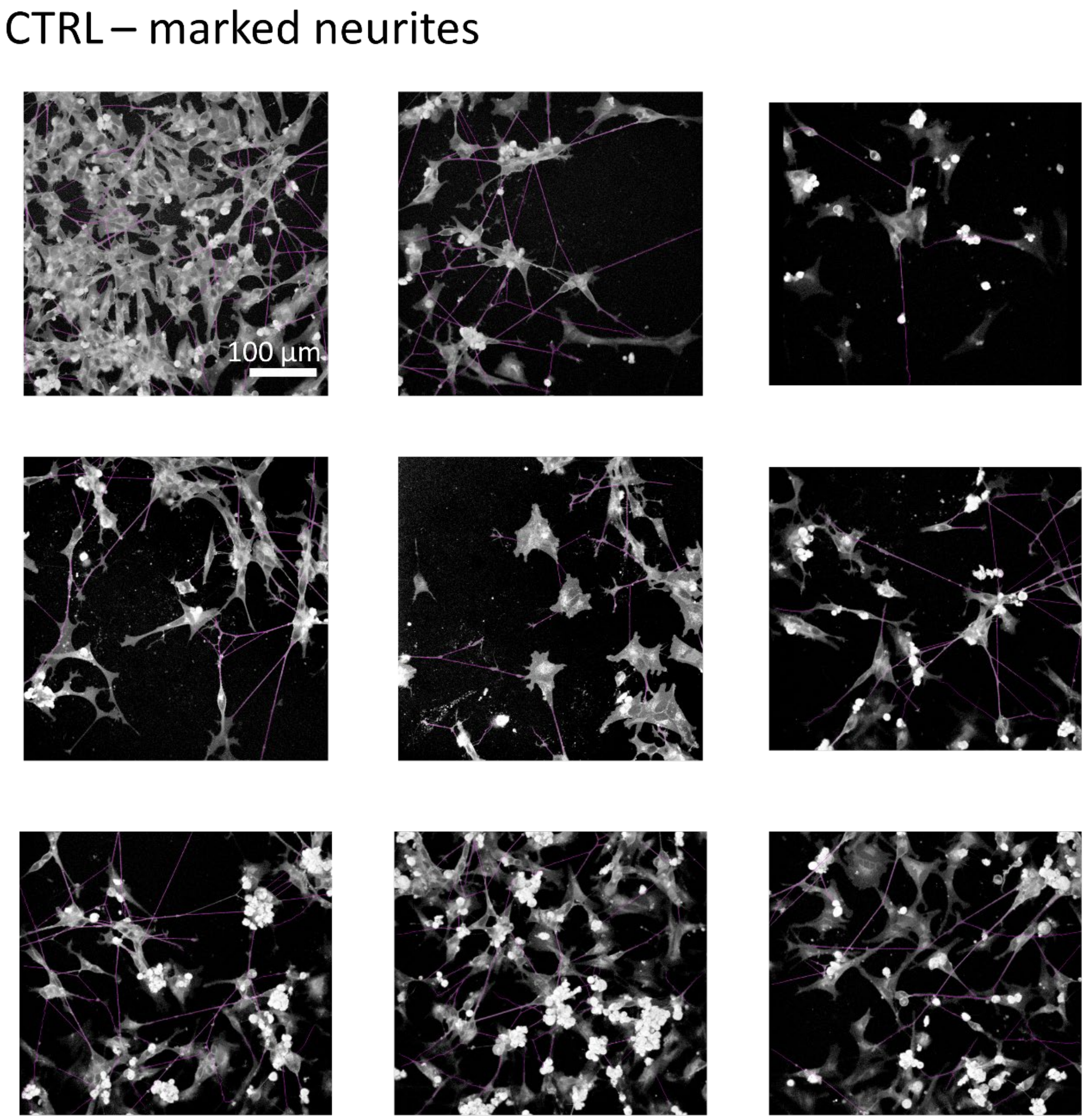

*Figure S 27: Images of unexposed control (CTRL) neurons used for the analysis of neurite length (violet) after 100h of incubation.*

#### $TiO_2$ – marked neurites

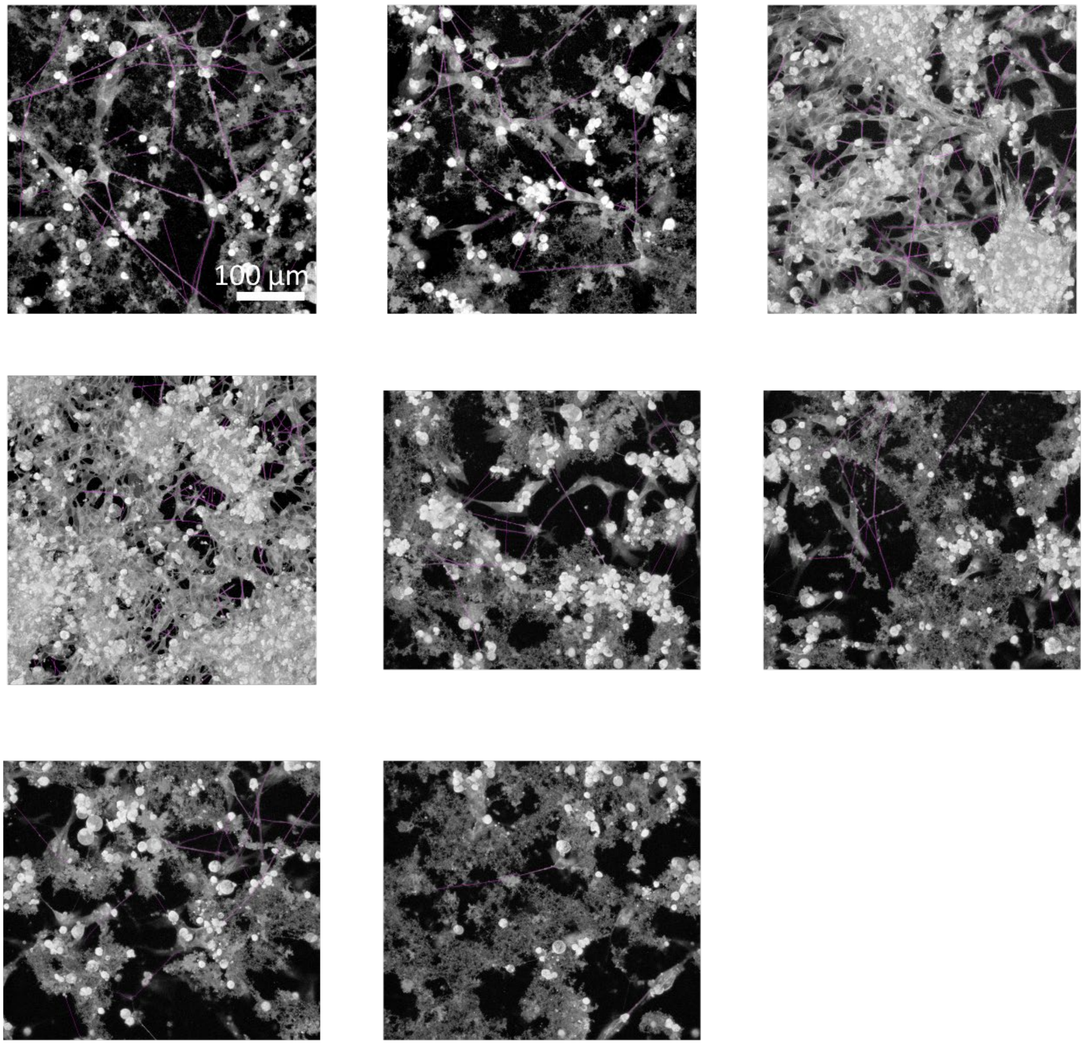

*Figure S 28: Images of $TiO_2$ nanotube exposed neurons used for the analysis of neurite length (violet) after 100h of incubation.*

### $CeO_2$ – marked neurites

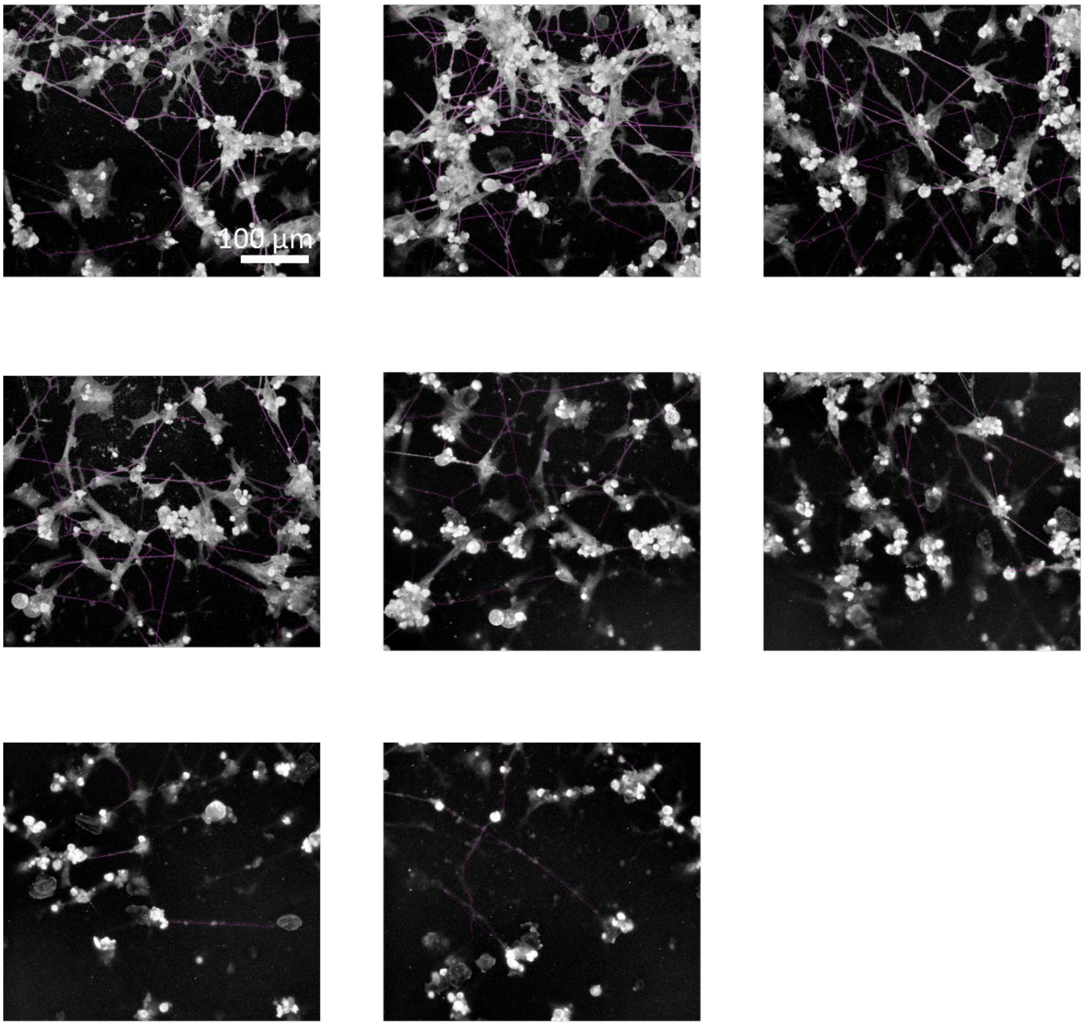

*Figure S 29: Images of $CeO_2$ exposed neurons used for the analysis of neurite length (violet) after 100h of incubation.*

### Diesel – marked neurites

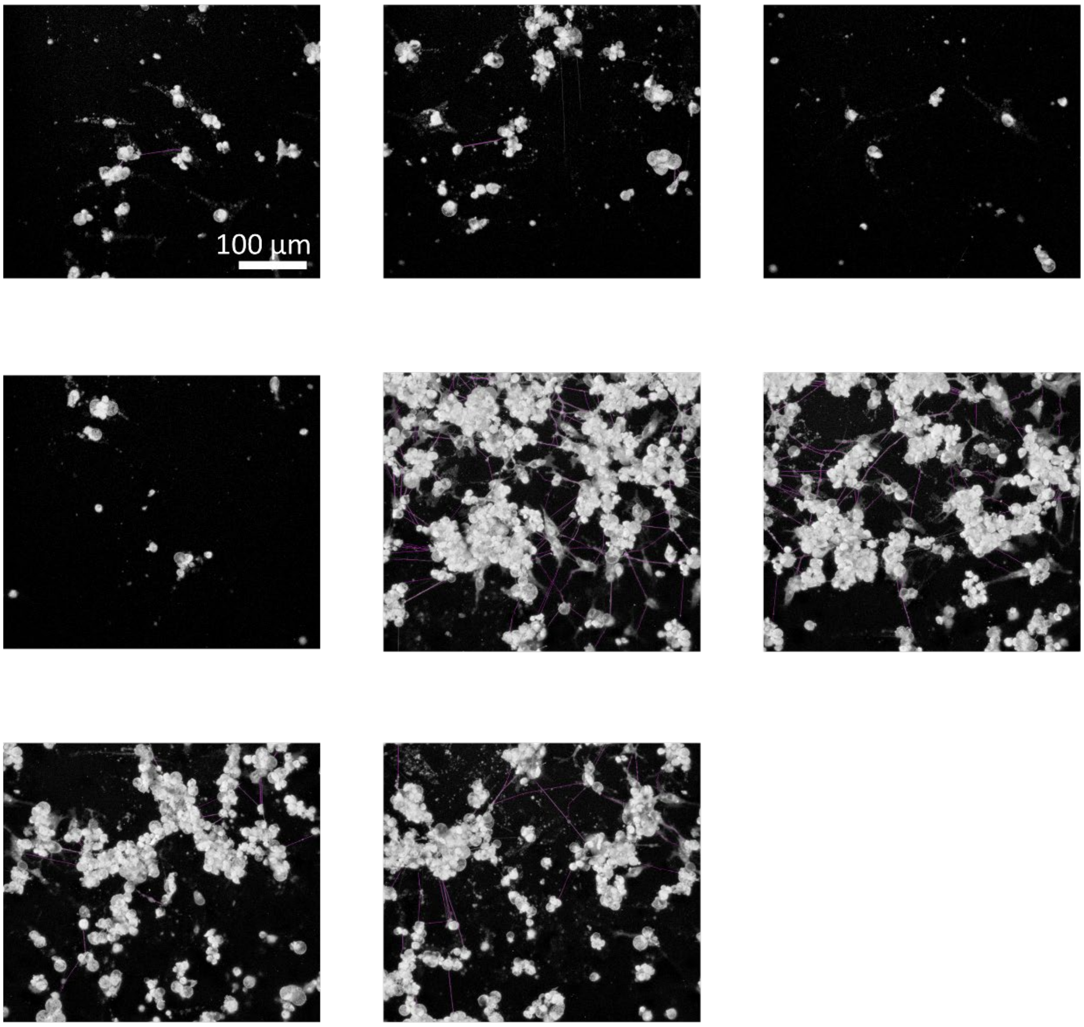

*Figure S 30: Images of Diesel exposed neurons used for the analysis of neurite length (violet) after 100h of incubation.*

maghemite γ-$Fe_2O_3$ - marked neurites

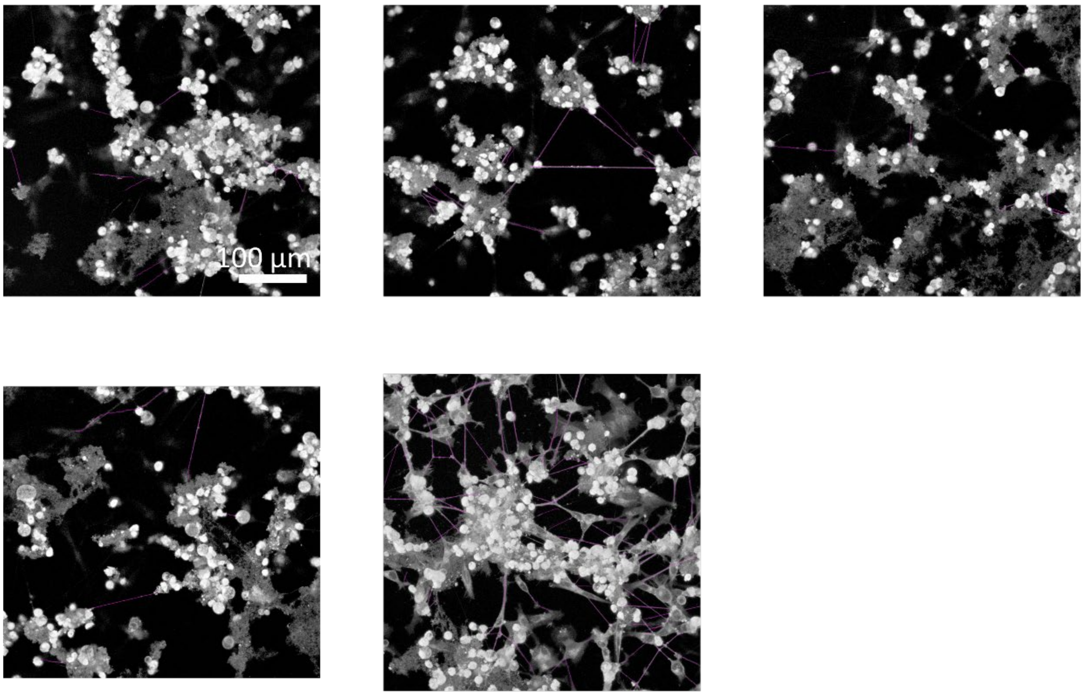

*Figure S 31: Images of γ-$Fe_2O_3$ exposed neurons used for the analysis of neurite length (violet) after 100h of incubation.*

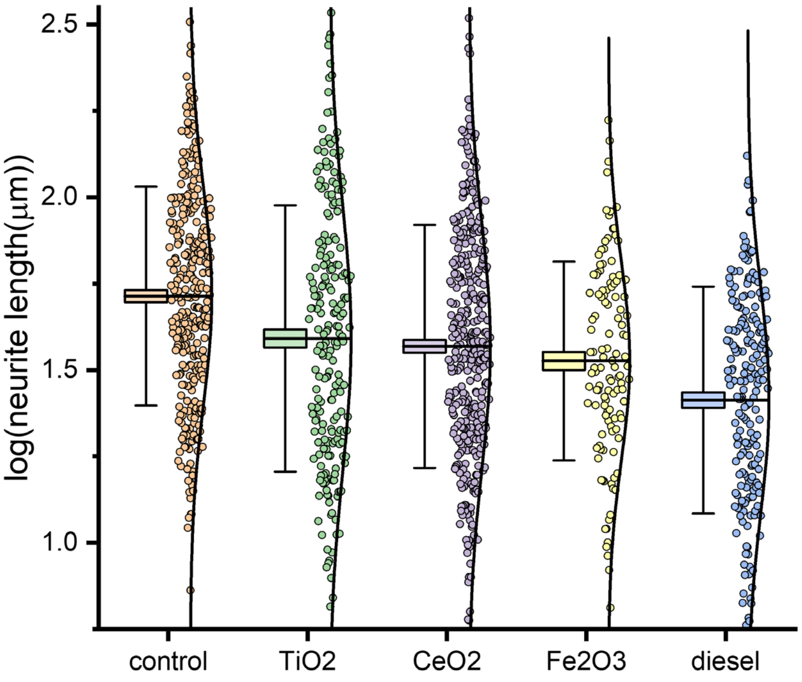

*Figure S 32: Distribution of log-transformed neurite lengths of neurons in control, $TiO_2$ nanotube, $CeO_2$, γ-$Fe_2O_3$, and diesel exhaust particles exposed groups.*

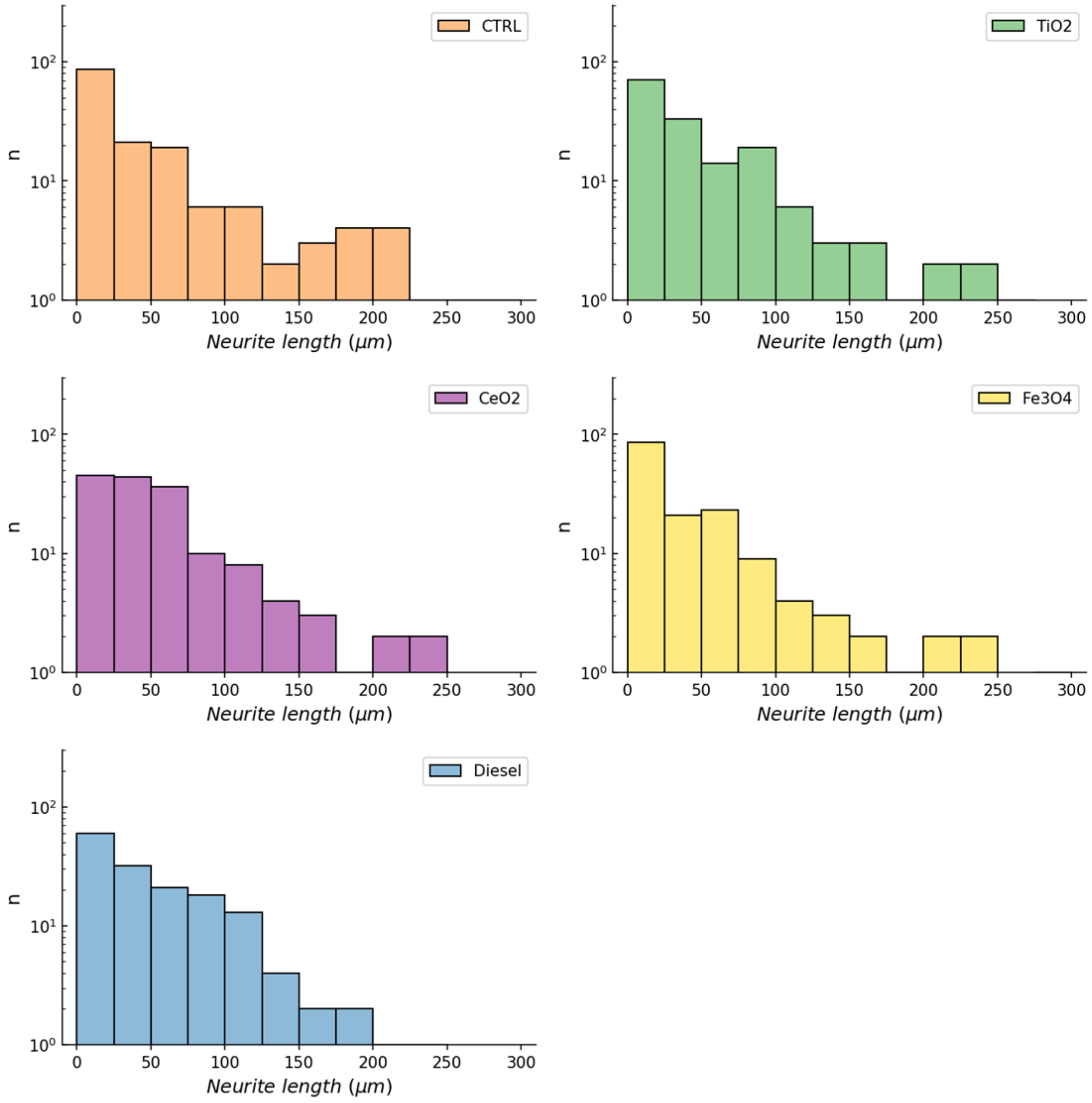

*Figure S 33. Histograms of neurite lengths of control, $TiO_2$, $CeO_2$, $Fe_2O_3$, and diesel exhaust particles of 1 hours exposed neurons. Data distribution for all groups is not normal as indicated by the histogram shapes. The normality of the data was further assessed using the D'Agostino and Pearson's Test and Shapiro-Wilk. Test p-values, p-normal and p-saphiro, respectively, < 0.05 indicate a deviation from normality.*

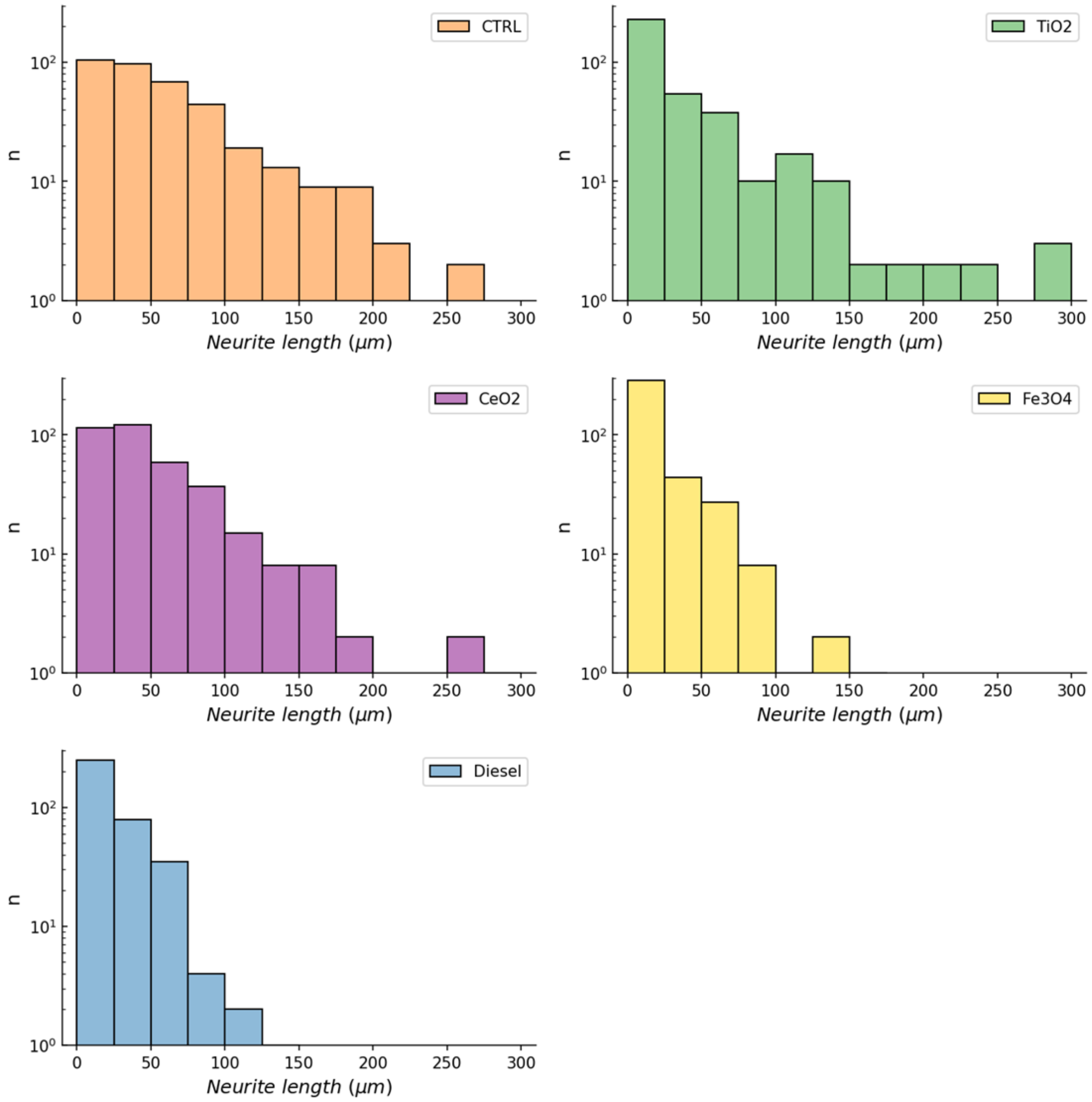

*Figure S 34: Histograms of neurite lengths of control, $TiO_2$, $CeO_2$, $Fe_2O_3$, and diesel exhaust particles of 100 hours exposed neurons. Data distribution for all groups is not normal as indicated by the histogram shapes. The normality of the data was further assessed using the D'Agostino and Pearson's Test and Shapiro-Wilk. Test p-values, p-normal and p-saphiro, respectively, < 0.05 indicate a deviation from normality.*

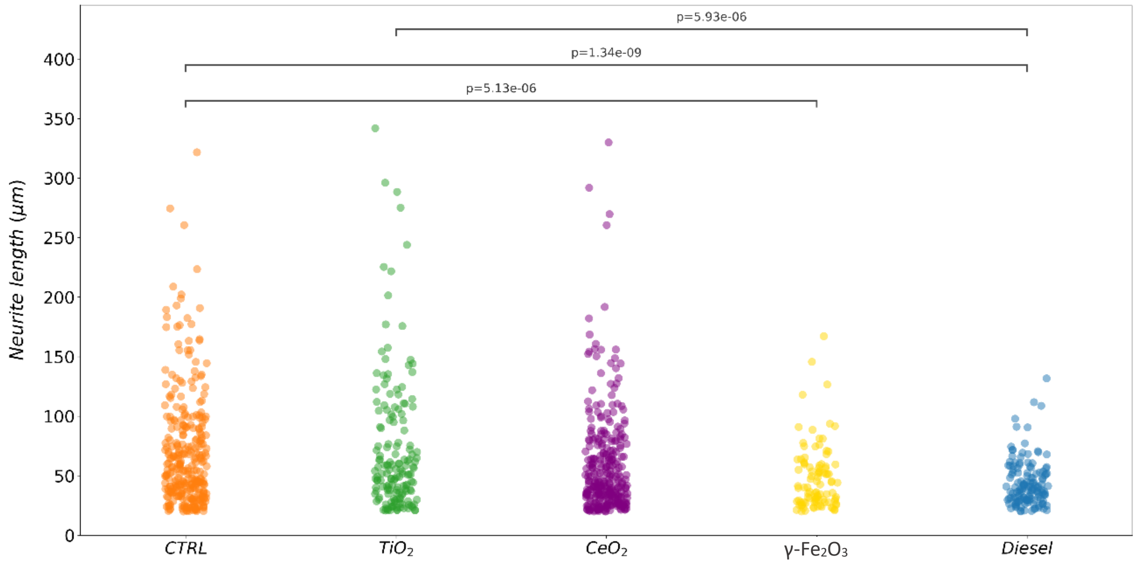

*Figure S 35: Mann–Whitney U test was performed on the neurite lengths of control, $TiO_2$, $CeO_2$, $Fe_2O_3$, and diesel exposed neurons due to the non-normal distribution of data. Significant difference occurred between the neurite lengths of control and $Fe_2O_3$ and Diesel exposed neurons and between $TiO_2$ and Diesel exhaust particles exposed neurons.*

### 6 Main article FIGURE 4 supplementary material

The samples were exposed to $TiO_2$, γ-$Fe_2O_3$, $CeO_2$, and diesel exhaust nanoparticles and incubated for 100 hours prior to microscopy. Images were acquired with the 20x magnification objective and 0.8 numeric aperture lens. The excitation/emmision combinations of lasers and detectors used are laid out in

Before differentiation, we seeded approximately 1 x $10^6$ of neurons. Considering that the average mass of a human cell is 3-4 ng (DOI: 10.1039/b615235j), we estimate that we had around 3 mg of cells before exposure to nanomaterials.

Volume of nanomaterials added to cells was 20 µl for all samples. Final mass of the material added to cells and the mass-to-mass ratio of nanomaterial to cell is given in Table S 6.

*Table S 6. Mass concentration of nanomaterials used for differentiated neurons exposure.*

| *Material* | *Mass of added material (mg)* | $\frac{mNM}{mCELLS}$ $({}^{mg}/_{g})$ |
|---|---|---|
| $TiO_2$ nanotubes | 0.013 | 4 |
| γ-$Fe_2O_3$ | 0.036 | 12 |
| $CeO_2$ | 0.032 | 11 |
| Diesel (DEP9.7) | 0.013 | 3 |

*Table S 7. Experimental setup for images in Figure 4.*

| *Label* | *$\lambda_{EXCITATION}$ (nm)* | *Laser power (%)* | *$\lambda_{EMMISION}$ (nm)* |
|---|---|---|---|
| CMDR | 640 | 1 | 650-700 |
| anti-β-Amyloid Ab | 488 | 10 | 650-700 |
| Backscattering | 488 | 1.3 | 488 |

Pixel size was 400 nm, pinhole 20 µm (1.01 AU @650 nm), and pixel dwell time 4 µs with 1 line accumulation. 3D stacks were acquired with 4 µm increments for a total of 20 µm in hight. Images presented in Figure4A-E are maximum intensity projections.

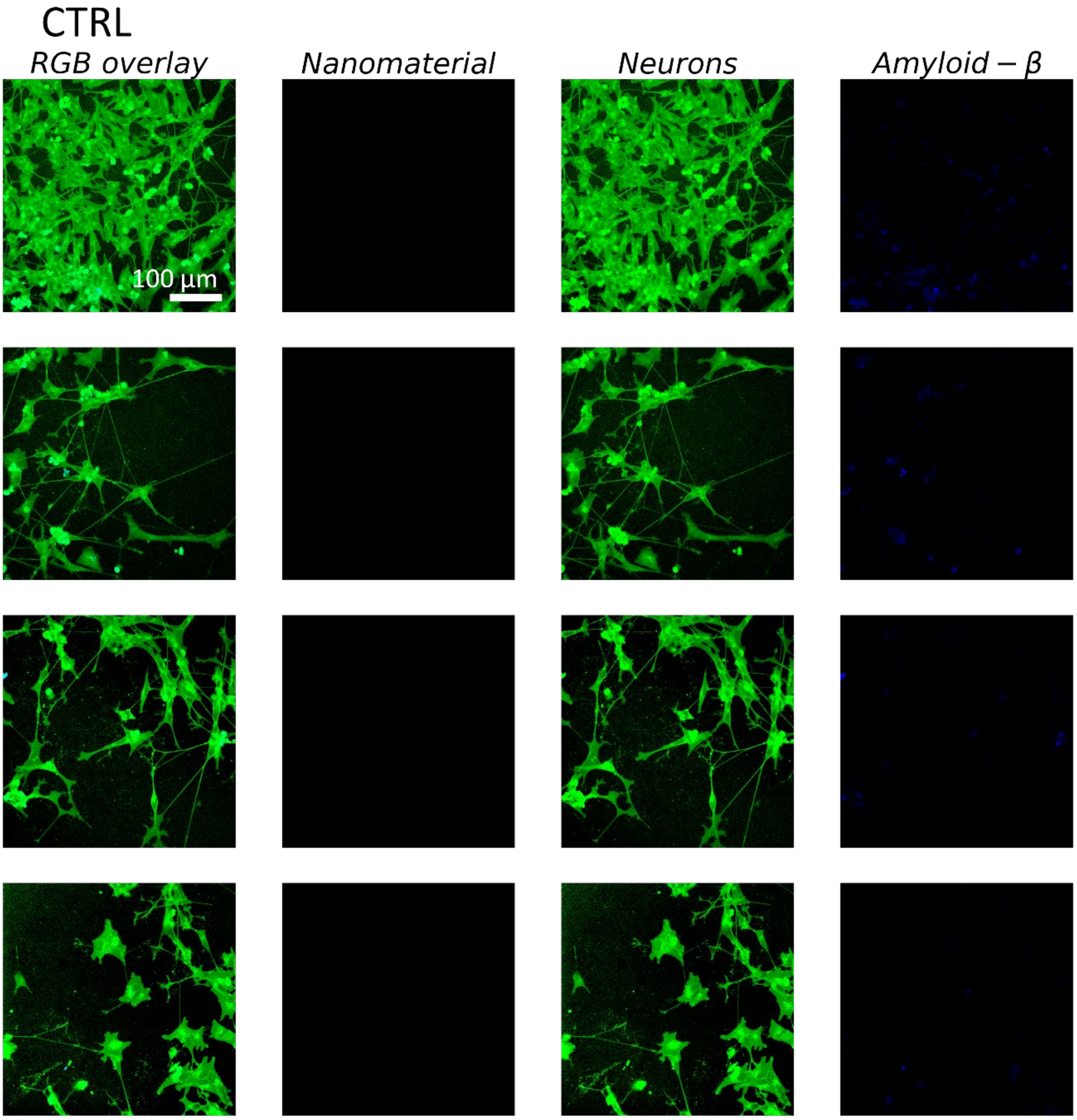

*Figure S 36: Micrographs of unexposed control neurons for neurite length measurement. RGB images with plasma membrane label in green (CellMask Orange - CMO) and anti-Aβ antibody in blue are in the first column. The remaining columns display decomposed channels depicting neurons (CMO) and Aβ signal, respectively.*

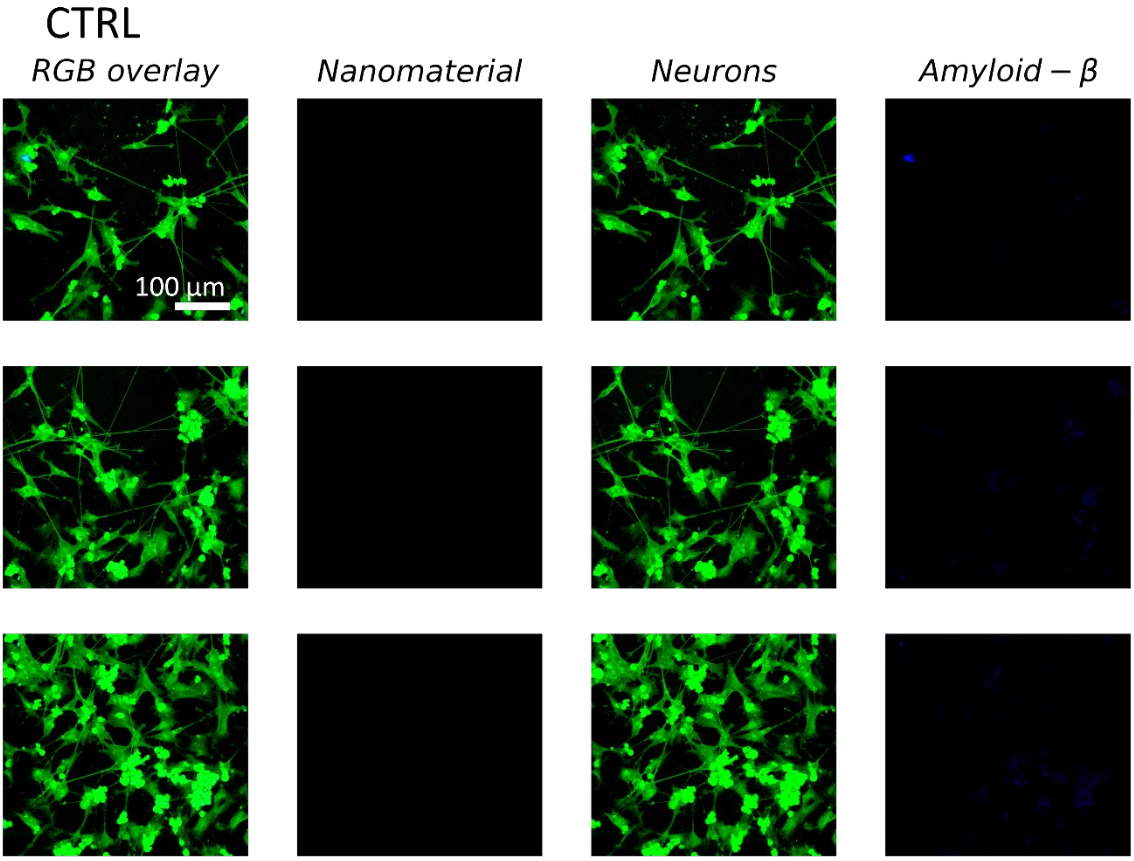

*Figure S 37: Micrographs of unexposed control neurons for neurite length measurement. RGB images with plasma membrane label in green (CellMask Orange - CMO) and anti-Aβ antibody in blue are in the first column. The remaining columns display decomposed channels depicting neurons (CMO) and Aβ signal, respectively.*

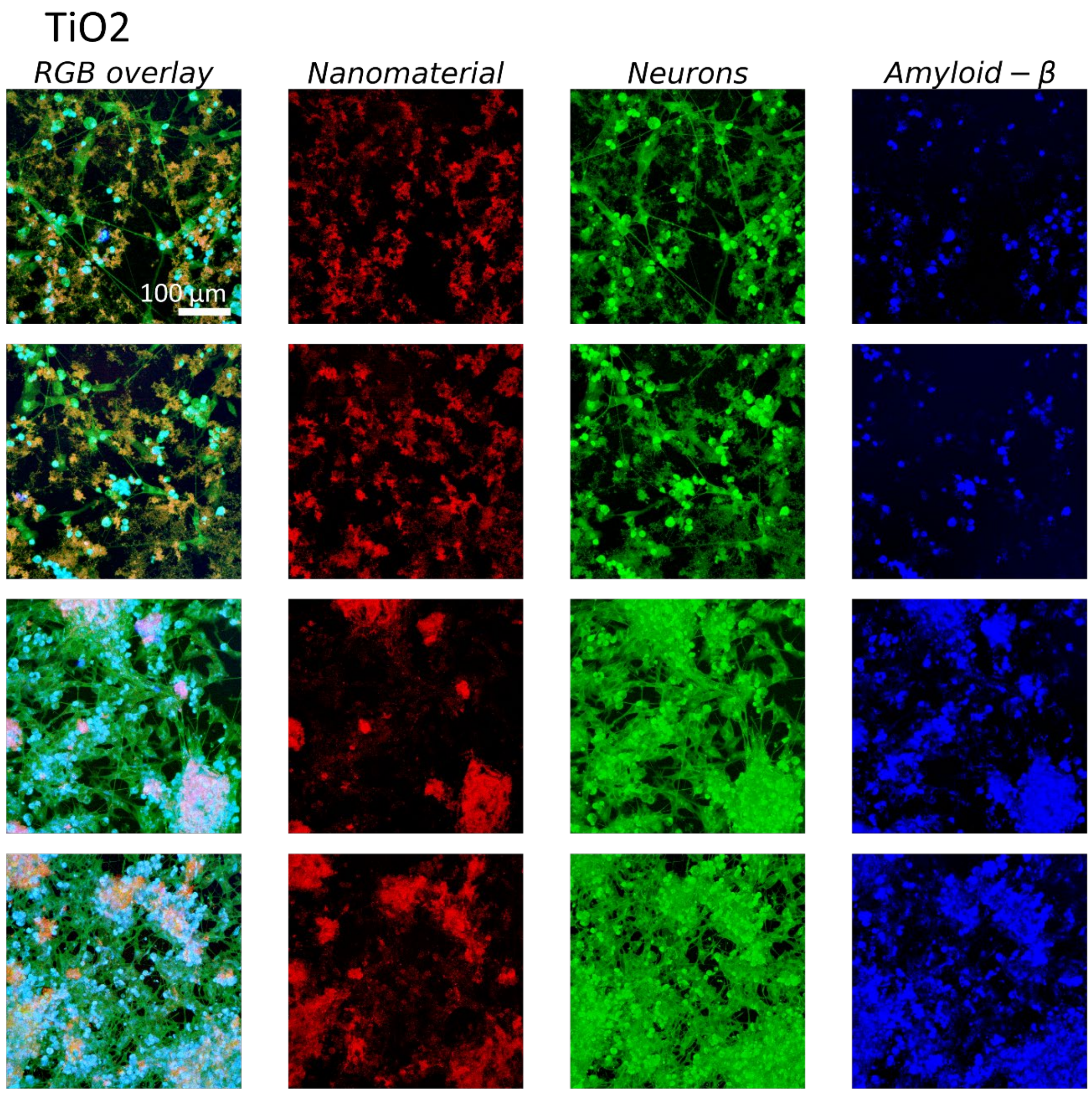

*Figure S 38: Micrographs of neurons exposed to $TiO_2$ nanotubes for neurite length measurement. RGB images with plasma membrane label in green (CellMask Orange - CMO) and anti-Aβ antibody in blue are in the first column. The remaining columns display decomposed channels depicting neurons (CMO), Aβ signal, and scattering signal, respectively.*

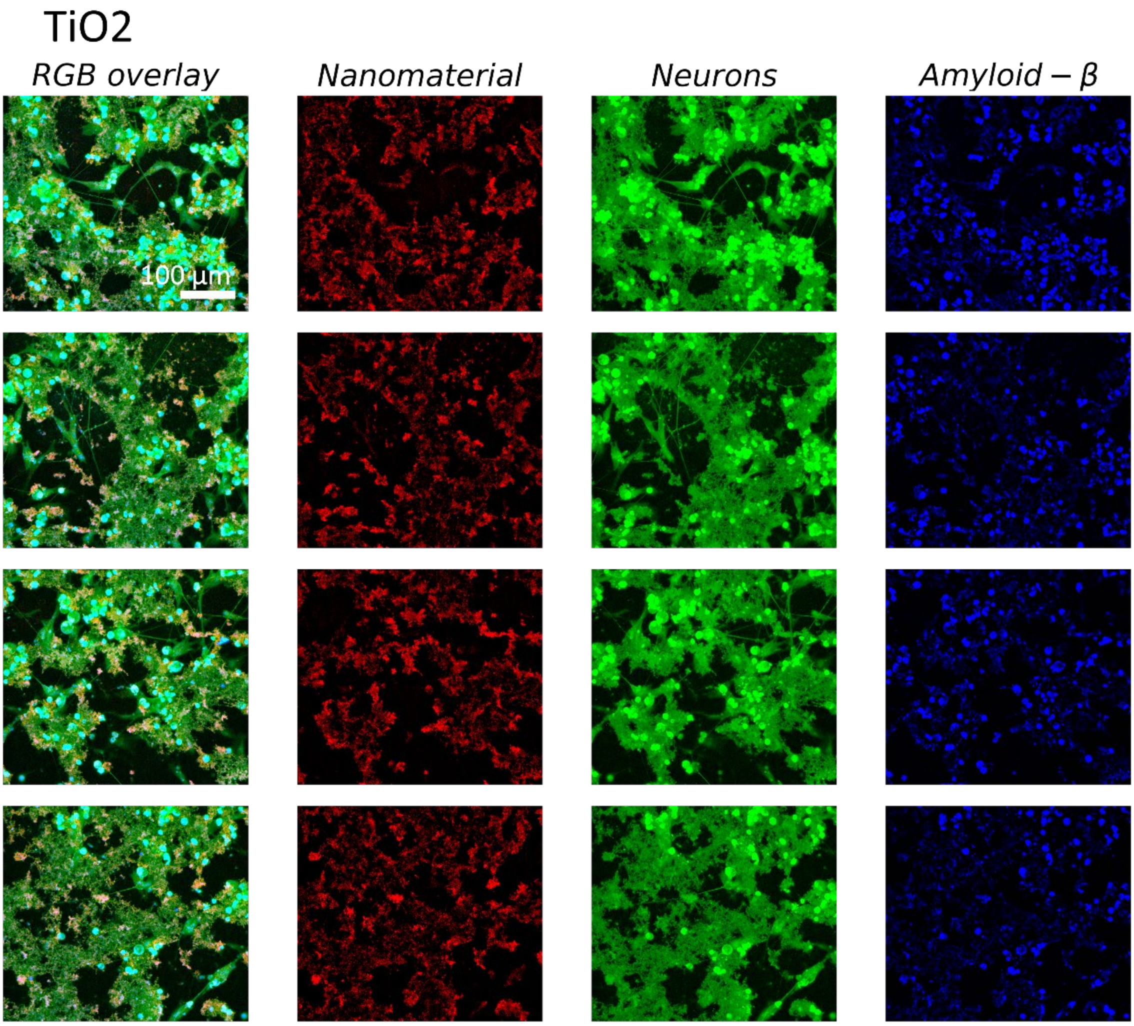

*Figure S 39: Micrographs of neurons exposed to $TiO_2$ nanotubes for neurite length measurement. RGB images with plasma membrane label in green (CellMask Orange - CMO) and anti-Aβ antibody in blue are in the first column. The remaining columns display decomposed channels depicting neurons (CMO), Aβ signal, and scattering signal, respectively.*

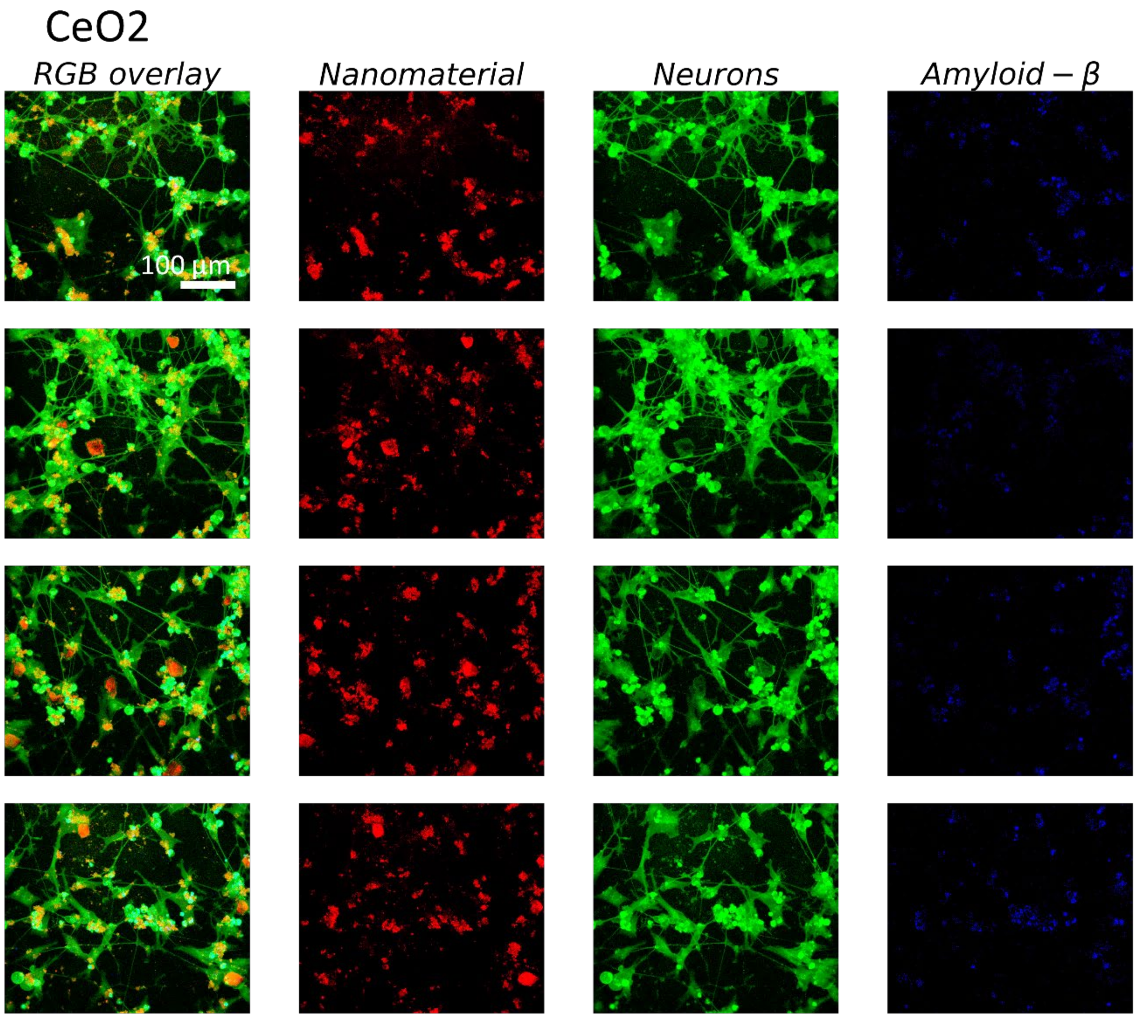

*Figure S 40: Micrographs of neurons exposed to $CeO_2$ for neurite length measurement. RGB images with plasma membrane label in green (CellMask Orange - CMO) and anti-Aβ antibody in blue are in the first column. The remaining columns display decomposed channels depicting neurons (CMO), Aβ signal, and scattering signal, respectively.*

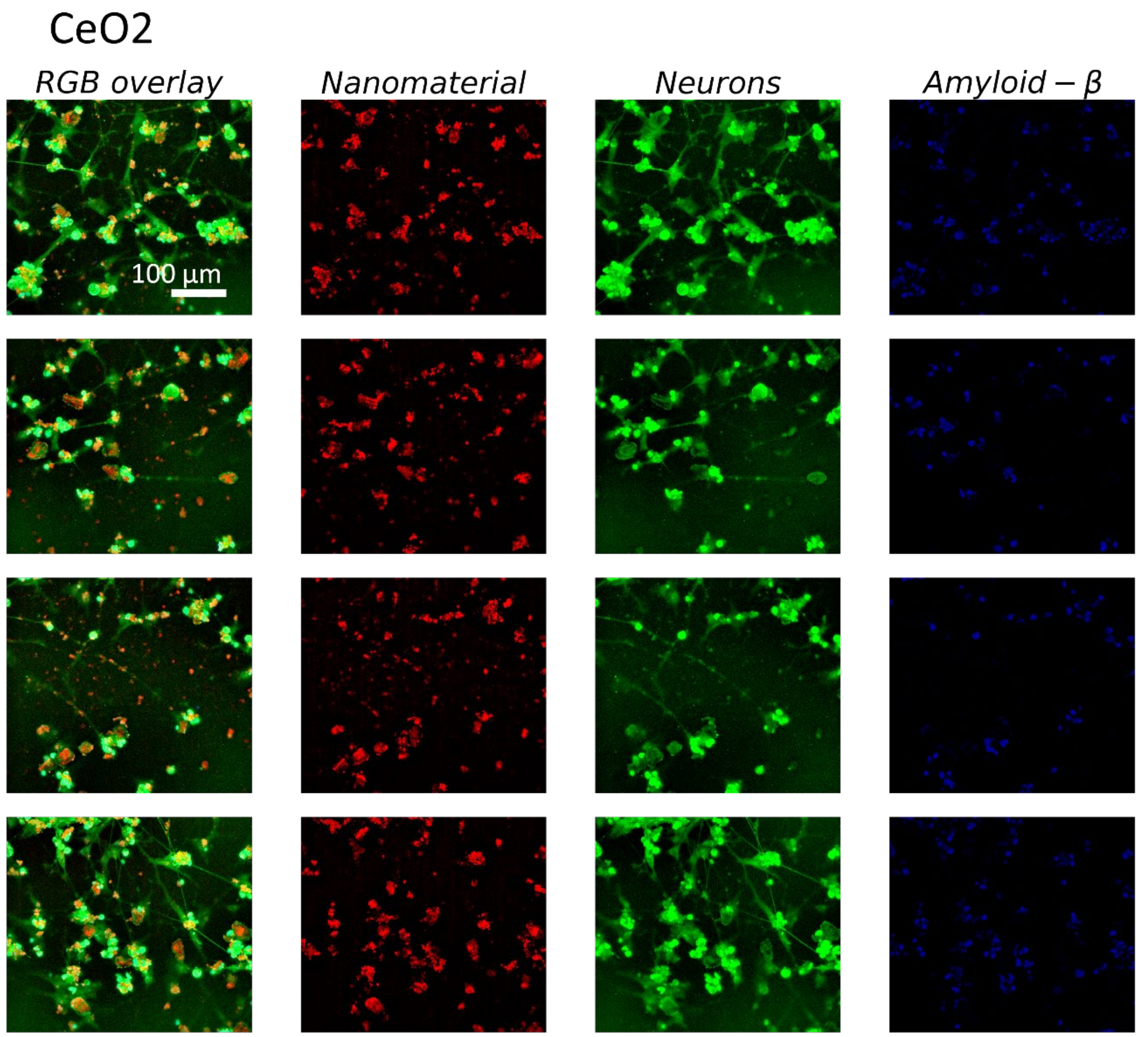

*Figure S 41: Micrographs of neurons exposed to $CeO_2$ for neurite length measurement. RGB images with plasma membrane label in green (CellMask Orange - CMO) and anti-Aβ antibody in blue are in the first column. The remaining columns display decomposed channels depicting neurons (CMO), Aβ signal, and scattering signal, respectively.*

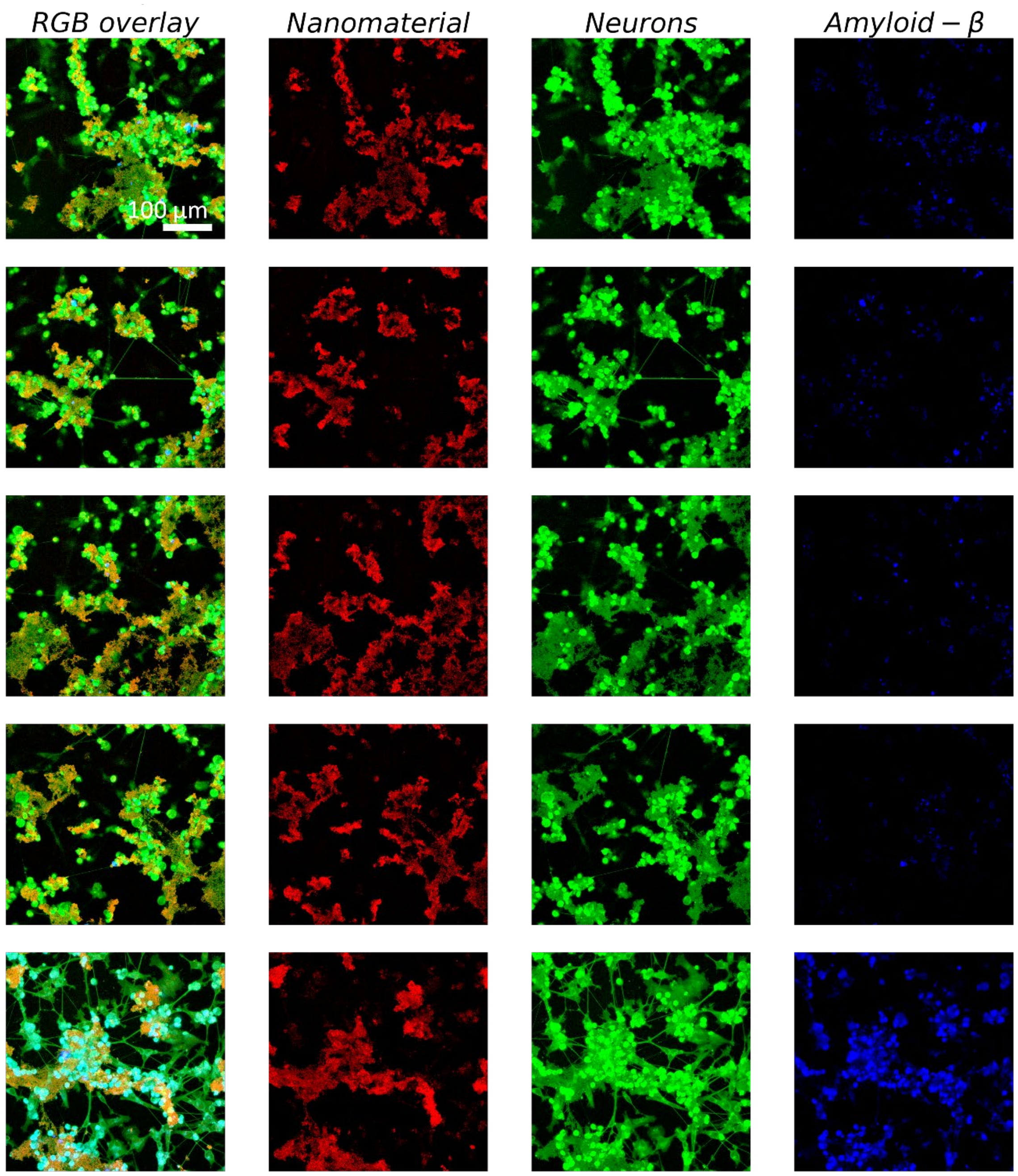

*Figure S 42: Micrographs of neurons exposed to γ-$Fe_2O_3$ for neurite length measurement. RGB images with plasma membrane label in green (CellMask Orange - CMO) and anti-Aβ antibody in blue are in the first column. The remaining columns display decomposed channels depicting neurons (CMO), Aβ signal, and scattering signal, respectively.*

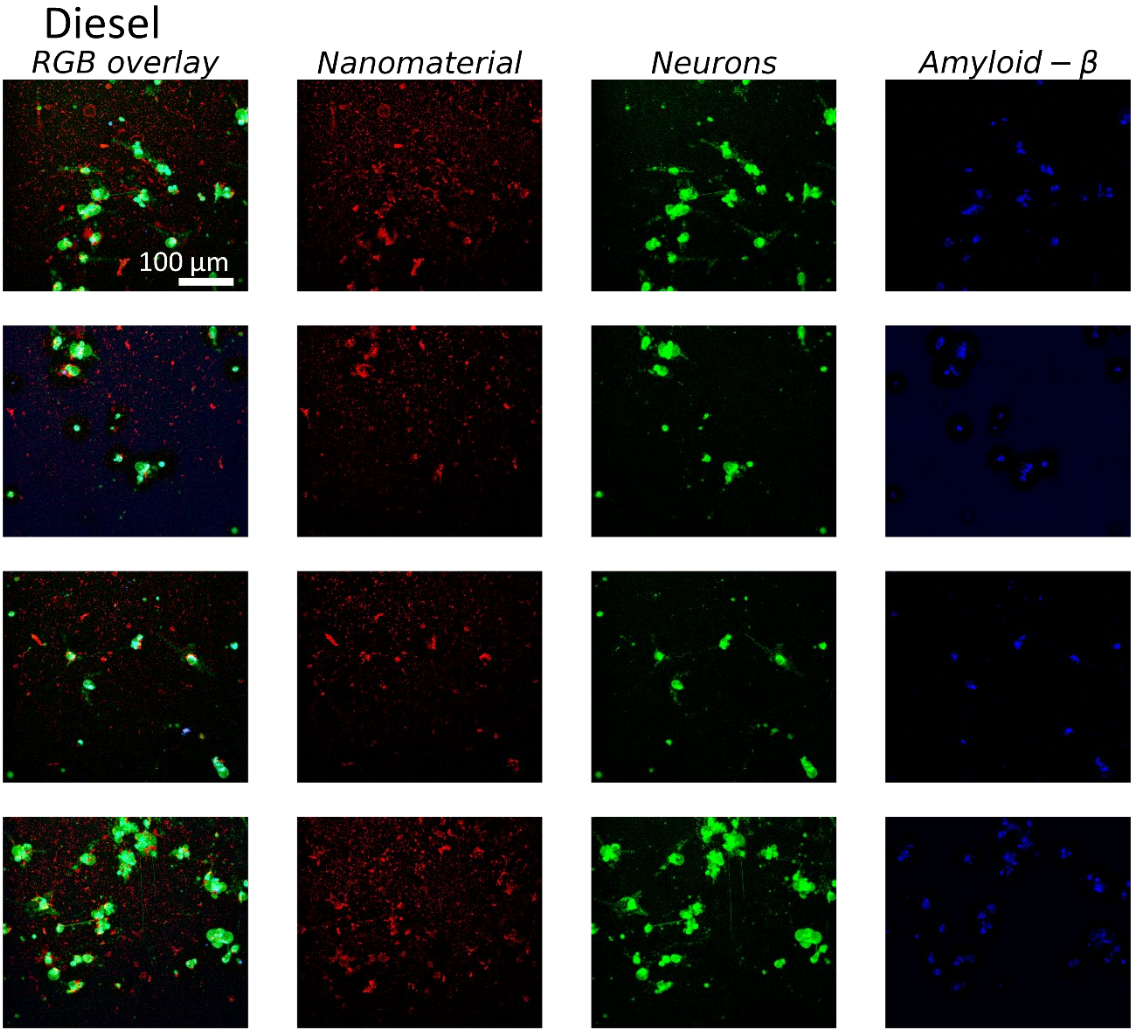

*Figure S 43: Micrographs of neurons exposed to Diesel exhaust nanomaterials for neurite length measurement. RGB images with plasma membrane label in green (CellMask Orange - CMO) and anti-Aβ antibody in blue are in the first column. The remaining columns display decomposed channels depicting neurons (CMO), Aβ signal, and scattering signal, respectively.*

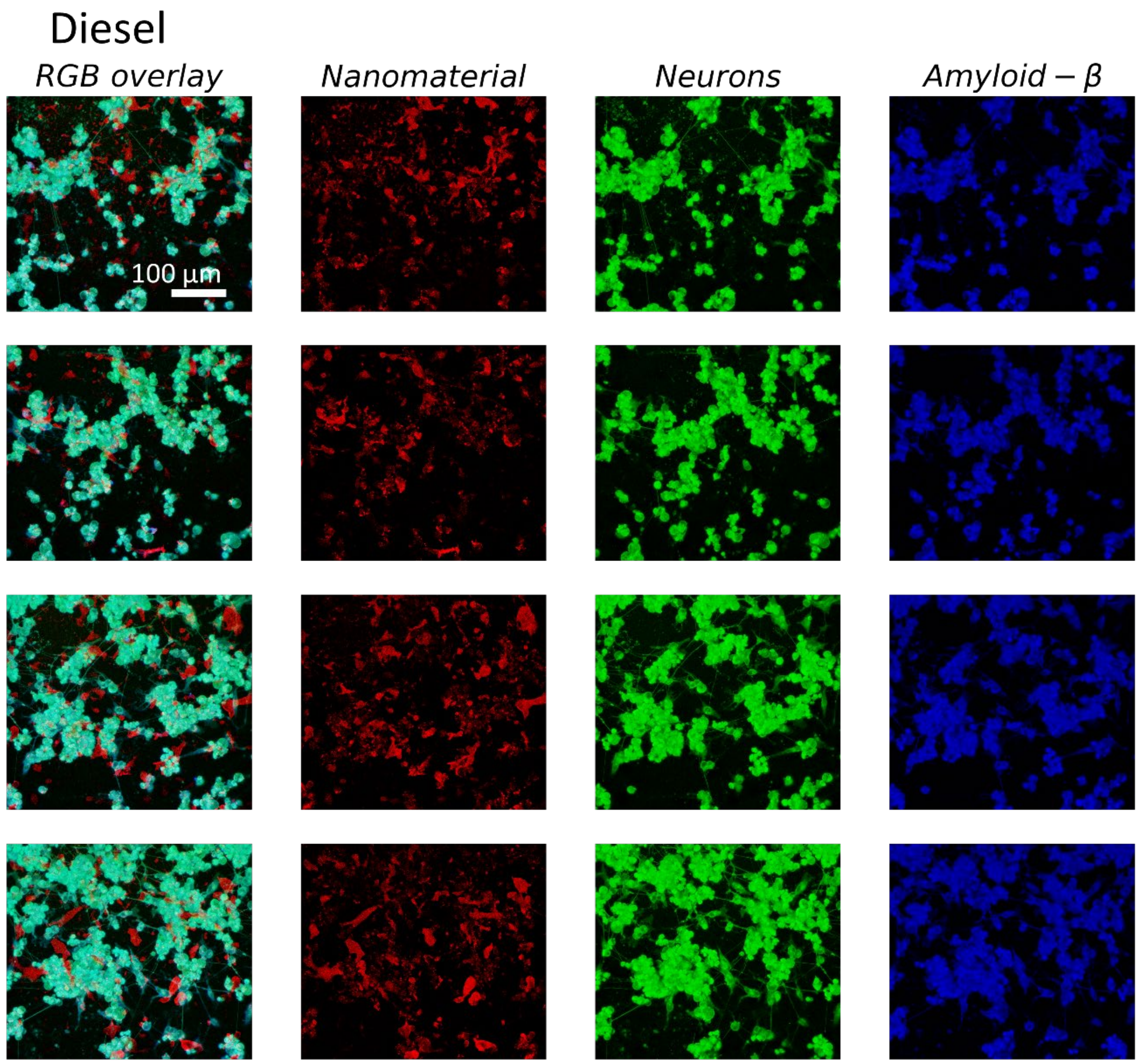

*Figure S 44: Micrographs of neurons exposed to Diesel exhaust nanomaterials for neurite length measurement. RGB images with plasma membrane label in green (CellMask Orange - CMO) and anti-Aβ antibody in blue are in the first column. The remaining columns display decomposed channels depicting neurons (CMO), Aβ signal, and scattering signal, respectively.*

For the analysis involving the intensity of Aβ and the proportion of Aβ signal colocalized with nanomaterial, Python scripts were utilized. The algorithm outlining the steps of this analysis is shown in the image below.

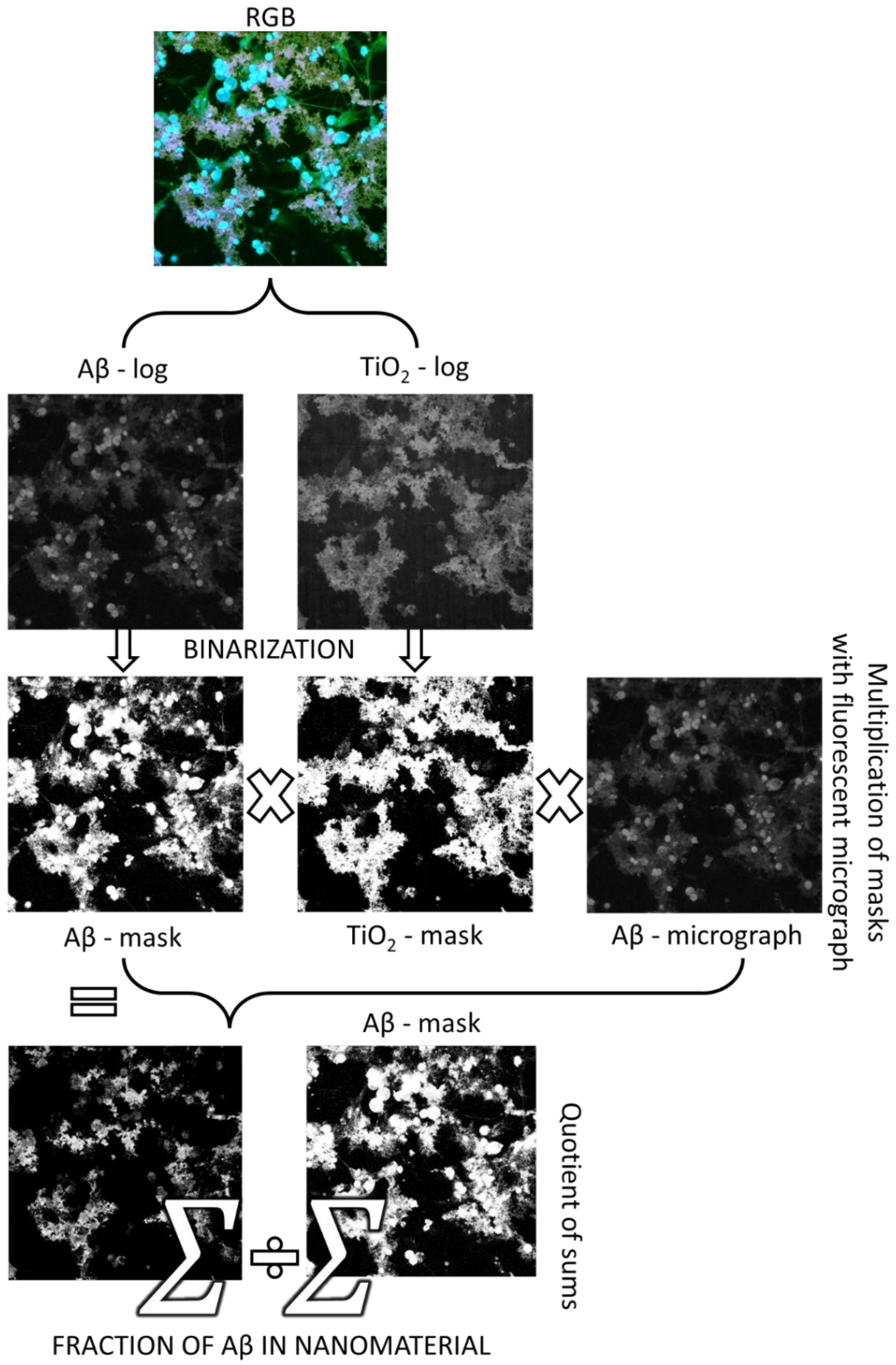

*Figure S 45: Aβ and total nanomaterial binary masks for Aβ intensity and Aβ fraction in nanomaterial calculation. Initially, RGB micrographs were decomposed into individual channels, which were then subjected to a logarithmic transformation. A Fast Fourier Transform bandpass filter was applied to eliminate structures smaller than 2 pixels and larger than 800 pixels in diameter for both the nanomaterial and AB channels. This effectively remove all objects smaller than 400 nm in one dimension (pixel size = 200 nm). Subsequently, the images were binarized to generate masks. To determine the density of Aβ plotted in Figure 4 of the main paper, we used the same algorithm as explained in Figure S 21. To assess the fraction of Aβ present in nanomaterial aggregates, the Aβ mask was multiplied with the nanomaterial mask, resulting in the co-localization of both masks. The surface area of this product divided by the surface area of the Aβ mask yields the fraction of Aβ in nanomaterials.*

### 7 Dose relevance

In order to determine whether the dosage we administered are relevant for human exposure we estimated the dose that might accumulate in the brain by reviewing the literature for the amount of each nanoparticle found in the brain of Alzheimer's disease patients and found that in the case of diesel exhaust and iron oxide the concentrations we used are not far from the amounts found in the brain.

$TiO_2$ nanotubes are an engineered nanomaterial relevant to occupational risk. $CeO_2$, iron oxide (γ-$Fe_2O_3$), and diesel exhaust are examples of particulate matter found in polluted air and are relevant as a public health risk. We selected $CeO_2$ due to its predominantly nontoxic nature, whereas iron oxide and diesel exhaust (DEP9.7) nanoparticles are linked to neurodegenerative diseases by epidemiological studies. Iron oxide nanoparticles were found in the brains of Alzheimer's disease patients living in polluted areas, such as Mexico City.

**$TiO_2$**, as a white pigment, is used as a food additive in personal care products (e.g. toothpaste) and in many other consumer products. It contains a fraction of nanosized primary particles (<100 nm) (Peters et al., 2014, 2020). Titanium can be found in human tissue at lower average concentrations ranging from 1 µg/g (Peters et al., 2020) up to 80 µg/g (Baj et al., 2023) in the human brain. We used a very high concentration of 3 mg/g, which resulted in extracellular amyloid beta plaque formation, yet the lengths of neurites didn't decrease as much as in the case of diesel exhaust nanoparticles, despite the high concentration used. Several in vivo studies reviewed by Song et al. (Song et al., 2015) have demonstrated that the $TiO_2$ NPs can be transported and accumulated in the brain, but mostly in intranasal instillation or inhalation studies, eventually leading to CNS dysfunctions. Titanium content in mouse hippocampus after 90 days of continuous intranasal exposure to $TiO_2$ nanoparticles was 0.6 mg/g tissue at the highest administered dose of 10 mg/kg body weight (Ze et al., 2014), which is also much higher than the levels observed in humans. Thus, neuroinflammation and impairment of spatial memory observed in mice and our in vitro experiment occur at more than 10 times higher brain content than it is found in humans. The effect of $TiO_2$ nanoparticles on the brain is thus still controversial since detrimental effects are observed at very high doses, which are not observed in human tissues.

On the other hand, we estimated the relevance of our exposure levels also by considering The National Institute for Occupational Safety and Health recommends occupational exposure limits of 2.4 $mg/m^3$ for fine $TiO_2$ and 0.3 $mg/m^3$ for ultrafine (including engineered nanoscale such as $TiO_2$ nanotubes we used in this manuscript). Assuming breathing rate of 15L/h, 40-hour week and body mass 70kg, one would have to breath about 3000 years to obtain the dose 3 mg/g body mass, where we neglected that exposed dose might not distribute equally over entire body mass but might accumulate preferentially in the brain during nasal breathing on one hand, and that not all of the nanoparticles are retained in the body on the other hand. Since the $TiO_2$ nanotubes didn't induce shortening of the neurites event at such high levels used in this manuscript we can conclude that $TiO_2$ nanotubes probably don't induce neurodegeneration and that the controversy regarding the neurotoxicity of $TiO_2$ observed in animals might arise from the high levels used in animal studies.

**$CeO_2$** nanoparticles are used as fuel additive in motor vehicles to reduce carbon monoxide, nitrogen oxides, and hydrocarbons in exhaust gases, thus the particles are released in the air.
Effects of $CeO_2$ nanoparticles are contradictory, demonstrating all kinds of effects, from protective to toxic (Gagnon & Fromm, 2015). For example $CeO_2$ nanospheres and nanorods improved cognitive impairment following mild traumatic brain injury in VC57BL/6J male mice at the dose of 0.02 mg/g body weight, although the concentration in the brain was not reported (Fiorani et al., 2015; Gagnon & Fromm, 2015).

The mean magnetite **iron** oxide ($Fe_2O_3$) concentration of several millimeters large, chemically unfixed post-mortem tissue from Alzheimer's cases can reach up to 400 µg/g (Finnegan et al., 2019). However, brain iron oxide (magnetite) airborne particles can be highly concentrated in sub-micrometer regions in the human brain (Maher et al., 2016). Such magnetite nanospheres are ubiquitous and abundant in airborne particulate matter pollution. They are $<\sim$200 nm in diameter and can enter the brain directly via the olfactory bulb (Maher et al., 2016). Examination of the human frontal cortex brain samples obtained from subjects who lived in Mexico City revealed that magnetite nanoparticles have an external, rather than an endogenous source (Maher et al., 2016). Although the highest brain magnetite concentration was 10 µg/g dry tissue, the transmission electron micrographs of brain thin sections showed that the nanoparticles are located in about 1% of the total area. Considering that some regions don't contain nanoparticles, we can assume that the local concentration of nanoparticles can be up to 100 times higher as compared to larger brain regions. Therefore, we can safely assume that relevant in vitro cell line exposure is about 1 mg/g, which about ten times less (12 mg/g) the levels we used. Thus, we believe that the toxic effect of the iron nanoparticles in the in vitro cell line system we used might not be relevant to human exposure.

We couldn't find the levels of **diesel exhaust** particles in brain tissue since they are mainly composed of carbon and probably can't be differentiated from the tissue. However, iron is present in most of diesel exhaust samples, the iron content can vary from 8 weight % to 55 weight %, being on average about 20 weight % (Viskup et al., 2020). Thus if we assume that the iron accumulated in samples of Alzheimer's cases which can reach up to 400 µg/g (Finnegan et al., 2019) originates from diesel exhaust particles then the brain burden of diesel exhaust particles might reach up to 2 mg/g tissue weight. We used 3 mg/g, thus the neurite degeneration and apoptosis of neurons we observed in cell culture might be relevant to diesel exhaust exposure.
